# Disclosing temperature sensitivity of West Nile virus transmission: novel computational approaches to mosquito-pathogen trait responses

**DOI:** 10.1101/2024.09.16.613097

**Authors:** Julian Heidecke, Jonas Wallin, Peter Fransson, Pratik Singh, Henrik Sjödin, Pascale Stiles, Marina Treskova, Joacim Rocklöv

## Abstract

Temperature influences the transmission of mosquito-borne pathogens with significant implications for disease risk in the context of climate change. Mathematical models of mosquito-borne infections rely on functions that capture mosquito-pathogen interactions in response to temperature to better estimate transmission dynamics. To derive these functions, experimental studies provide valuable data on the temperature sensitivity of mosquito life-history traits and pathogen transmission. However, the scarcity of experimental data and inconsistencies in methodologies for analysing and comparing temperature responses across mosquito species and pathogens present challenges to accurately modelling mosquito-borne infections. In this study, we investigated the thermal biology of West Nile virus (WNV), a major mosquito-borne infection. We critically reviewed existing experimental studies, obtaining temperature responses for eight mosquito-pathogen traits across 15 mosquito species. Using this data, we employed Bayesian hierarchical models to estimate temperature response functions for each trait and estimate their variation between species and experiments. We incorporate the resulting functions into mathematical models to estimate the temperature sensitivity of WNV transmission, focusing on six competent mosquito species of the genus *Culex*. Our study finds similarities in the temperature response among *Culex* species, with a general optimal transmission temperature around 24°C. We demonstrate that differing mechanistic assumptions in published models can result in a temperature optimum variation exceeding 3°C, underscoring the need for model scrutiny. Additionally, we identify substantial variability between trait temperature responses across experiments on the same mosquito species, indicating significant intra-species variation in trait performance. We provide recommendations for future experimental studies, emphasizing the critical need for additional data on all mosquito-pathogen traits, except for mosquito larva-to-adult development rate and survival, which are relatively well-documented. Incorporating additional data into our multi-species approach enhances the accuracy of temperature trait response estimates. These improvements are critical for forecasting future shifts in mosquito-borne disease risk due to climate change.

## Introduction

West Nile virus (WNV) is a pathogenic multi-host and multi-vector flavivirus. Transmission of the virus is maintained in an enzootic cycle between various wild birds and mosquitoes, primarily of the genus *Culex* (*Cx*.) [1,2]. Due to their opportunistic feeding behaviour, many *Cx*. mosquitoes can act as bridge vectors by transmitting the virus from its enzootic cycle to humans, equids, and other mammals [3–5]. The latter are dead-end hosts which cannot transmit the virus back to mosquitoes but can be infected. In humans, approximately 25% of infections progress to West Nile fever, while less than 1% result in a neuroinvasive disease with a case fatality ratio of 10% [1]. Several vaccines have been licensed for use in horses, but there are currently no vaccines nor therapeutic drugs available for use in humans [6,7]. Historically, WNV was associated with outbreaks across Africa, Eurasia, Australia, and the Middle East [8]. In 1999, WNV was introduced to the United States, initially in New York City. The virus found a conducive environment and quickly spread across the United States from coast to coast within four years. WNV has since been introduced in other countries across the Americas and can now be found almost globally [9]. WNV has also been extending its spatial range in Europe since the late 1990s, although much more slowly than in the United States. However, major outbreaks in 2010 and 2018 marked unprecedented large transmission events with significant geographical expansion to previously unaffected areas. For example, the 2018 outbreak in Europe was characterized by 2,083 locally reported human cases, a 7.2-fold increase compared to the previous season [10].

Climate change alters the global landscape in which WNV and other mosquito-borne diseases (MBDs) manifest, creating more suitable conditions for mosquitoes to transmit diseases in temperate regions [11]. In fact, the expansion and increasing frequency of outbreaks of WNV in Europe appears to be driven by climate change [12]. Overall, there is ample evidence that climate change has been and is projected to amplify WNV risk in several areas across the globe [11,13–17]. While climate change is only one of several drivers, along with land use, socio-economic conditions, and host mobility [17], suitable climatic conditions are necessary for the establishment and local transmission of any MBD [18]. Therefore, understanding climate-driven changes in WNV risk and having the tools to project such changes is essential for proactive adaptation to climate change.

Temperature is one of the key climatic drivers often limiting mosquito-borne disease risk in temperate regions [18,19]. Since mosquitoes are ectothermic organisms, temperature directly impacts their life-history traits such as lifespan, biting activity, and reproductive success. Furthermore, temperature impacts viral dynamics within mosquitoes, which affects the physical ability of mosquitoes to transmit viruses and the extrinsic incubation period [19]. An extensive body of laboratory-based studies reports reactions of mosquito-pathogen traits to changes in temperature. Mechanistic models of MBDs based on such data predict that transmission suitability responds to temperature in a nonlinear, unimodal fashion with optimal temperatures depending on the mosquito species and pathogen combination [19–23]. Recent predictions for WNV transmission hint towards optimal temperatures around 22-25°C [20,24–26].

Several studies have attempted to compare how the temperature suitability for WNV varies across *Cx*. species [15,20,26–28]. These comparisons are challenging due to the scarcity of laboratory experimental data to quantify point estimates and uncertainty for species-specific temperature response functions. In a previous study [20], this challenge was approached by partially pooling data between species. Empirical priors, derived from pooled experimental data from various other species in combination with subjectively chosen prior weightings, were employed to constrain the species-specific temperature response fits. Similar methods have been used in studies of other MBDs utilizing different prior datasets and weightings [22,29]. The subjective selection of prior datasets and weightings complicates objective comparisons of temperature responses across species. Here, we approach partial pooling of the temperature response fits more formally. We introduce a novel approach to analyse and compare the temperature response of mosquito-pathogen traits between species using Bayesian hierarchical models. We estimate hierarchical priors that represent the central tendency of the mosquito-pathogen trait temperature response and its variability between species and experiments within the species. In addition, we extend previous WNV temperature suitability analyses, by linking our transmission suitability models to mosquito population dynamics [20,25,26,28].

We update and expand a previously compiled dataset of laboratory experimental studies testing the temperature response of mosquito-pathogen traits focusing on mosquitoes acting as important disease vectors in temperate regions [20]. We ensure data integrity by thoroughly reviewing and applying conceptual changes to the original dataset. Based on this data, we derive temperature response fits for eight mosquito-pathogen traits across 15 mosquito species using the Bayesian hierarchical models. We investigate the generality of the temperature response of mosquito-pathogen traits between species, virus genotypes, and between experiments on the same species. Focusing on six *Cx*. mosquitoes (*Cx. pipiens, Cx. quinquefasciatus, Cx. pipiens molestus, Cx. pipiens pallens, Cx. restuans, Cx. tarsalis*) and their ability to transmit WNV, we build species-specific temperature-dependent WNV transmission suitability models. Lending towards a range of published models, we also demonstrate how mechanistic assumptions underlying transmission suitability models impact predictions of temperature optima for WNV transmission. Furthermore, we investigate how the common practice of ignoring statistical dependencies introduced by experiment identity (i.e., when integrating outcomes from multiple experiments on the same species) [20,22,23,25,29] biases temperature response estimates and uncertainty quantification. From our analysis, we derive recommendations for MBD thermal biology analyses and identify critical data needs where experimental data on traits between and within each species is scarce. Further temperature response laboratory experimental studies on these key traits are needed to strengthen the presented approach and enhance our understanding of climate change impacts on WNV and MBDs.

## Methods

We initially introduce the mathematical model for WNV transmission suitability using a trait-based approach. We then describe the process of deriving a species-specific parameterization of this model. First, we outline the data collection strategy that updates and extends a previously published dataset compiling outcomes of laboratory experiments measuring species’ trait performance at different temperatures [20]. We then introduce the functions that we fit to this data to describe the temperature response of each mosquito-pathogen trait. A description of the Bayesian hierarchical modelling approach and the Markov Chain Monte Carlo implementation to estimate the parameters of these models follows. The implementation of our analysis and its added value to previous analyses of MBD thermal biology of WNV [20,25,26] and other pathogens [19,22,23,29–32] are summarized in Figure 1.

**Figure 1.**
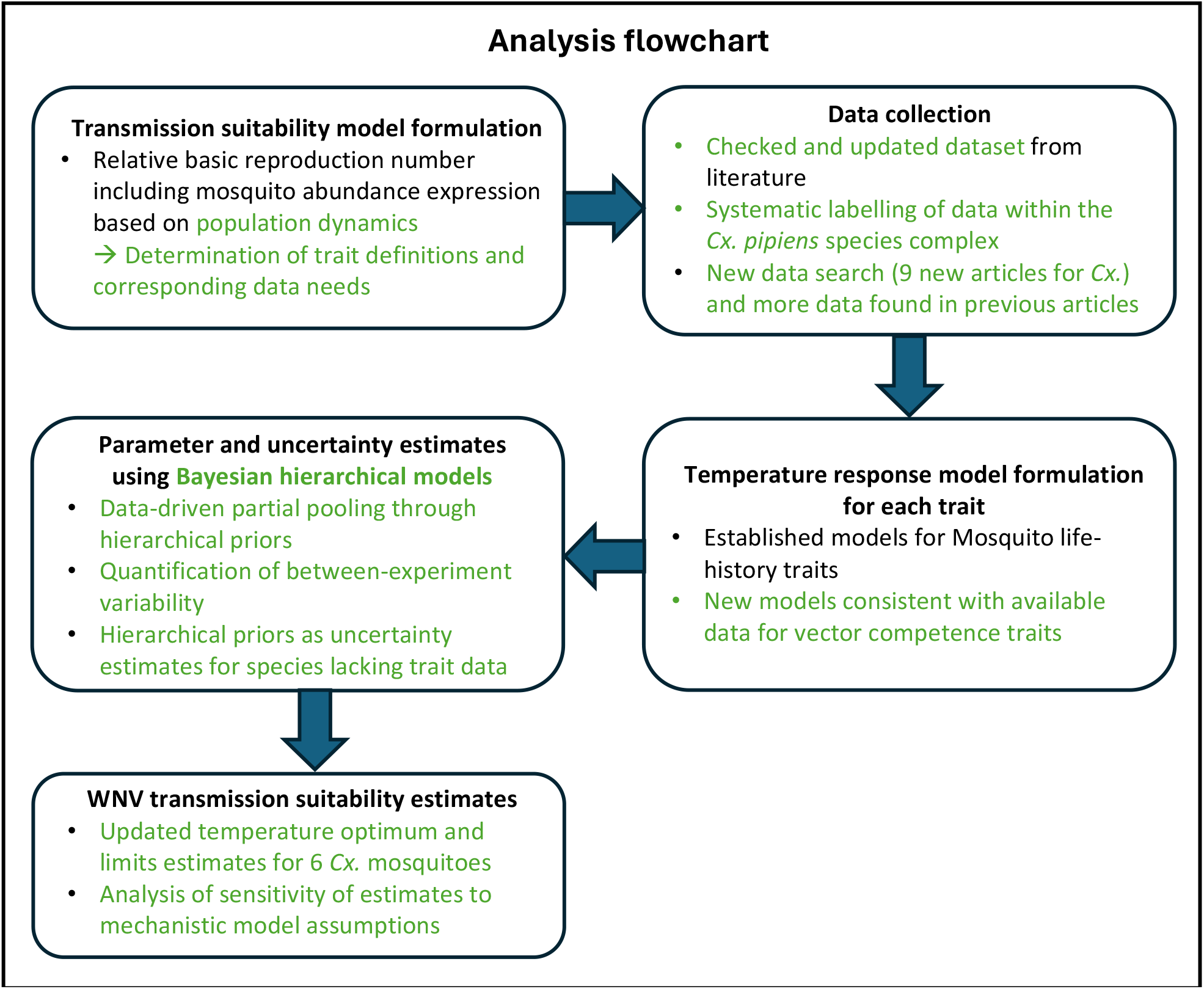
Graphical representation of the process of our analysis. Key novelties in our appraoch compared to previous analyses of MBD thermal biology are highlighted in green.

### Mathematical models of WNV transmission suitability

The basic reproduction number *R*_0_, i.e., the expected number of secondary infections produced by a single infected host in a completely susceptible population (and related variables such as vectorial capacity) is arguably one of the most important metrics in infectious disease epidemiology and has been used in the past to capture the effect of temperature on different mosquito-borne diseases [19,21]. Here, we modelled the impact of temperature on transmission suitability by considering a relative version of the basic reproduction number. Figure 2 illustrates the WNV transmission cycle and *Cx*. mosquito life cycle underlying our models, including the temperature-dependent parameters influencing both processes.

**Figure 2.**
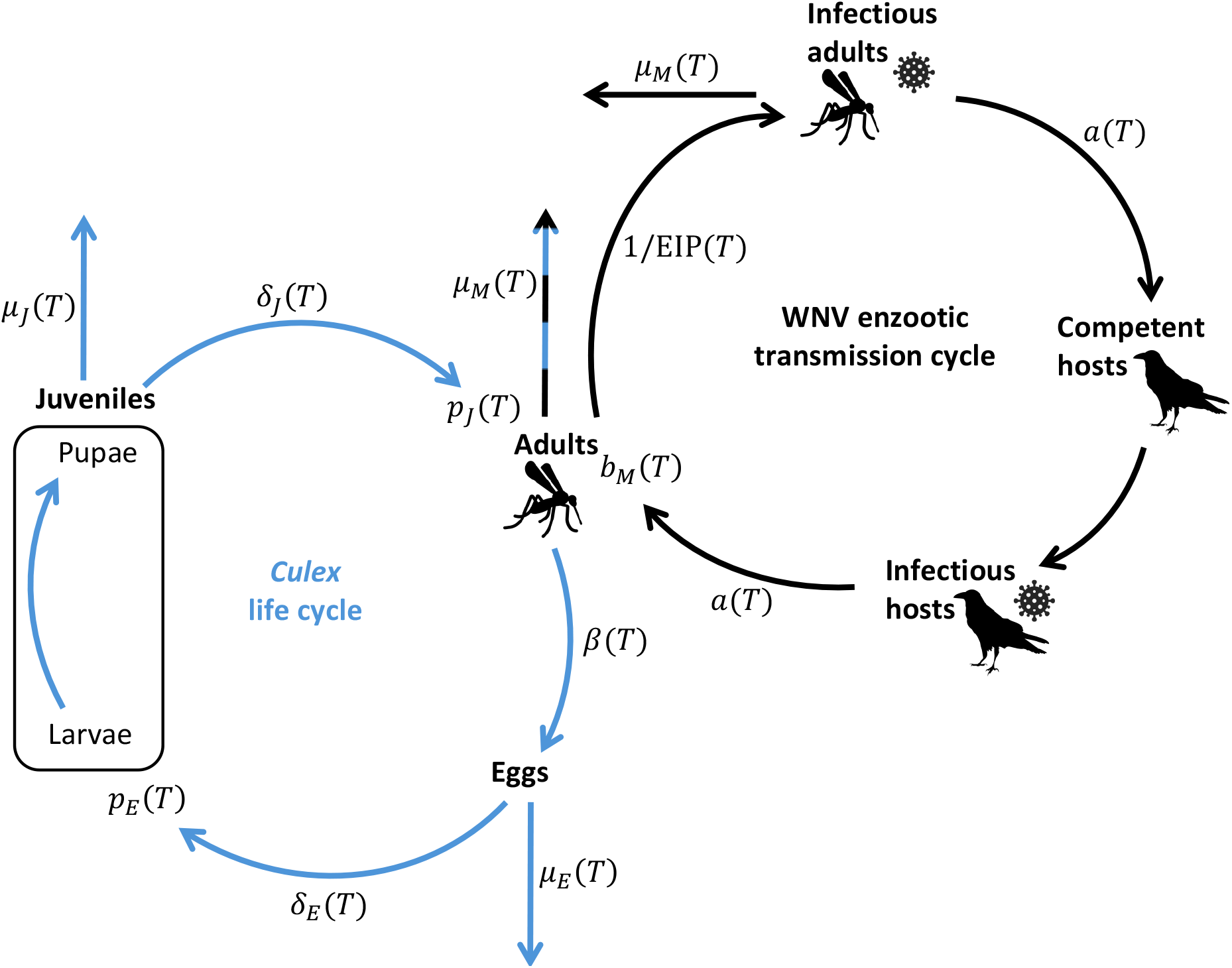
*Cx*. life cycle (blue) and WNV transmission cycle (black) as well as the temperature-dependent parameters involved in both processes. Parameters to the side of the arrows represent transition rates. Parameters at the end of the arrows represent transition probabilites. Temperature-independent parameters are left-out of the figure. A detailed description of the temperature-dependent parameters can be found in Table 1.

We start by considering a classical Ross-Macdonald equation [33,34] that models *R*_0_ as given by

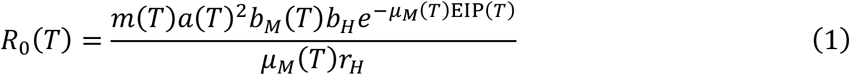

which incorporates the following temperature(*T*)-dependent mosquito-pathogen traits:

- Adult mosquito biting rate *a*(*T*)
- Adult mosquito mortality rate *μ*_*M*_(*T*)
- Mosquito infection probability *b*_*M*_(*T*) (the competence of mosquitoes to develop a midgut infection after exposure to infected blood)
- Extrinsic incubation period EIP(*T*) (average time required for competent mosquitoes to develop a salivary gland infection after exposure to infected blood)
- Mosquito to host ratio *m*(*T*) = *M*(*T*)/*H*

Additionally, the *R*_0_ model includes the temperature-independent host competence *b*_*H*_ and host recovery rate *r*_*H*_. The model could be further extended by other temperature-independent traits such as host preference [25]. However, instead of focusing on the classical interpretation of *R*_0_ as a threshold quantity we follow previous works [19,20,25,26] and consider a relative version of *R*_0_ that isolates the impact of temperature via the nonlinear interaction of mosquito-pathogen traits. To this end, we remove the temperature-independent host competence *b*_*h*_, host recovery rate *r*_*h*_, and host density *H* from Equation (1) and get the following relative version of *R*_0_:

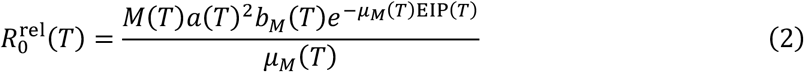

We normalise the resulting metric 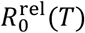 to [0,1] by dividing by the maximum obtained value with respect to temperature.

**Table 1.** Description of the temperature-dependent parameters appearing in 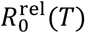 and the mosquito population dynamics model.

| Parameter (unit) | Description | Data type used for fitting/Calculation |
| --- | --- | --- |
| $a(T)$ (1/time) | Adult mosquito biting rate | The inverse of the gonotrophic cycle duration as measured from blood meal ingestion to egg laying |
| $lf(T)$ (time) | Adult mosquito lifespan | Adult mosquito lifespan observations |
| $\mu_M(T)$ (1/time) | Adult mosquito mortality rate | Derived via $\mu_M(T) = 1/lf(T)$ |
| $\delta_J(T)$ (1/time) | Juvenile mosquito development rate | The inverse of the time to develop from egg hatch to adult emergence (encompassing the larva and pupa stage) |
| $p_J(T)$ (dimensionless) | Probability of surviving larva and pupa development stage | The percentage of juveniles surviving from egg hatch to adult emergence (encompassing the larva and pupa stage) |
| $\mu_J(T)$ (1/time) | Juvenile mosquito mortality rate | Can be approximated via $p_J(T) = \frac{\delta_J(T)}{\delta_J(T) + \mu_J(T)}$ |
| $\delta_E(T)$ (1/time) | Egg hatching rate | The inverse of the time from egg laying to larva emergence |
| $p_E(T)$ (dimensionless) | Egg viability: probability that egg hatches | The percentage off eggs hatching |
| $\mu_E(T)$ (1/time) | Egg mortality rate | Can be approximated via $p_E(T) = \frac{\delta_E(T)}{\delta_E(T) + \mu_E(T)}$ |
| $\beta_{ER}$ (dimensionless) | Number of eggs per egg raft | Egg count per egg raft |
| $\beta(T)$ (1/time) | Female mosquito egg laying rate | Derived via $\beta(T) = a(T)\beta_{ER}$ |
| $b_M(T)$ (dimensionless) | Mosquito infection probability: competence of mosquitoes to develop a midgut infection after exposure to infected blood | Data from WNV vector competence studies testing mosquitoes for virus in their bodies after exposure to infected blood |
| $EIP(T)$ (time) | Extrinsic incubation period: average time required for competent mosquitoes to develop a salivary gland infection after exposure to infected blood | Data from WNV vector competence studies testing mosquitoes for virus in their salivary glands at different time points after exposure to infected blood |

To capture the impact of temperature on mosquito abundance *M*(*T*), we consider a simple stage-structured mosquito population dynamics model:

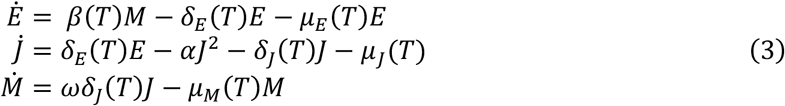

The model describes the dynamics across mosquito eggs *E*, juveniles *J* (encompassing larvae and pupae), and female adults *M* and depends on the following additional temperature-dependent mosquito life-history traits:

- Adult mosquito egg laying rate *β*(*T*)
- Egg hatching rate *δ*_*E*_(*T*)
- Egg mortality rate *μ*_*E*_(*T*)
- Juvenile mosquito development rate *δ*_*J*_(*T*)
- Juvenile mortality rate *μ*_*J*_(*T*)

The egg hatching and juvenile development rate in relation to the egg and juvenile mortality rate relate to egg viability (the percentage of eggs hatching) *p*_*E*_(*T*) and juvenile survival from egg hatch to adult emergence *p*_*J*_(*T*) via:

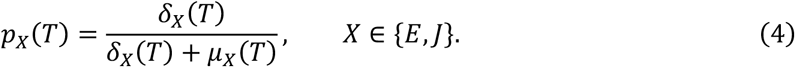

We further define *p*_*EJ*_(*T*) = *p*_*E*_(*T*)*p*_*J*_(*T*) as the probability of surviving both the egg and the juvenile stage.

The decision to separate the aquatic stage into two stages represented by eggs and juveniles (larvae and pupae combined) was driven by data availability. A more detailed separation of the different stages would need experiments that report trait data for larvae separate from pupae or even on different larvae instar stages [35,36]. However, such a model would not allow using the large body of experiments that report development times and survival only on the combined larvae and pupae stage [26,37–50] (see also Table 10 in SI9). The parameter *ω* represents the proportion of female mosquitoes at adult emergence and is assumed to be 0.5. Furthermore, the model incorporates a competition-driven juvenile mosquito mortality controlled by the parameter *α*. Competition in the juvenile aquatic stage is a common phenomenon leading to increased mortality and prolonged development times as well as carry-over effects on adult traits [41–43,51,52]. Here we incorporate the effect of increased competition-driven mortality at high larva density using a quadratic term −*αJ*^2^ which assumes pair-wise individual interactions driving competition [53,54]. In the above model, this mechanism prevents unlimited mosquito population growth.

We use the adult female demographic equilibrium of the dynamic model (3) given by

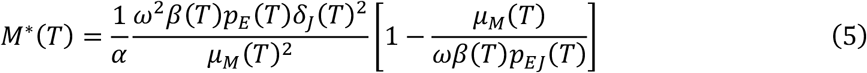

as a proxy for mosquito abundance in the 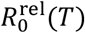 model (2) whereby *M*^∗^(*T*) = 0 if the population reproduction number

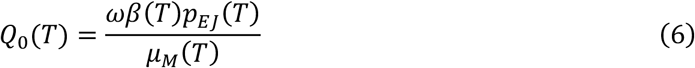

is less than or equal to one. The realized value of *α* depends on the availability of suitable space for juvenile development which we do not further consider in this work. At the demographic equilibrium, the factor 1/*α* is simply a scaling factor, and therefore the normalised 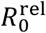 is invariant to changes in *α*.

The 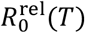 metric (2) describes a temperature-dependent transmission risk space that predicts a temperature at which WNV transmission would be optimized 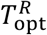 (i.e., where 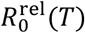 reaches one) and approximates lower and upper temperature limits 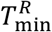 and 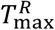 (i.e., where 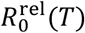becomes zero). Since we model the mosquito infection probability *b*_*M*_(*T*) as strictly positive (see following sections), it is straightforward to see that 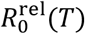 is zero if and only if *M*^∗^(*T*) is zero. Therefore, these temperature limits can be interpreted as limits for the mosquito population to thrive. If temperatures stay outside these limits for an extended period, mosquito populations, and thus transmission cycles, could unlikely be sustained. The temperature interval that would support disease transmission is located within these temperature limits but depends not only on 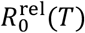 being larger than zero but on the absolute *R*_0_ (1) being greater than one. This cannot be predicted from 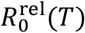 alone because it depends on additional location-specific, temperature-independent factors such as host population density and susceptibility.

The mosquito population dynamics model (3) underlying the adult mosquito equilibrium expression (5) makes specific assumptions on mosquito ecology. To test if our 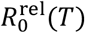 estimates might be affected by the mosquito model selection, we consider alternative formulations from the literature that differ from system (3) in their incorporation of intra-specific competition into the mosquito population dynamic model and thus for integrating the impact of temperature on the mosquito-to-host ratio [55–60]. We also consider an expression that lacks a rigorous link to mosquito population dynamics but has been used in a series of previous analyses of MBD thermal biology [19,20,22,23,25,26,28–30,32,61]. The equations for these different models can be found in SI7 in S1 Text, where we also discuss their theoretical foundation.

### Data collection

An overview of the temperature-dependent mosquito-pathogen traits appearing in 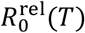 (2) and the mosquito population dynamics model (3) can be found in Table 1. This table also includes a description of the data type used to fit the temperature response of each trait or how the trait is derived from other traits.

The starting point of our data collection is a previously published dataset [20], which compiled outcomes of experimental studies testing the temperature dependence of multiple mosquito-pathogen traits. This dataset primarily focused on viruses causing MBDs in the temperate areas and data on mosquito species capable of transmitting these pathogens. Many of these species belong to the genus *Culex*, but the dataset also contains information about temperate species from the genera *Aedes* and *Culiseta*. Here, we provide an updated and enriched version of this dataset.

As a first step to gathering the updated dataset, we double-checked each entry in the original dataset [20] with the content of the primary articles. This screening led to several changes to the dataset, including the correction of typos and the application of conceptional differences when handling the data. A detailed description of these changes can be found in SI8 in S1 Text. To ensure compiling the most up-to-date data for the main genus of interest in our analysis, we ran a literature search on PubMed and Web of Science using the search query (*Culex*) AND (temperature* OR thermal*) restricted to 2019-2023. Following the inclusion criteria established for collecting the initial dataset [20], we only incorporated results from experiments that assessed trait performance across a minimum of three distinct constant temperature settings within controlled environments. This search identified three studies [35,51,62] that matched the inclusion criteria. We also added experimental data from a study published after we conducted our literature search [26]. Additionally, during the search we identified recent review articles [28,63] from which we scanned the reference lists to uncover additional relevant studies. This approach allowed us to incorporate data from further experimental studies [37,64–67]. During this step we made an exception to our inclusion criteria and included data on the lifespan of *Cx. pipiens pallens* from a study that only tested two temperature settings due to data scarcity on this species-trait combination. The final dataset entails data from 40 experimental studies on 8 different mosquito-pathogen traits across 15 mosquito species [26,35–51,62,64–84]. Whenever data was only available as figures, we used a plot digitizer tool for data extraction [85]. An overview of the studies included in our final data collection can be found in Table SI9.1 in S1 Text.

In our analysis, we focus on six *Cx*. species (*Cx. pipiens, Cx. quinquefasciatus, Cx. pipiens molestus, Cx. pipiens pallens, Cx. restuans, Cx. tarsalis*) and their potential to transmit WNV. Nevertheless, we keep life-history information on the other temperate *Culex, Aedes*, and *Culiseta* species in the dataset. This additional data helps to derive hierarchical priors during model fitting and as a side product of our models, we provide updated trait temperature response estimates for these species as well. To note, our use of the classification “species” is taxonomically inaccurate and applied for simplicity. In fact, the mosquito *Cx. pipiens pallens* represents a *Cx. pipiens* x *Cx. quinquefasciatus* subspecies and *Cx. pipiens molestus* an ecotype of *Cx. pipiens* (for more details, see SI6 in S1 Text).

### Modelling the temperature-response of mosquito-pathogen traits

For each mosquito-pathogen trait we chose a suitable parametric function by visual inspection of the data and by drawing on prior works and theories in mosquito thermal biology [19,20,23,86]. Below, we define the functions used for fitting these different traits. A more detailed reasoning for our modelling choices also highlighting important differences in our approach to earlier works [20,22,23,25,26,29–31] can be found in SI1 in S1 Text.

We model the juvenile mosquito development rate *δ*_*J*_(*T*), egg hatching rate *δ*_*E*_(*T*), and adult biting rate *a*(*T*) with a modified Briére function which describes a left-skewed unimodal response to temperature [87]:

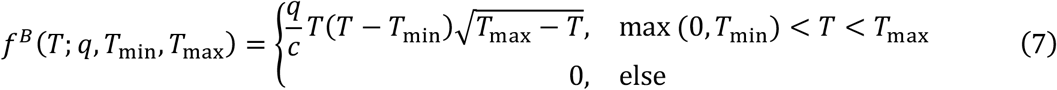

The parameters *q, T*_mim_, *T*_max_ are the estimation targets whereas *c* represents a constant that acts to scale *q* on a similar scale as *T*_mim_ and *T*_max_ which helps to improve the speed and stability of the fitting procedure. In addition, we set the scaling factor such that the parameter *q* obtains similar values between the three traits.

For juvenile survival *p*_*J*_(*T*) and egg viability *p*_*E*_(*T*) we fit a modified quadratic function that describes a symmetric unimodal temperature response:

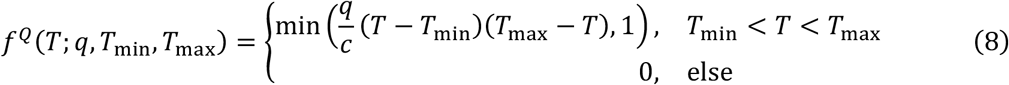

Here again, we incorporate the constant *c* to bring the parameter *q* on a similar scale as *T*_mim_ and *T*_max_.

For adult mosquito lifespan lf(*T*) we follow previous work [20,26] and model it by a linearly decreasing function truncated at zero:

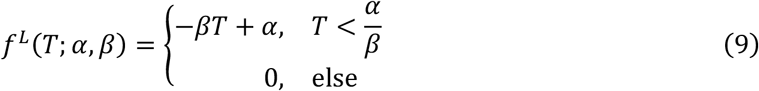

We estimated the parameters *β* and *T*_max_ ≔ *α*/*β* since we found it more intuitive to define priors for the transformed parameter *T*_max_ than for the intercept *α*. As a conservative approach to trait performance at low temperatures, we plateau the linear function after model fitting at the lowest observed temperature point in the dataset across all species (14°C).

The female mosquito egg-laying rate *β*(*T*) is derived by multiplying the biting rate *a*(*T*) with the number of eggs per egg raft *β*_ER_. Data on eggs per egg raft *β*_*E*R_ of *Cx*. mosquitoes under different temperature settings is very limited [26,39,70]. The data is confined to the temperature range 15-30°C and while the data on *Cx. pipiens molestus* shows the most notable reduction in *β*_*E*R_ at high temperature, the remaining data indicates that the trait is somewhat stable over the observed temperature range, although with tendencies for the highest trait values at intermediate temperatures. Overall, we found the information on *β*_ER_ too scarce to fit a temperature-dependent function. Therefore, we decided to set this trait to a constant value determined by the mean across all observations in the dataset (given by 140).

We model the mosquito infection probability *b*_*M*_(*T*) using a sigmoidal function:

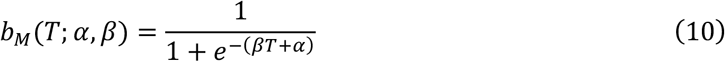

As a model for the extrinsic incubation period EIP(*T*) we chose an exponential decay function:

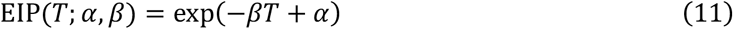

### Statistical inference of temperature-response functions

To estimate the temperature response function parameters for each trait, we implemented Bayesian hierarchical models. For mosquito-life history traits, our models include species and experiment identity as hierarchical levels of parameter estimates while we estimated the pathogen-related traits using experiment identity as the only hierarchical level. We assigned different experiment identities if the data came from separate articles or when an article investigated the temperature response of different populations of the same species (such as different strains [69], laboratory versus field-derived populations [44], or field-derived populations of different geographical origin [46]). Below, we provide a general summary of our models. The formal mathematical descriptions of our Bayesian models as well as prior and hyperprior specifications can be found in S3 in S1 File.

### Mosquito life-history traits

The hierarchical models for life-history traits include two hierarchical levels: species and experiment identity. We model the trait performance *Y*_*i,j,T*_ of species *i* in experiment *j* at a particular temperature *T* using a normal distribution likelihood with mean given by the trait-specific parametric function evaluated at the given temperature (see Equations (7)-(9)) with a parameter vector θ_*i,j*_ (e.g., in case of juvenile development rate: 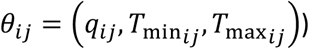 depending on species and experiment identity and an estimated standard deviation *s*:

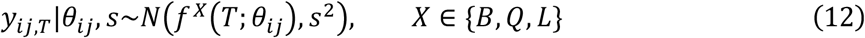

We assign the model parameter vector θ_*i,j*_ a multivariate normal hierarchical prior with mean parameter vector θ_*i*_, which represents the mean parameter realizations for species *i*, and diagonal covariance matrix defined by between-experiment standard deviations σ^exp^, measuring the variability of parameter realizations across experiments on the same species:

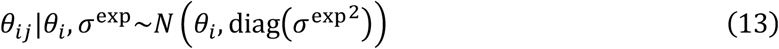

The mean parameter realizations θ_*i*_ are assigned another multivariate normal hierarchical prior

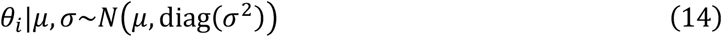

with population-level means *μ*, reflecting the mean parameter realization across species, and between-species standard deviations σ, measuring the variability of parameter realizations across species.

We use the mean parameter realizations θ_*i*_ to calculate the expected trait temperature response of each species. We include the between-experiment variability σ^exp^ since already a visual inspection of the data indicated a substantial variation in the temperature response between separate experiments on the same trait and species. Neglecting these experiment effects ignores statistical dependencies within the data which can lead to biased and overconfident model fits (see SI5 in S1 Text).

Note that *μ*, σ, σ^exp^ are vectors whose dimensionality depends on the considered trait (e.g., in case of juvenile development rate: 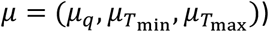. Our choice of diagonal covariance matrices implies that we assign independent normal priors to the individual parameters in θ_*i,j*_. To ensure that the estimates of the parameter *q* in case of the Briére and quadratic function as well as the parameter *β* in case of the linear model for mosquito adult lifespan are non-negative, the normal hierarchical priors were placed on log-transformed versions of these parameters. For simplicity of presentation, we neglected this detail in our notation above (see SI3 in S1 Text for all details).

We defined the hyperpriors of hierarchical prior parameters (population-level means, between-species standard deviations, and between-experiment standard deviations) following different strategies depending on trait data availability. For traits with extensive data available (juvenile mosquito development rate and survival) we used vague hyperpriors on all three types of hierarchical prior parameters. We loosely informed these hyperpriors based on the extensive literature on the thermal biology of mosquito-borne diseases to guide parameter estimates to biologically reasonable regions [19]. For less studied traits (adult mosquito lifespan, egg hatching rate, adult mosquito biting rate, and egg viability) it was not possible to derive reasonable estimates without including additional prior knowledge. Therefore, for these traits we only defined vague hyperpriors on the population-level means. In case of mosquito adult lifespan, we also assigned vague hyperpriors to the between-species and between-experiment standard deviations of the slope parameter of the linear function fitted for this trait. For all other between-species and between-experiment standard deviations, we defined informative hyperpriors by drawing on the estimates derived for juvenile mosquito development rate and survival. To this end, we fitted gamma distributions to the posterior draws generated for the between-species and between-experiment standard deviations of these traits. These were then used as informative priors in the models of the less studied traits, ensuring biologically reasonable estimates. We informed the hyperpriors of egg hatching rate and adult mosquito biting rate by the estimates generated for juvenile mosquito development rate since all three traits represent metabolic rates and are modelled by a Briére function. For the hyperpriors of egg viability, we used the gamma distributions derived from the juvenile mosquito survival fit due to the biological similarity of these traits and since both traits were fitted using a quadratic function. We also used the fit for the between-species and between-experiment standard deviation of the temperature maximum parameter of mosquito juvenile survival to inform the hyperpriors of the temperature maximum parameter in the mosquito lifespan model.

### Mosquito infection probability and extrinsic incubation period

For the pathogen-related traits, we used a simpler hierarchical model with experiment identity as the only hierarchical level. We did not have sufficient data available to estimate parameters for groups of experiments on the same mosquito species or on the same mosquito species and virus strain combination. Data on the mosquito infection probability was available in binomial form. Therefore, we model the number *n*_*j,T*_ of mosquitoes with a midgut infection (virus detected in body) upon the number *N*_*j,T*_ of all mosquitoes tested in experiment *j* at temperature *T* using a binomial likelihood with success probability given by the sigmoidal function that we chose for this trait with parameter vector θ_*j*_ = (*α*_*j*_, *β*_*j*_):

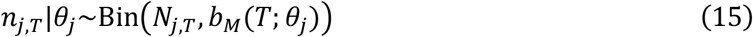

For the extrinsic incubation period, data was available as the percentage of “transmitting mosquitoes” (virus detected in salivary glands) upon all mosquitoes with a midgut infection (virus detected in body) in dependence on the days post infection *t* and the temperature setting *T*. We model the relationship between days post infection *t* and the percentage *p*_*j,t,T*_ of transmitting mosquitoes in experiment *j* at temperature *T* using a normal likelihood with an estimated standard deviation *s* and mean given by a cumulative Gaussian distribution Φ. We estimate the standard deviation *s*_EIP,_ of the cumulative gaussian and model its mean (i.e., the point in time at which 50% of infected mosquitoes are expected to have a salivary gland infection) with the exponential decay function that we selected for the extrinsic incubation period trait:

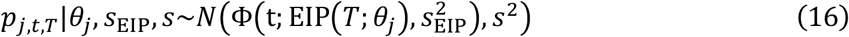

For both pathogen-related traits, we assigned a multivariate normal hierarchical prior with population-level means *μ* and diagonal covariance matrix defined by between-experiment standard deviations σ^exp^ to the parameter vector θ_*j*_:

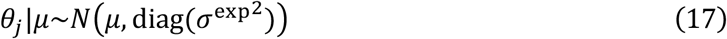

The population-level means of each parameter were assigned vague hyperpriors. However, for both traits data from only a few experiments were available, which is why it was impossible to derive reasonable estimates of the between-experiment standard deviations. In contrast to mosquito life-history traits, we did also not have any prior knowledge of the between-experiment variability of parameter estimates from biologically similar traits. Therefore, we fixed these parameters to a specific value. These values were chosen following a conservative approach that enforces some amount of pooling of the parameter estimates across experiments but also allows for uncertainty of the temperature response estimate at the population-level (i.e., the temperature response generated by sampling from the hierarchical prior). Since we set these between-experiment standard deviations to subjectively chosen values we investigated the sensitivity of our estimates to changes in these values (see SI4).

### Implementation

We fitted the Bayesian models using Stan [88] through the Rstan package [89]. Stan provides full Bayesian inference via Markov chain Monte Carlo (MCMC) methods. Here, we used the No-U-Turn sampler, a form of Hamiltonian Monte Carlo sampling. For each trait, we ran 4 MCMC chains for 4000 iterations, of which 2000 iterations served as warmup. We checked the Markov chains for convergence using the Gelman-Rubin statistic and by visual inspection. For the six WNV vectors, we obtained species-specific posterior estimates for 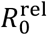 by applying the posterior samples of each mosquito-pathogen trait model fit to Equation (2). From this posterior distribution, we calculated mean estimates and 95% credible intervals (95% CIs) for the temperature limits 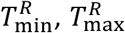and the temperature optima 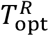 of WNV transmission suitability for each species.

To build the 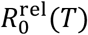 model for species that lacked data on one or more life-history traits, we substituted missing temperature response estimates by population-level estimates generated by sampling parameter values from the hierarchical prior *N*(*μ*, diag(σ^2^)) (Equation 14). In the context of our Bayesian models these hierarchical priors represent the uncertainty about a species’ expected trait temperature response in the absence of data on this specific species (see Figure SI4.1 in S1 Text). For mosquito infection probability, we used the population-level estimates generated by sampling from the hierarchical prior *N*(*μ*, diag(σ^exp2^)) (Equation 17) in each species’ 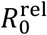 model since for this trait only data for *Cx. pipiens* is available. In case of the extrinsic incubation period, we used the fits derived for WN02 in *Cx. pipiens* in the 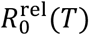 model for *Cx. pipiens*, NY99 in *Cx. tarsalis* for *Cx. tarsalis*, and the population-level estimates for all other species (see Figure SI4.2 in S1 Text).

## Results

Here, we present the results on temperature suitability and estimates for the expected trait temperature response for the main species of interest in the transmission of WNV: *Cx. pipiens, Cx. quinquefasciatus, Cx. pipiens molestus, Cx. pipiens pallens, Cx. restuans, Cx. tarsalis*. An overview of all trait temperature response parameter estimates including other species can be found in Tables SI2.1-8 in S1 Text.

### 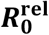 transmission suitability temperature response

Across the six WNV vectors considered here, we find striking similarities in the temperature-sensitivity of 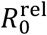 (Figure 3 and Table 2). We predict that WNV transmission suitability peaks at intermediate temperatures with mean estimates across species ranging 23.5-25.6°C and broadly overlapping 95% CIs. Similarly, our estimates of temperature limits indicate uniformity between species with the lowest mean estimate for the upper temperature limit at 31.9°C (95% CI: 26.8-35.1°C) for *Cx. restuans* and the highest at 34.5°C for *Cx. tarsalis* (95% CI: 29.6-37.3°C). The mean estimates for lower temperature limits range from the lowest estimate for *Cx. restuans* at 9.4°C (95% CI: 4.9-16.1°C) to the highest estimate for *Cx. quinquefasciatus at* 12.5°C (95% CI: 9.3-15.6°C). When a species lacked data on a specific trait, we substituted its temperature response with a population-level estimate generated by sampling from hierarchical priors. Consequently, the uncertainty in temperature optima and limits varied by species depending on trait data availability. Especially, species lacking specific data on egg viability (*Cx. pipiens, Cx. restuans, Cx. tarsalis*) showed relatively high uncertainty in temperature limits estimates due to the highly uncertain population-level estimate for this trait (see Figure SI4.1 in S1 Text). Among these, *Cx. restuans* exhibited the highest uncertainty, also in its temperature optimum estimate, as it lacked specific data on adult biting rate, egg viability, and pathogen-related traits, and had limited data available on adult mosquito lifespan.

**Table 2.** Overview of estimates of the optimal temperature and temperature limits of 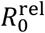 for WNV in *Cx*. mosquitoes from our study and previous studies.

| WNV in | $T_{\text{min}}^R$ | $T_{\text{opt}}^R$ | $T_{\text{max}}^R$ |
| --- | --- | --- | --- |
| **Our study** |  |  |  |
| *Cx. pipiens* | 9.8 (6.6-16) | 24.5 (23.0-25.9) | 33.6 (30.3-35.4) |
| *Cx. quinquefasciatus* | 12.5 (9.3-15.6) | 24.5 (23.2-25.8) | 33.8 (31.8-35.6) |
| *Cx. pipiens molestus* | 10.7 (8.5-12.9) | 23.7 (21.8-25.6) | 32.1 (29.6-34.1) |
| *Cx. pipiens pallens* | 10.2 (8.0-12.7) | 24.9 (22.9-27.2) | 34.4 (31.3-37.5) |
| *Cx. restuans* | 9.4 (4.9-16.1) | 23.5 (20.4-26.0) | 31.9 (26.8-35.1) |
| *Cx. tarsalis* | 11.3 (8.4-16.0) | 25.6 (23.4-27.8) | 34.5 (29.6-37.3) |
| **Fay et al. 2024 [26]** |  |  |  |
| *Cx. pipiens*<br>(NYS downstate) | 17.8 (15.6-20.2) | 24.6 (22.9-26.0) | 33.9 (30.6-37.2) |
| *Cx. pipiens*<br>(NYS upstate) | 16.7 (15.0-17.9) | 21.7 (20.7-22.8) | 30.2 (28.2-33.3) |
| **Di Pol et al. 2022 [25]** |  |  |  |
| *Cx. pipiens* | 14 | 23.7 | 34.3 |
| **Shocket et al. 2020 [20]** |  |  |  |
| *Cx. pipiens* | 16.8 (14.9-17.8) | 24.5 (23.6-25.5) | 34.9 (32.9-37.6) |
| *Cx. quinquefasciatus* | 19.0 (14.1-20.9) | 25.2 (23.9-27.1) | 31.8 (31.1-32.2) |
| *Cx. tarsalis* | 12.1 (9.6-15.2) | 23.9 (22.9-25.9) | 32.0 (30.6-38.6) |
| *Cx. univittatus* | 11.0 (8.0-15.3) | 23.8 (22.7-25.0) | 33.6 (31.2-36.9) |
| **McMillan et al. 2019 [24]** |  |  |  |
| *Cx. spp.* | 14.5 | 23.0 (20.8-25.1) | 34.7 |
| **Paull et al. 2017 [15]** |  |  |  |
| *Cx. pipiens* | 17 | 24.9 | — |
| *Cx. quinquefasciatus* | 17 | 24.3 | — |
| *Cx. tarsalis* | 15 | 24.9 | — |
| **Vogels et al. 2017 [27]**<br>(model not unimodal) |  |  |  |
| *Cx. pipiens* | 18 | 28 | — |
| *Cx. pipiens molestus* | 18 | 28 | — |
| **Kushmaro et al. 2015 [90]**<br>(model not unimodal) |  |  |  |
| *Cx.* and *Ae. spp.* | 18 | 35 | — |

**Figure 3.**
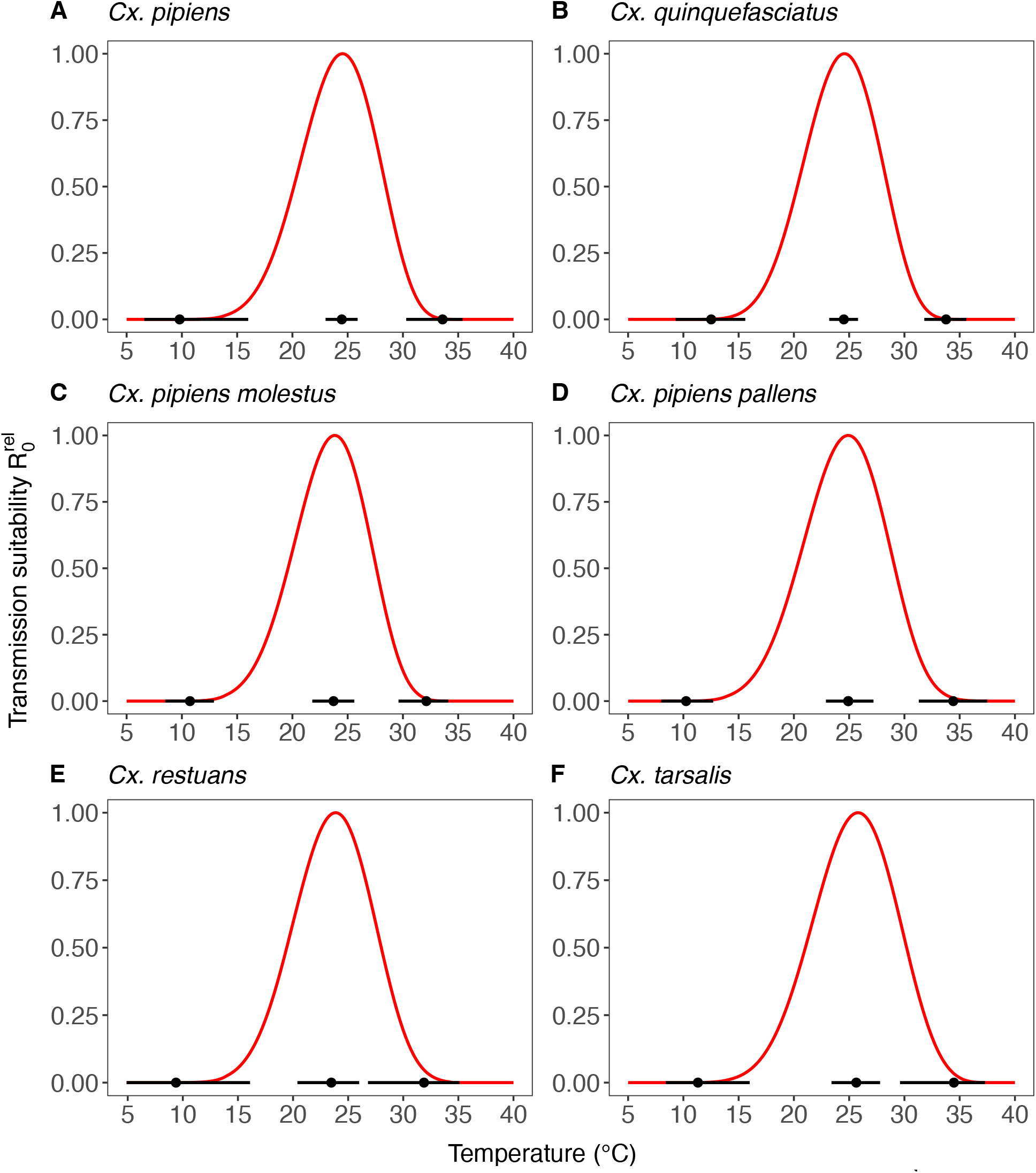
Estimated temperature response of the relative basic reproduction number 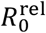 for WNV in six *Cx*. mosquitoes. Red solid lines indicate the mean temperature response. Black dots indicate mean estimates of the optimal temperature and temperature limits of 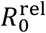. Black solid lines represent the corresponding central/equal-tailed 95% credible interval.

### Impact of mechanistic model assumptions on transmission suitability estimates

We tested whether changes to mechanistic assumptions underlying the 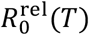 model would impact our results on WNV temperature suitability. Our main model incorporates the impact of temperature on mosquito abundance suitability through an equilibrium expression derived from a mosquito population dynamics model. We compared the results for *Cx. pipiens* using this model against four alternative models. Three alternative models use different mosquito equilibrium expressions derived from mosquito population dynamic models, each varying in their representation of mosquito intra-specific competition. The fourth alternative model uses a mosquito abundance expression previously used in various studies on MBD thermal biology [19,20,22,23,25,28–30,32], which lacks a formal connection to a mosquito population dynamics model (see SI7 in S1 Text). Figure 4 demonstrates a notable impact of mosquito modelling assumptions on the transmission suitability temperature response curve. Across the different models, the mean estimates for the temperature optimum for WNV suitability transmitted by *Cx. pipiens* vary by up to 3.1°C, with the estimate from our main model falling between those of the alternative models. The estimates of the WNV suitability temperature optimum by model are given by: 24.5°C (95% CI: 23.0-25.9°C) for the main model (using Equation (5)), 22.6°C (95% CI: 21.5-24.0°C) for alternative model 1 (using Equation (SI7.2) in S1 Text), 24.4°C (95% CI: 23.1-25.8°C) alternative model 2 (using Equation (SI7.4) in S1 Text), 25.7°C (95% CI: 24.1-27.3°C) alternative model 3 (using Equation (SI7.6) in S1 Text), and 23.5°C (95% CI: 22.2-24.8°C) for alternative model 4 (using Equation (SI7.7) in S1 Text).

**Figure 4.**
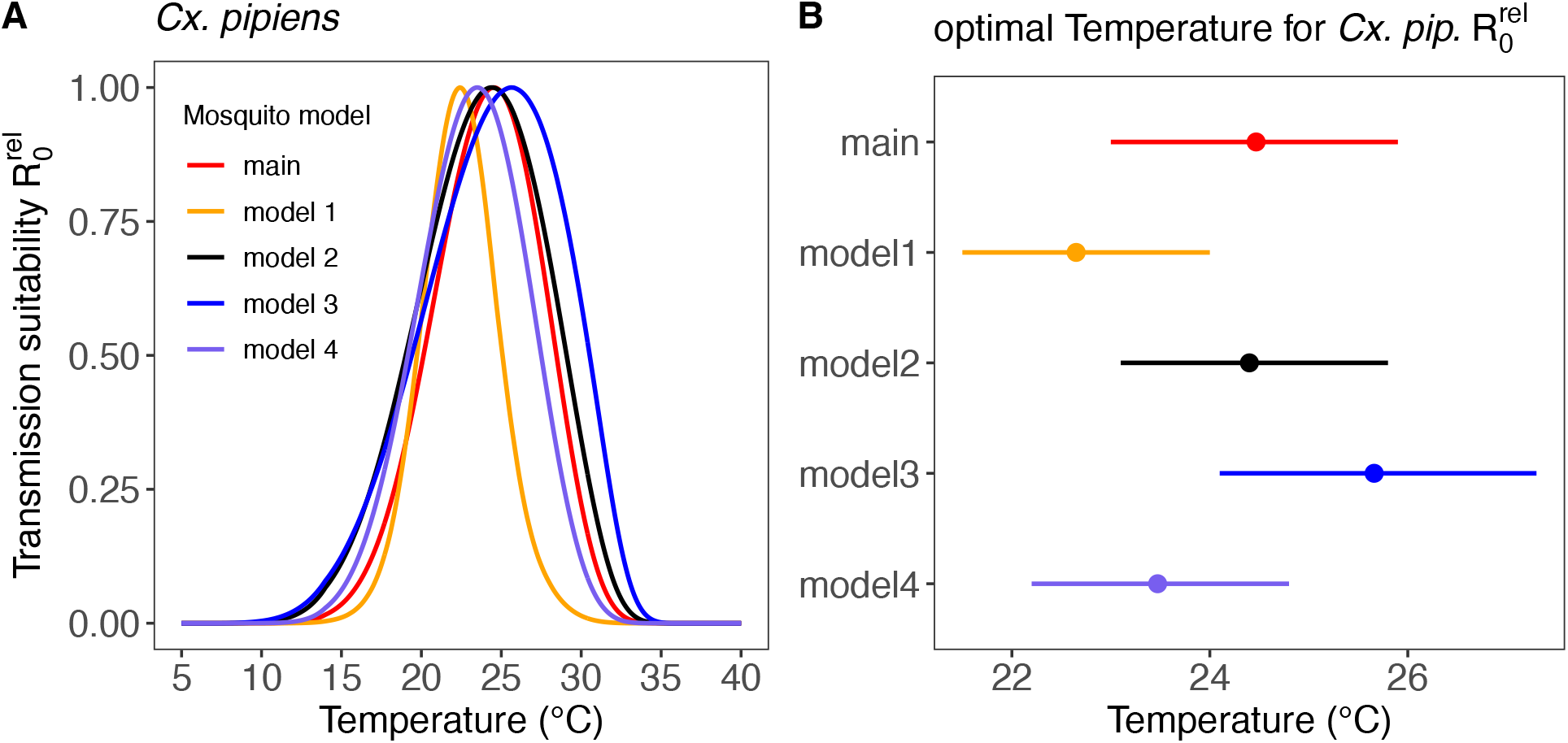
(A) Temperature response of the relative basic reproduction number 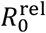 for WNV in *Cx*. pipiens contrasting the estimate from transmission suitability model introduced in the main text (see Equation (5)) as well as from four alternative model formulations from the published literature (see Equations (SI7.1-7.7) in S1 Text). (B) Mean estimates of the optimal temperature of 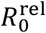 per model (dots) and the corresponding central/equal-tailed 95% credible interval (lines).

### Temperature response of mosquito life-history traits

The estimates of the expected temperature responses for each life-history trait are shown in Figure 5-10. An overview of the corresponding posterior mean parameter estimates and 95% CIs for each life-history trait can be found in Tables SI2.1-6 in S1 Text. The model fits for juvenile development rates (Figure 5), and similar for egg development rates (Figure 6), indicate relatively high temperature optima with mean estimates for juveniles ranging 33.69-35.38°C across the six *Cx*. species. Furthermore, temperature limits estimates are beyond the limits dictated by other traits with upper limit estimates ranging 41.90-43.82°C and lower limits 0.59-2.98°C. Given the little information within the data on development rates at high temperatures, uncertainty in our model fits for these traits was quite large in the high-temperature regimes. In contrast to juvenile development rates, the model fits for juvenile survival (Figure 7) indicate lower temperature optima (mean estimates ranging 20.01-24.42°C) and tighter temperature limits across the six species (upper limits ranging 34.29-38.79°C and lower 5.09-10.55°C). The temperature response fits for adult lifespan (Figure 8) indicate that some species, e.g., *Cx. pipiens* and *Cx. quinquefasciatus*, have substantially higher lifespan at lower temperatures than others, e.g., *Cx. tarsalis*. The estimate of the upper temperature limit was highest for *Cx. tarsalis* with 37.66°C (95% CI: 34.42-41.51) and lowest for *Cx. restuans* with 32.48 (95% CI: 27.34-36.98). For egg viability (Figure 9) data from only three of the *Cx*. species was available (and of four species in total) which resulted in a relatively large uncertainty in model fits. Our estimates for the temperature limit parameters of this trait ranged 33.27-39.44°C for the upper limits and 6.61-12.38°C for the lower limits across the six species. Temperature optima ranged from 19.94-25.42°C. The estimates of the temperature sensitivity of biting rates (Figure 10) were also limited by the available data and therefore resulted in a relatively large uncertainty. Our fits indicate that the upper temperature limit parameter for this trait range between 40.75-45.66°C, the lower temperature limit 0.16-1.93°C, and the temperature optimum 32.65-36.69°C between species. The 95% CIs of parameter estimates largely overlapped across species for all traits. Population-level temperature responses generated by sampling parameters from the hierachical priors determined by the estimated population-level means and between-species standard deviations, showed a relatively large uncertainty in our knowledege about the trait temperature responses of a new species in the absence of data (Figure SI4.1 in S1 Text).

**Figure 5.**
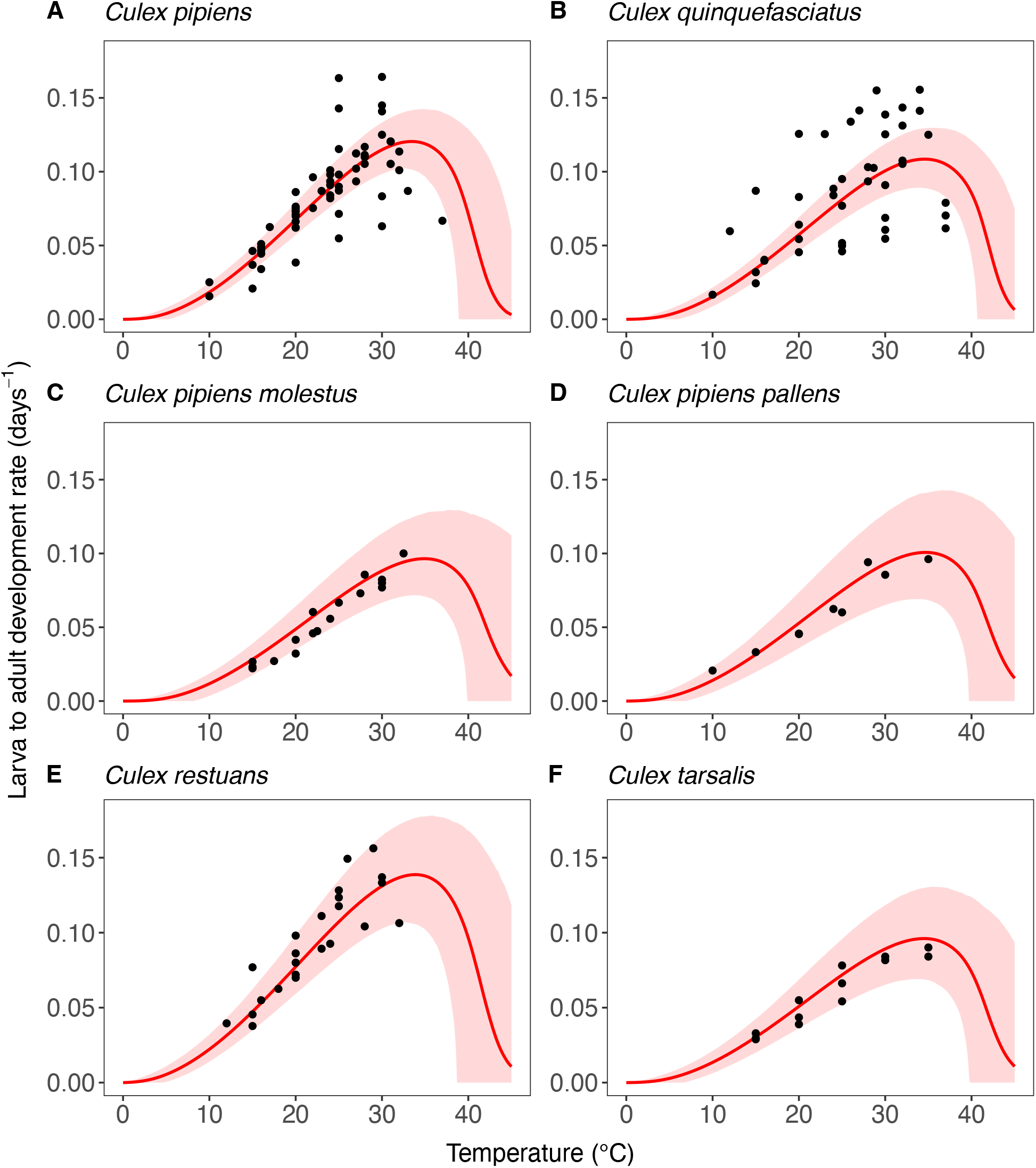
Estimates of the expected temperature response of the larva to adult development rate for six *Cx*. mosquitoes. Black dots represent data from experimental studies. Red solid lines represent posterior distribution mean model fits. Red shaded areas represent the corresponding central/equal-tailed 95% credible interval.

**Figure 6.**
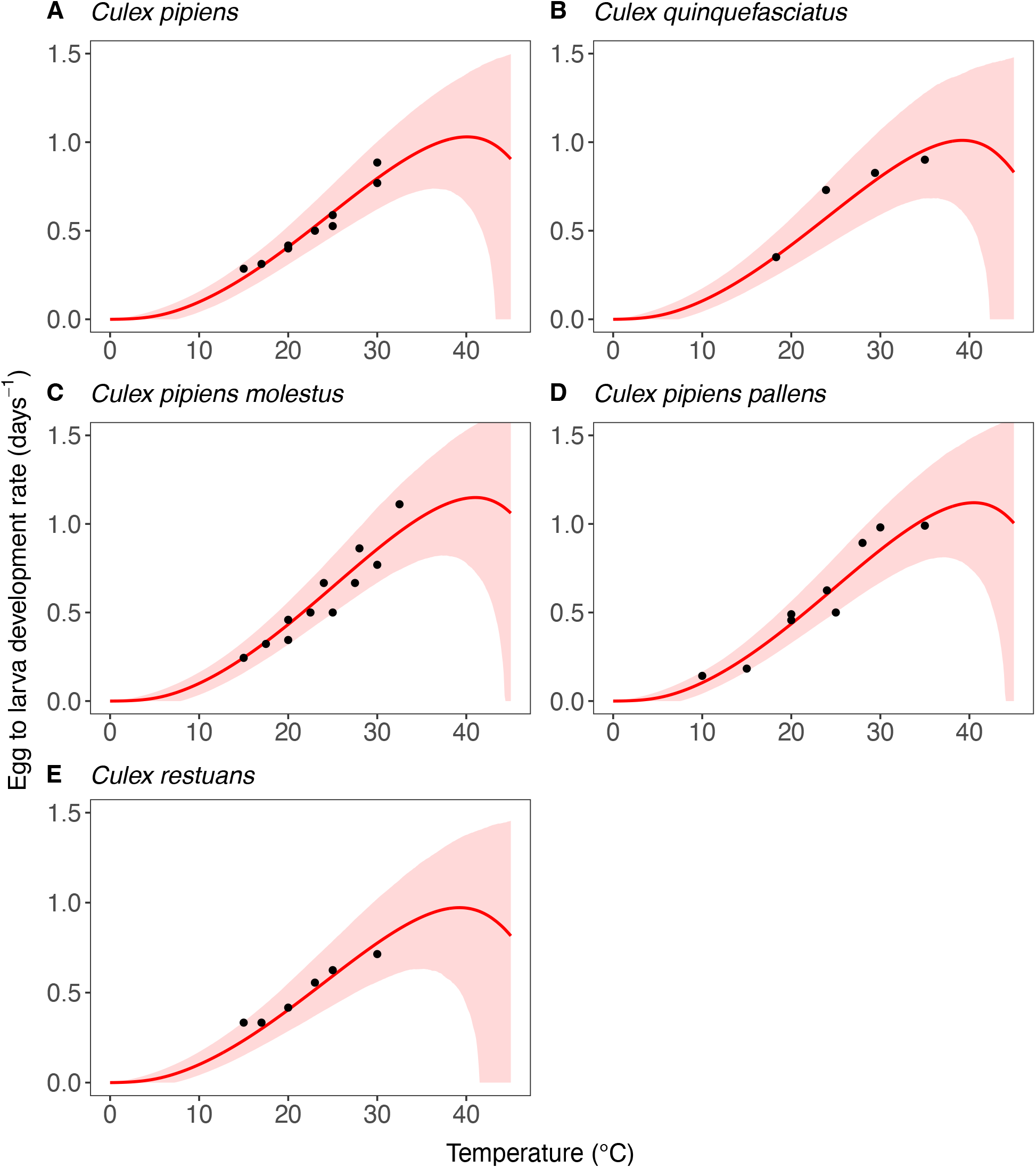
Estimates of the expected temperature response of the egg to larva development rate for five *Cx*. mosquitoes. Black dots represent data from experimental studies. Red solid lines represent posterior distribution mean model fits. Red shaded areas represent the corresponding central/equal-tailed 95% credible interval.

**Figure 7.**
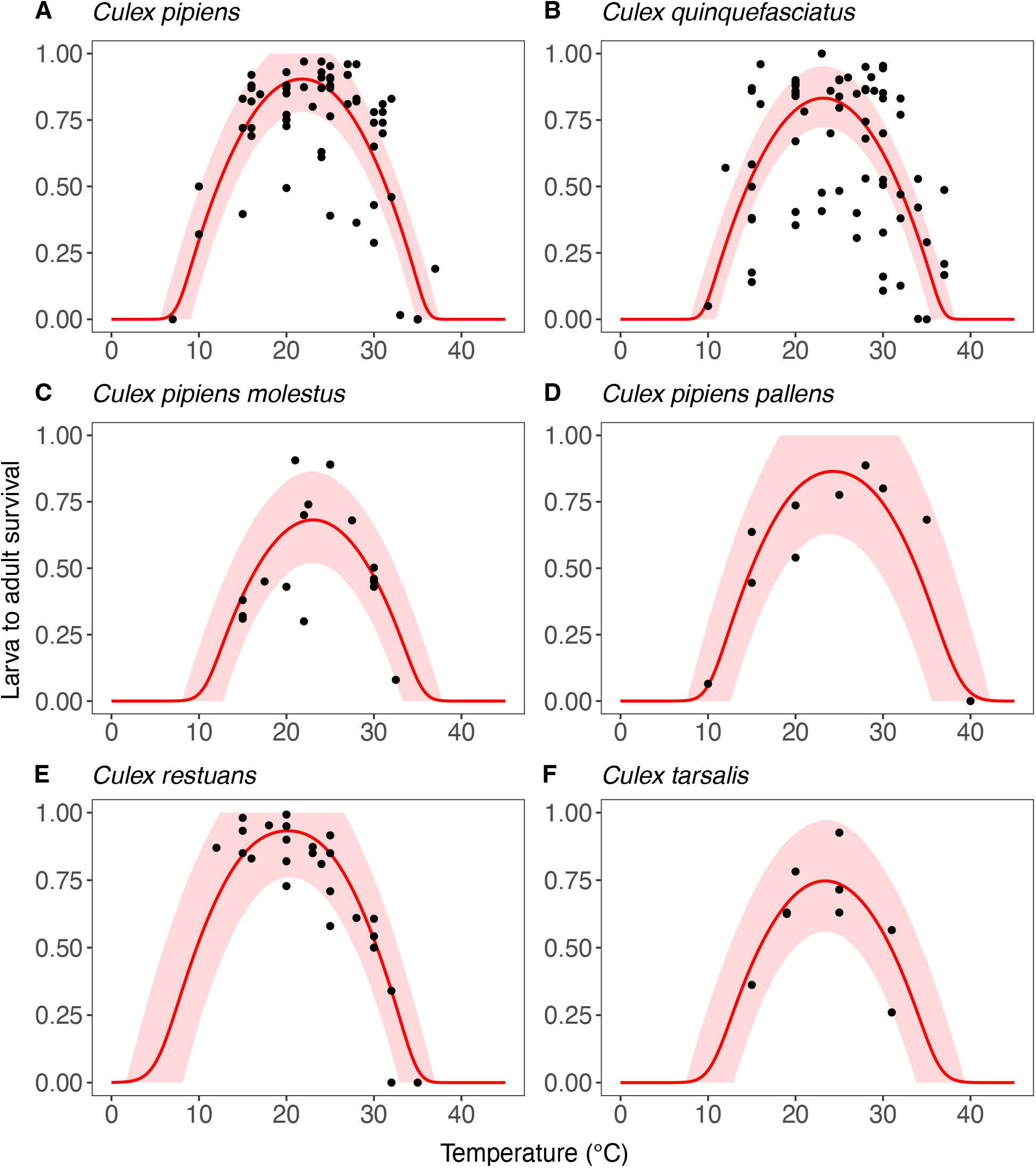
Estimates of the expected temperature response of larva to adult survival for six *Cx*. mosquitoes. Black dots represent data from experimental studies. Red solid lines represent posterior distribution mean model fits. Red shaded areas represent the corresponding central/equal-tailed 95% credible interval.

**Figure 8.**
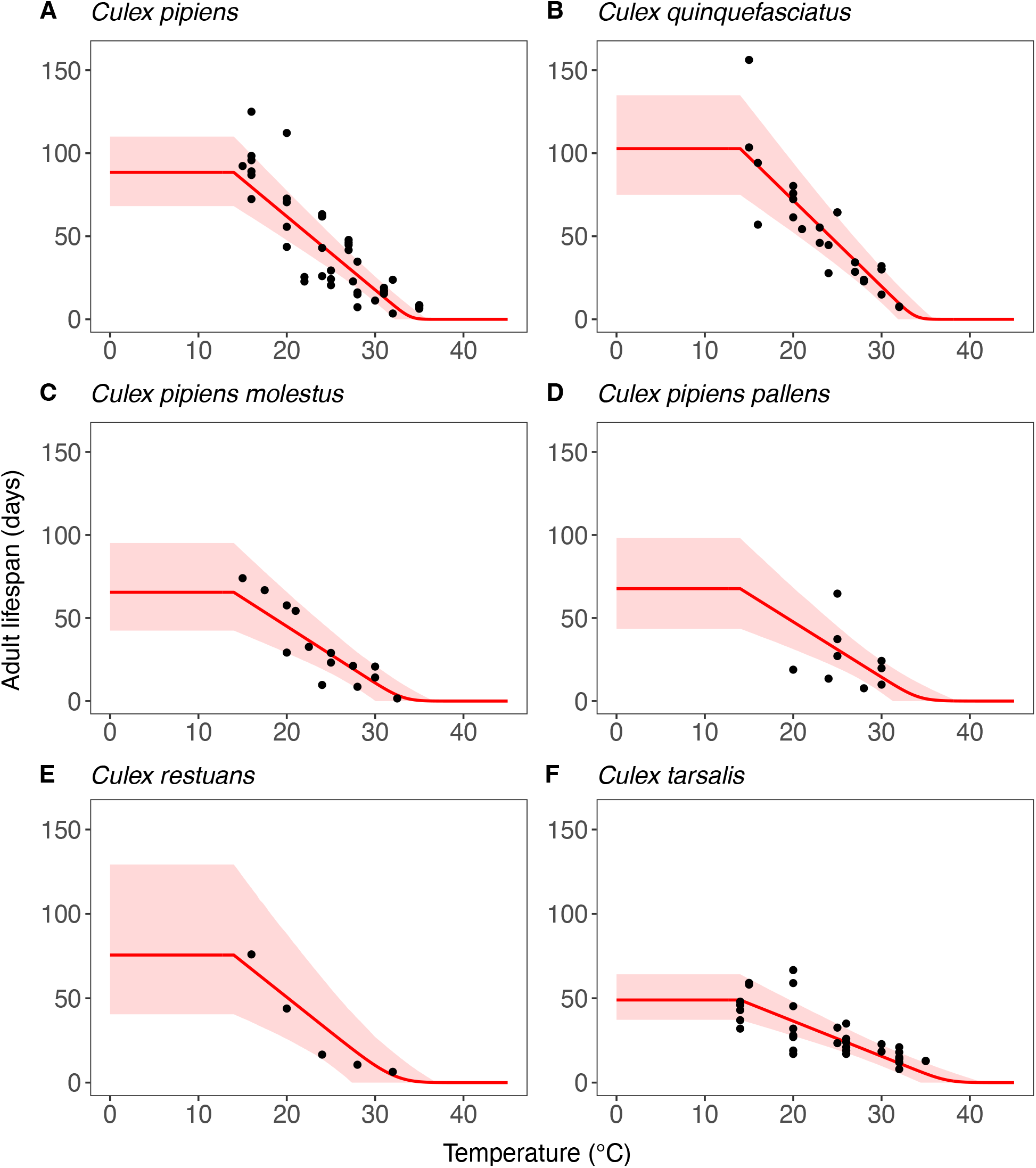
Estimates of the expected response of adult mosquito lifespan for six *Cx*. mosquitoes. Black dots represent data from experimental studies. Red solid lines represent posterior distribution mean model fits. Red shaded areas represent the corresponding central/equal-tailed 95% credible interval.

**Figure 9.**
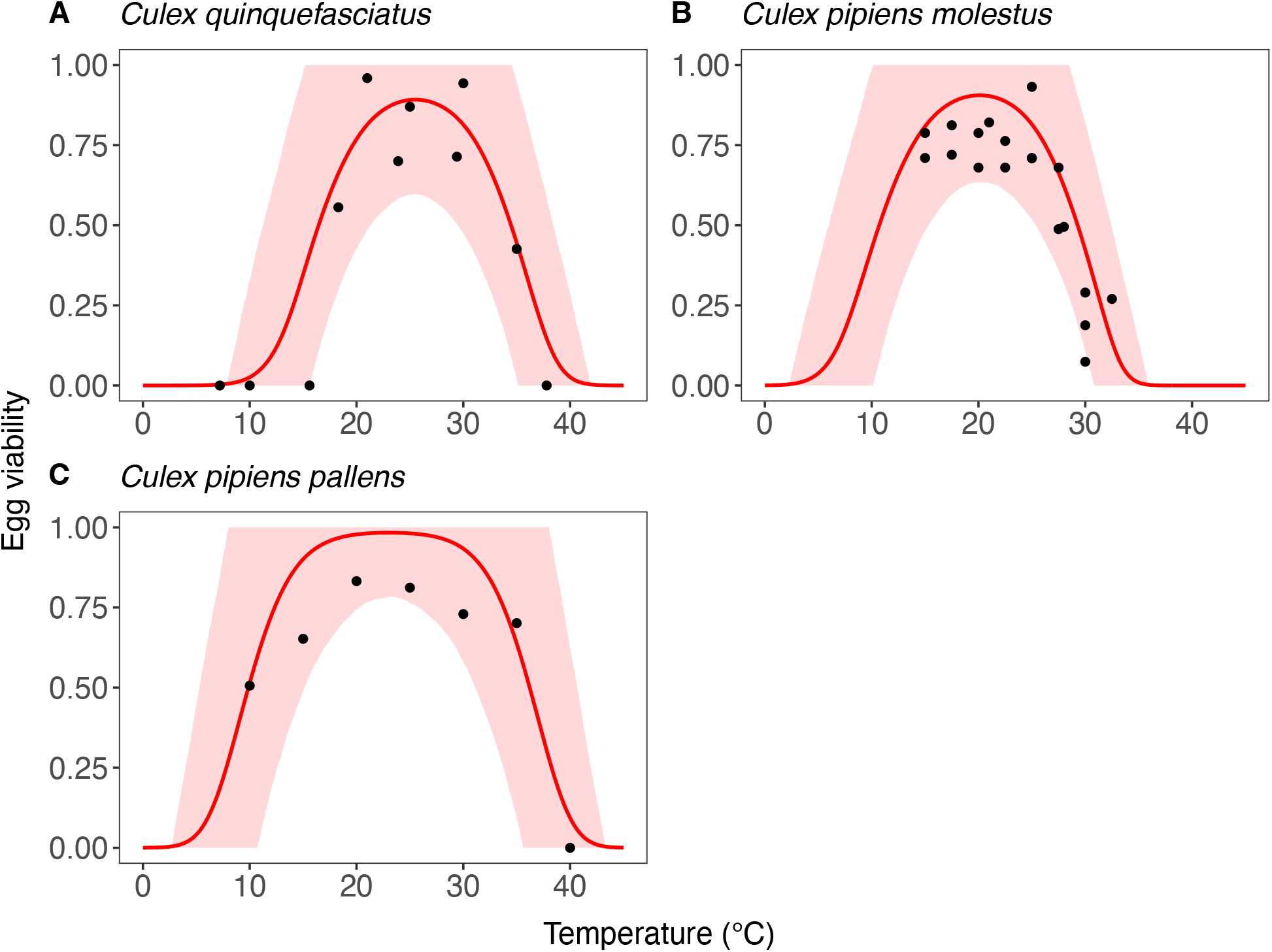
Estimates of the expected temperature response of egg viability for three *Cx*. mosquitoes. Black dots represent data from experimental studies. Red solid lines represent posterior distribution mean model fits. Red shaded areas represent the corresponding central/equal-tailed 95% credible interval.

**Figure 10.**
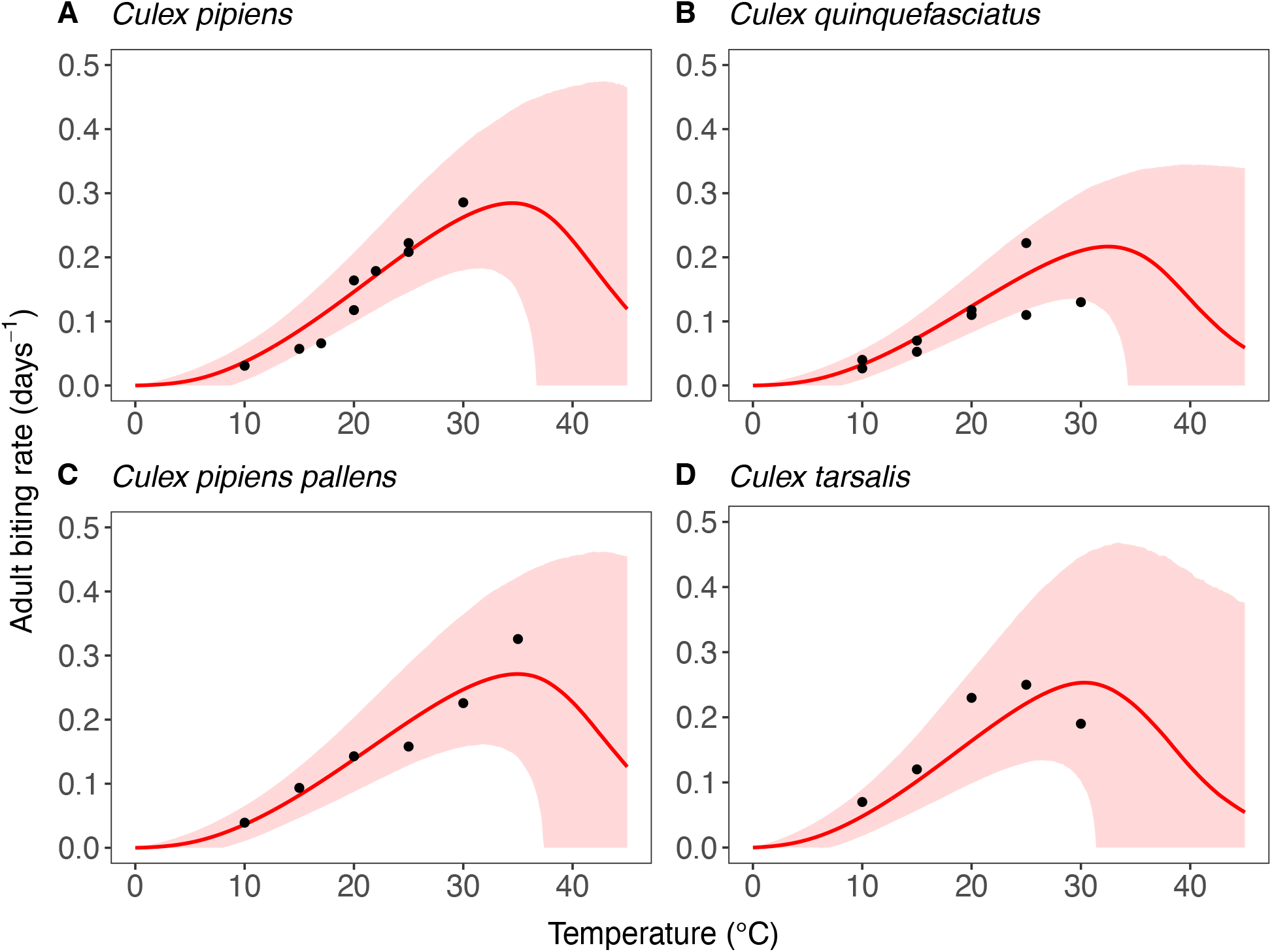
Estimates of the expected temperature response of the adult biting rate for four *Cx*. mosquitoes. Black dots represent data from experimental studies. Red solid lines represent posterior distribution mean model fits. Red shaded areas represent the corresponding central/equal-tailed 95% credible interval.

For the juvenile development rate, the estimate of the between-experiment standard deviation for the temperature limit parameters of the Briére function was larger than the between-species standard deviation. Conversely, the between-species standard deviation was larger for the parameter *q* of the same function. In contrast, for juvenile survival, we estimated a higher variability in the temperature limit parameters of the quadratic function between species than between experiments, and the opposite was true for the parameter *q*. For the slope parameter of the linear function for mosquito adult lifespan, we estimated a greater variability across species than between experiments. Despite these differences, the 95% CIs for between-species and between-experiment standard deviations largely overlapped for all three traits. The estimates of all other between-species and between-experiment variabilities showed similar patterns. However, due to data scarcity, these estimates were derived using informative hyperpriors to regularize parameter estimation.

### Temperature response of pathogen-related traits

Figures 11 and 12 show the estimated temperature responses of the mosquito infection probability and extrinsic incubation period, respectively. Posterior mean parameter estimates and 95% CIs for the pathogen-related traits can be found in Tables SI2.7-8 in S1 Text. For mosquito infection probability, only data for two WNV strains in *Cx. pipiens* was available. The sigmoidal model described well the monotonic increase of this trait with temperature seen in the data. For the extrinsic incubation period, data was available for three different WNV strains tested in *Cx. pipiens, Cx. univittatus*, or *Cx. tarsalis*. The experiment level estimates shown in Figure 12E indicate differences in the extrinsic incubation period across the different experiments. The experiment on WNV H442 in *Cx. univittatus* is estimated to have the shortest extrinsic incubation period across all temperature settings, followed by WNV NY99 in *Cx. tarsalis*, WNV WN02 in *Cx. pipiens*, and WNV NY99 in *Cx. pipiens*. Our exponentially decreasing model for the extrinsic incubation period was generally well in line with the available data. Only in the case of WNV H442 in *Cx. univittatus* the data suggested an increase of the extrinsic incubation period at high temperatures which resulted in a poor model fit for the 30°C setting for this experiment (Figure 12C). Due to the limited data available, we fixed the between-experiment standard deviation of parameters for these traits and could not estimate it. As a result, the population-level temperature responses generated for these traits (see Figure SI4.2 in S1 Text) should be seen as provisional. We investigate the sensitivity of our results to this data gap in SI4 in S1 Text.

**Figure 11.**
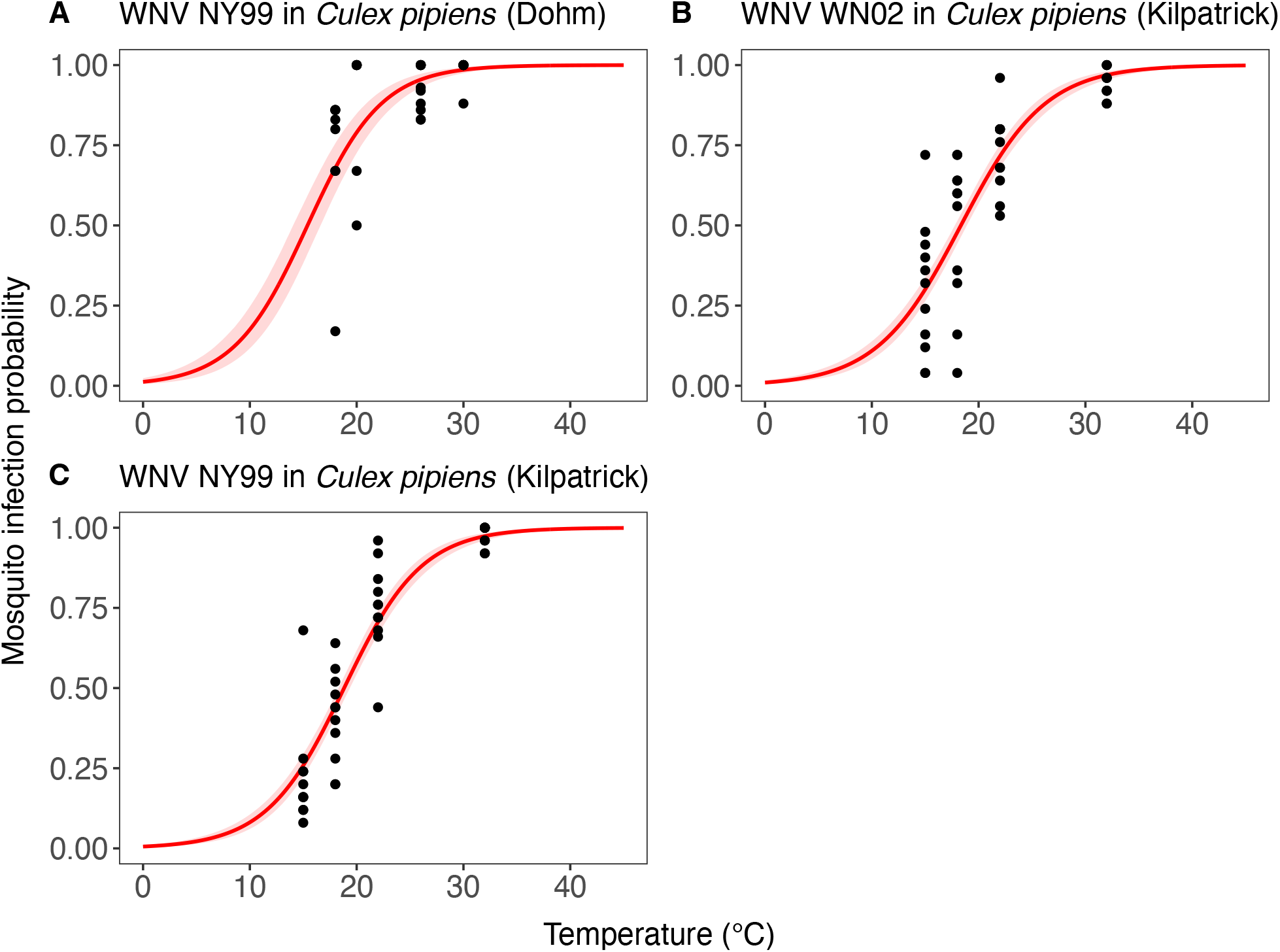
Estimates of the expected temperature response of the mosquito infection probability for three experiments testing different WNV strains in *Cx. pipiens*. Black dots represent data from experimental studies. Red solid lines represent posterior distribution mean model fits. Red shaded areas represent the corresponding central/equal-tailed 95% credible interval.

**Figure 12.**
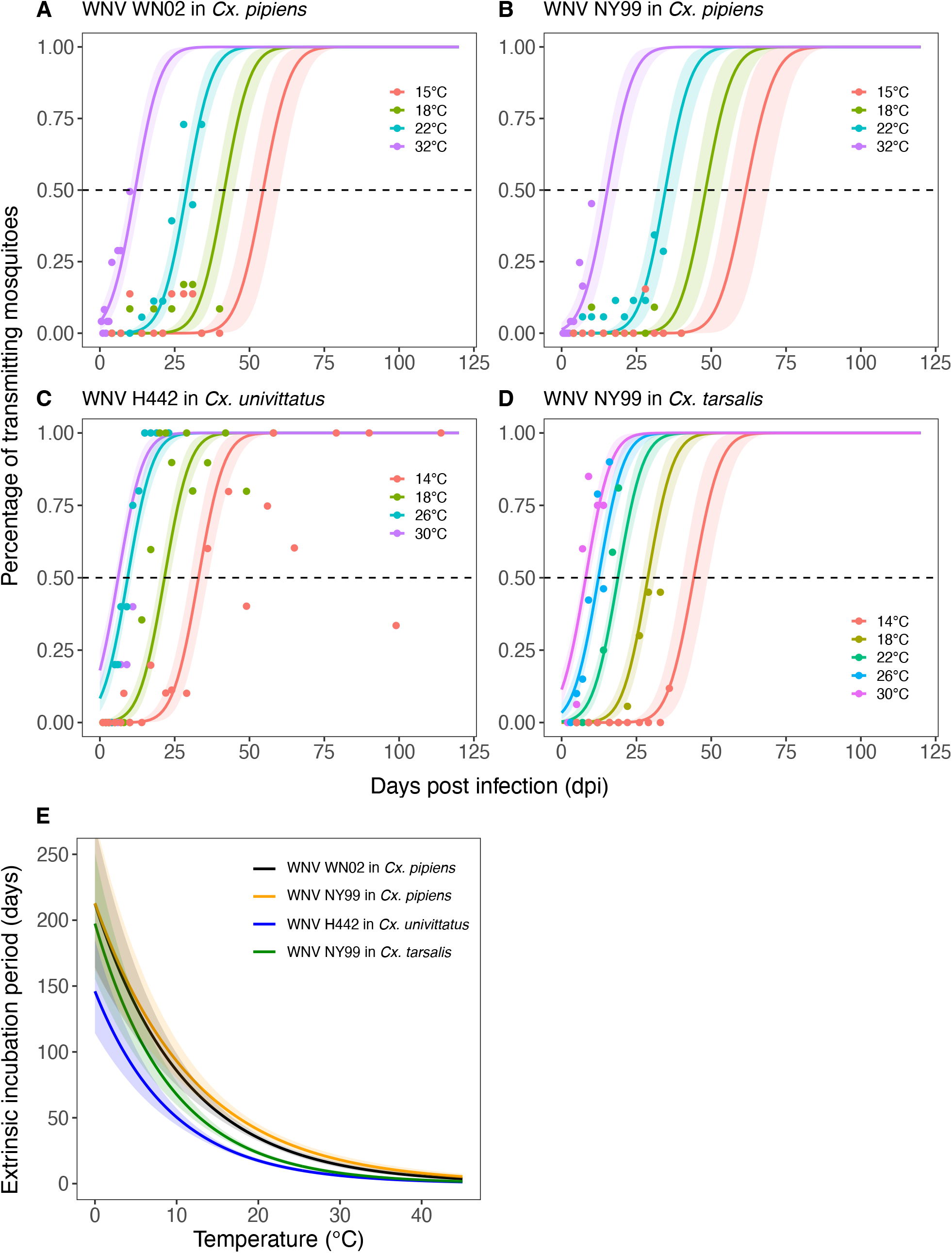
(A-D) Estimated relationship between the days post-infection and the percentage of transmitting mosquitoes upon infected mosquitoes for four different experiments each testing multiple temperatures. (E) Corresponding estimates of the expected temperature response of the extrinsic incubation period (in our statistical model defined as the time when 50% transmitting mosquitoes are expected to be reached, see Equation (16)) for the four different experiments. Solid lines represent posterior distribution mean model fits. Shaded areas represent the corresponding central/equal-tailed 95% credible interval.

## Discussion

### Research in context and novel contributions

This study provides novel estimates for the temperature response of WNV transmission suitability using a relative version of the basic reproduction number 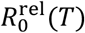. We constructed Bayesian hierarchical models to estimate species-specific temperature response functions for several mosquito-pathogen traits influencing transmission suitability. For this novel multi-species analysis, we updated a dataset compiling outcomes of laboratory studies that test trait responses to temperature. While the dataset includes various other mosquito species that help to estimate hierarchical priors and regularize model fits, our analysis focuses on *Cx*. mosquitoes and their potential to transmit WNV.

Our results suggest that WNV transmission suitability is optimized around 24°C with striking similarities between all species considered. These results are in line with some previous estimates of WNV temperature suitability [15,20,24–26] but notably lower than others [27,90]. We estimated the highest lower temperature limit for *Cx. quinquefasciatus* which could be explained by the more tropical distribution range of this species. However, we estimate two other species, *Cx. pipiens pallens* and *Cx. tarsalis*, to have a higher temperature optimum and upper temperature limit. In addition, the 95% credible intervals of temperature limits and the temperature optimum of all species were largely overlapping. We substituted a species’ trait temperature response with a population-level estimate when species-specific trait data was lacking. This led to relatively large uncertainties in the temperature limits and temperature optimum estimates for some species. Therefore, while our analysis suggests that all six considered WNV vectors respond similarly to temperature, the data currently available are insufficient to detect statistically robust differences between species.

A direct comparison between temperature limits estimated in our study and previous studies on WNV temperature suitability is difficult due to conceptual differences in the modelling approaches and in trait definitions. Our 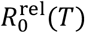 model is zero if and only if the mosquito population equilibrium *M*^∗^(*T*) is zero, determined by the mosquito life-history traits. Therefore, the temperature limits in our study have a clear interpretation as limits for viable mosquito populations. This contrasts with previous studies [15,20,25,26] where pathogen-related traits were not modelled as strictly positive and could therefore affect the temperature limits of 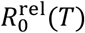.

We integrated the impact of temperature on mosquito abundance into the transmission suitability model using an equilibrium expression derived from a mosquito population dynamics model. Several previous works on mosquito thermal biology have used a similar approach but based their mosquito abundance expression on an erroneous equilibrium derivation (see SI7 in S1 Text) [19,20,22,23,25,26,28–30,32,61]. We compared our result on temperature suitability for WNV in *Cx. pipiens* to a model that uses this previous expression and to equilibria derived from alternative mosquito population dynamic models that differ in their representation of intra-specific competition [55–60]. We showed that the essential estimate of the optimal temperature for transmission suitability can vary by over 3°C across these models. While this variation does not question that WNV transmission suitability peaks at moderate temperatures, a 3°C difference is in the range of mean temperature changes by the end of century in suboptimal climate change scenarios and should therefore heavily impact WNV risk change projections. The primary model introduced in the main text draws on standard model formulations in mosquito ecology but is still a highly simplified representation of reality. For this reason, we cannot conclusively assert that our predictions for the optimum temperature are the most accurate. Currently, we emphasize the need to further investigate this sensitivity, to transparently outline the assumptions and mechanisms underlying modelling choices, and to scrutinize these models.

For a multi-species comparison, as considered in this work, the Bayesian hierarchical models introduced here have important advantages for parameter estimation over the statistical models used in previous works on the thermal biology of mosquito-borne diseases [20,22,29]. The usage of hierarchical priors offers a data-driven approach to introduce partial pooling into the species-level (and experiment-level) trait temperature response estimates which helps constrain model fits in cases with sparse data. In addition, by sampling from the hierarchical, we can generate uncertainty estimates for the temperature response of mosquito species that lack trait data. Earlier works [20,22,29] instead constructed empirical priors from a pooled dataset on other mosquito species. In contrast to the hierarchical priors introduced here, these priors lack a direct interpretation. In addition, since derived from a pooled dataset, they can act overly restrictive on the posterior estimates. Therefore, to regulate the influence of these priors, their variance was inflated in some cases before usage in the final Bayesian model fitting. However, the choice of variance scaling factor for each trait and species is subjective, hard to reproduce given new data, and its impact on the final parameter estimates is untransparent. Moreover, previous studies integrating data from multiple different experiments on the same species often ignored to account for between-experiment variability in the temperature response estimates [20,22,23,25,29]. This leads to omission of dependencies in the data and, ultimately, causes biased and overconfident model fits as illustrated in SI5 in S1 Text. The new approach introduced here solves these issues.

For a few life-history traits (juvenile development rate, juvenile survival, and partly for mosquito adult lifespan), we had sufficient data to estimate the between-species and between-experiment variability of parameter estimates using vague hyperpriors. Our results indicate that trait temperature responses exhibit similar variability across species as across experiments on the same species. While we observe potential differences in the relative importance of these variabilities among traits, further data are needed to confirm these findings. Variability in the trait temperature response across experiments on the same species may, for instance, arise due to unaccounted differences in experimental setups (such as differences in food supply for juvenile development), intraspecific genetic variation, or phenotypic plasticity. Whatever the reason, this variability suggests that outcomes from single experiments can misrepresent the expected temperature response of a species and ultimately hindering comparisons between species. Moreover, these findings emphasise that if we are interested in the response of a very specific field population, our temperature response estimates may well serve as a valuable prior assumption but could probably be refined by experimentation on the field population. A recent study [26] looking at two distinct *Cx. pipiens* populations from New York State showed variation in life history trait temperature responses between the populations consistent with local temperature conditions and WNV prevalence. This further underlines the potential and importance of intraspecific temperature response variation.

Data for the other life-history traits (egg viability, egg development rate, adult biting rate, and partly adult lifespan) was too scarce to estimate the full hierarchical model without including prior knowledge. In these cases, we defined informative hyperpriors on the between-species and between-experiment standard deviations. We assigned these hyperpriors drawing on either the fit for juvenile development rates or juvenile survival according to the biological similarity between traits. We considered this modelling step necessary to regularize the temperature response fits by implying realistic variabilities of parameter estimates across species and experiments. For the pathogen-related traits, mosquito infection probability and extrinsic incubation period, data was even scarcer, and we defined a less complex hierarchical model that only estimates parameters for each individual experiment. Here, we fixed the between-experiment standard deviation of parameters to a conservative value and investigated the sensitivity of our results to this subjective choice (see SI4 in S1 Text).

Additional experimental data on mosquito adult lifespan, adult biting rate, egg viability, egg development rate, and pathogen-related traits are thus needed to allow less restrictive modelling assumptions and derive more accurate estimates of how these traits vary across mosquito species and WNV strains. For life-history traits, we suggest focusing experimental studies on trait and species combinations that currently lack any data, such as *Cx. pipiens, Cx. tarsalis*, and *Cx. restuans* for egg viability or *Cx. pipiens molestus* and *Cx. restuans* for biting rate. For the pathogen-related traits, experiments should optimally not only test multiple temperature settings but also sample across at least three different time points post-infection to allow estimation of sophisticated temperature response functions. For example, none of the vector competence experiments in our dataset was conducted using mosquito populations or WNV strains of European origin (see Table SI9.1 in S1 Text), despite the importance of WNV in Europe. Several WNV vector competence studies have been conducted with European species and strains but to the best of our knowledge all of them either test less than three temperatures or sample only once or twice post-infection [91–97] challenging the use of their results for mathematical modelling. In addition to collecting more data on the *Cx*. species that our study focuses on, enriching the dataset with additional trait data from other (temperate) species could also improve the estimation of hierarchical priors.

### Limitations and methodological considerations

Our analysis has several limitations. We use the relative basic reproduction number as a static measure of transmission suitability under constant temperatures. This model captures the nonlinear effects of constant temperatures on the different mosquito-pathogen traits and detects optimal temperature conditions for disease transmission assuming a static environment. Similar approaches have been widely adopted to different mosquito-pathogen systems and successfully applied to predict general patterns of disease occurrence [19,20,22,25,25,26]. Nonetheless, this approach might fail to capture additional nonlinear effects created by temperature fluctuations which naturally occur in space and time. This raises the question how applicable our results and models are to real-world scenarios, both to predict individual mosquito-pathogen trait performance and transmission suitability. This question is subject to current research in insect thermal biology and has not yet been conclusively clarified [19,31,98]. As a result of Jensen’s inequality and of the nonlinear temperature response of mosquito-pathogen traits and transmission suitability, estimates for the same mean temperature are expected to alter for different forms of temperature fluctuations around the mean [31,99,100]. At the trait level, this difference can directly be predicted from our temperature response estimates by rate summation applied to the instantaneous rates that are directly (e.g., development rates) or indirectly (e.g., mortality rates derived from survival probabilities) connected to the mosquito-pathogen traits. If rate summation fails to predict trait performance under fluctuating conditions, fluctuating temperatures would have additional influences on trait performance through mechanisms such as stress accumulation, temperature acclimation, repair mechanisms during exposures to favourable temperatures, or ontogenetic shifts [98]. There is evidence that insect development rates, for example, can be accurately predicted via rate summation from experimental data generated under constant temperature conditions [98], while the same might not be true for survival, where constant temperature exposures might fail to capture time-dependent, non-lethal effects from which individuals might recover when returned to more favourable conditions [19,98,101,102]. A comprehensive analysis under which conditions rate summation can accurately predict mosquito-pathogen trait performance is currently lacking [20]. We want to note that using trait performance data from laboratory studies conducted under fluctuating instead of constant temperatures to fit temperature response functions [28] makes rate summation and therefore prediction of other fluctuation scenarios inapplicable. To relax assumptions of constant temperature in the transmission suitability model, analytical approaches developed under assumptions of periodic environments could be applied in future works [103,104]. Furthermore, our mosquito-pathogen trait response functions could also be used to drive simulations of dynamic models. This would allow to investigate the impact of any form of temperature fluctuations on mosquito population and disease dynamics given fine-resolution temperature input data, relying on the assumption that rate summation offers accurate predictions. In contrast to these more complex approaches the 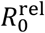 model probably has less explanatory power but represents a general model to compare temperature suitability across mosquito-pathogen systems [19].

Laboratory estimates of mosquito-pathogen traits can differ significantly from field performance. For example, it is well known that adult mosquito longevity is substantially shorter in the field than under laboratory conditions [105,106]. Similarly, the use of lab-colonized mosquito populations, partly applying to our dataset (see Table SI9.1 in S1 Text), can impact trait performance. This is a well-known issue for vector competence estimates [93,107,108] where field estimates often show a complicated picture of geographic and temporal variation [108,109].

Data across the experimental studies gathered in our database is reported quite heterogeneously. While some data is available as numbers in tables or text, a significant amount of data had to be extracted from figures using a plot digitizer tool [85]. Moreover, data either represent summary statistics across outcomes of multiple replicates (e.g., [46,69,78]) or are the outcome of a single replicate (e.g., [36,81]). The number of replicates, the sample size within each replicate, and measures of uncertainty were often not fully reported. These data limitations restricted our modelling possibilities. For instance, we approximated juvenile survival and egg viability using a normal distribution likelihood. A binomial distribution likelihood would be a more suitable model for these traits but would require access to raw data (sample size and number of successes) as we had available for the mosquito infection probability. Overall, more standardised data reporting across mosquito temperature response experiments would be highly valuable for the subsequent use of this data for modelling and the synthesis of results across studies.

Our analysis is confined to the effect of constant temperature on mosquito-pathogen traits and WNV transmission suitability. While our results can be used to gain important insights into potential shifts in distribution and seasonality of WNV under climate change, the *Cx*. life cycle and transmission of WNV depend on a wide range of other factors like precipitation, humidity, land cover, socio-demographic variables, as well as host abundance, diversity, phenology, and immunity amongst others [17]. A first step to enable the extension of our models beyond temperature would be to generate extensive data using multifactorial experiments. Bayesian hierarchical models could then be used to fit temperature response surfaces that show how trait performance varies across temperature and an additional interacting factor such as humidity [110]. These trait performance surfaces could then be incorporated into transmission dynamic and suitability models to allow better predictions of transmission risk in time and space.

## Conclusion

Understanding how mosquito-borne disease transmission responds to changes in climatic conditions is critical for proactive climate change adaptation. Our novel mosquito-pathogen trait temperature response fits can be used to parameterize mathematical models of mosquito population and disease transmission dynamics. Furthermore, our temperature-driven transmission suitability models can be used to monitor general patterns and trends in WNV suitability. Additional experimental studies are needed to further enhance the robustness of our approach. These should focus on traits currently presenting the least data, i.e., adult biting rate, egg viability, mosquito adult lifespan, egg development rate, and vector competence studies. To increase the value of these experiments for mathematical modelling they should optimally test multiple temperatures over a wide range of values. Furthermore, openly sharing raw data would give modellers the possibility to use the most suitable statistical approach for fitting each trait. Given extended databases, future studies could extend our analyses to more species, consider other and more detailed hierarchies in the statistical model, or extend our models to additional factors beyond temperature.

## Acknowledgements

The authors want to thank Dionysis Terzis for supporting their data collection. Further thanks go to Jordan Ruybal, Marm Kilpatrick, and William Reisen for sharing raw data to support the analysis.

## Data and code availability

All data and code supporting this study are available on GitHub: https://github.com/julehe/WNV-temperature

## Supporting information

**S1 Text**. SI1, Details on functions fitted to mosquito-pathogen traits. SI2, Summary of mosquito-pathogen temperature response fits. SI3, Formal mathematical description of Bayesian hierarchical models and (hyper)prior specifications. SI4, Population-level trait estimates and sensitivity to between-experiment variability. SI5, Importance of accounting for between-experiment variability. SI6, Taxonomic considerations. SI7, Alternative mosquito abundance models. SI8, Key changes applied to original dataset. SI9, Overview of collected data.

## References

1. Petersen LR, Brault AC, Nasci RS. West Nile Virus: Review of the Literature. JAMA. 2013;310: 308. doi:10.1001/jama.2013.8042

2. Habarugira G, Suen WW, Hobson-Peters J, Hall RA, Bielefeldt-Ohmann H. West Nile Virus: An Update on Pathobiology, Epidemiology, Diagnostics, Control and “One Health” Implications. Pathogens. 2020;9: 589. doi:10.3390/pathogens9070589

3. Farajollahi A, Fonseca DM, Kramer LD, Marm Kilpatrick A. “Bird biting” mosquitoes and human disease: A review of the role of Culex pipiens complex mosquitoes in epidemiology. Infection, Genetics and Evolution. 2011;11: 1577–1585. doi:10.1016/j.meegid.2011.08.013

4. Goldberg TL, Ruiz MO, Hamer GL, Brawn JD, Kitron UD, Hayes DB, et al. Host Selection by Culex pipiens Mosquitoes and West Nile Virus Amplification. The American Journal of Tropical Medicine and Hygiene. 2009;80: 268–278. doi:10.4269/ajtmh.2009.80.268

5. Blom R, Krol L, Langezaal M, Schrama M, Trimbos KB, Wassenaar D, et al. Blood-feeding patterns of Culex pipiens biotype pipiens and pipiens/molestus hybrids in relation to avian community composition in urban habitats. Parasites Vectors. 2024;17: 95. doi:10.1186/s13071-024-06186-9

6. Saiz J-C. Animal and Human Vaccines against West Nile Virus. Pathogens. 2020;9: 1073. doi:10.3390/pathogens9121073

7. Ulbert S. West Nile virus vaccines – current situation and future directions. Human Vaccines & Immunotherapeutics. 2019;15: 2337–2342. doi:10.1080/21645515.2019.1621149

8. Gray T, Webb CE. A review of the epidemiological and clinical aspects of West Nile virus. IJGM. 2014; 193. doi:10.2147/IJGM.S59902

9. Kramer LD, Ciota AT, Kilpatrick AM. Introduction, Spread, and Establishment of West Nile Virus in the Americas. Reisen W, editor. Journal of Medical Entomology. 2019;56: 1448–1455. doi:10.1093/jme/tjz151

10. Epidemiological update: West Nile virus transmission season in Europe, 2018. 14 Dec 2018 [cited 5 Mar 2023]. Available: https://www.ecdc.europa.eu/en/news-events/epidemiological-update-west-nile-virus-transmission-season-europe-2018

11. Romanello M, Di Napoli C, Drummond P, Green C, Kennard H, Lampard P, et al. The 2022 report of the Lancet Countdown on health and climate change: health at the mercy of fossil fuels. The Lancet. 2022;400: 1619–1654. doi:10.1016/S0140-6736(22)01540-9

12. Erazo D, Grant L, Ghisbain G, Marini G, Colón-González FJ, Wint W, et al. Contribution of climate change to the spatial expansion of West Nile virus in Europe. Nat Commun. 2024;15: 1196. doi:10.1038/s41467-024-45290-3

13. van Daalen KR, Romanello M, Rocklöv J, Semenza JC, Tonne C, Markandya A, et al. The 2022 Europe report of the Lancet Countdown on health and climate change: towards a climate resilient future. The Lancet Public Health. 2022; S2468266722001979. doi:10.1016/S2468-2667(22)00197-9

14. Farooq Z, Sjödin H, Semenza JC, Tozan Y, Sewe MO, Wallin J, et al. European projections of West Nile virus transmission under climate change scenarios. One Health. 2023;16: 100509. doi:10.1016/j.onehlt.2023.100509

15. Paull SH, Horton DE, Ashfaq M, Rastogi D, Kramer LD, Diffenbaugh NS, et al. Drought and immunity determine the intensity of West Nile virus epidemics and climate change impacts. Proc R Soc B. 2017;284: 20162078. doi:10.1098/rspb.2016.2078

16. Ewing DA, Purse BV, Cobbold CA, White SM. A novel approach for predicting risk of vector-borne disease establishment in marginal temperate environments under climate change: West Nile virus in the UK. J R Soc Interface. 2021;18: 20210049. doi:10.1098/rsif.2021.0049

17. Heidecke J, Lavarello Schettini A, Rocklöv J. West Nile virus eco-epidemiology and climate change. Paz S, editor. PLOS Clim. 2023;2: e0000129. doi:10.1371/journal.pclm.0000129

18. Rocklöv J, Dubrow R. Climate change: an enduring challenge for vector-borne disease prevention and control. Nat Immunol. 2020;21: 479–483. doi:10.1038/s41590-020-0648-y

19. Mordecai EA, Caldwell JM, Grossman MK, Lippi CA, Johnson LR, Neira M, et al. Thermal biology of mosquito-borne disease. Byers J (Jeb), editor. Ecol Lett. 2019;22: 1690–1708. doi:10.1111/ele.13335

20. Shocket MS, Verwillow AB, Numazu MG, Slamani H, Cohen JM, El Moustaid F, et al. Transmission of West Nile and five other temperate mosquito-borne viruses peaks at temperatures between 23°C and 26°C. eLife. 2020;9: e58511. doi:10.7554/eLife.58511

21. Liu-Helmersson J, Stenlund H, Wilder-Smith A, Rocklöv J. Vectorial Capacity of Aedes aegypti: Effects of Temperature and Implications for Global Dengue Epidemic Potential. Moreira LA, editor. PLoS ONE. 2014;9: e89783. doi:10.1371/journal.pone.0089783

22. Mordecai EA, Cohen JM, Evans MV, Gudapati P, Johnson LR, Lippi CA, et al. Detecting the impact of temperature on transmission of Zika, dengue, and chikungunya using mechanistic models. Althouse B, editor. PLoS Negl Trop Dis. 2017;11: e0005568. doi:10.1371/journal.pntd.0005568

23. Villena OC, Ryan SJ, Murdock CC, Johnson LR. Temperature impacts the environmental suitability for malaria transmission by Anopheles gambiae and Anopheles stephensi. Ecology. 2022;103. doi:10.1002/ecy.3685

24. McMillan JR, Blakney RA, Mead DG, Koval WT, Coker SM, Waller LA, et al. Linking the vectorial capacity of multiple vectors to observed patterns of West Nile virus transmission. Nuñez M, editor. Journal of Applied Ecology. 2019;56: 956–965. doi:10.1111/1365-2664.13322

25. Di Pol G, Crotta M, Taylor RA. Modelling the temperature suitability for the risk of West Nile Virus establishment in European Culex pipiens populations. Transbounding Emerging Dis. 2022;69. doi:10.1111/tbed.14513

26. Fay RL, Cruz-Loya M, Keyel AC, Price DC, Zink SD, Mordecai EA, et al. Population-specific thermal responses contribute to regional variability in arbovirus transmission with changing climates. iScience. 2024;27: 109934. doi:10.1016/j.isci.2024.109934

27. Vogels CBF, Hartemink N, Koenraadt CJM. Modelling West Nile virus transmission risk in Europe: effect of temperature and mosquito biotypes on the basic reproduction number. Sci Rep. 2017;7: 5022. doi:10.1038/s41598-017-05185-4

28. Moser SK, Barnard M, Frantz RM, Spencer JA, Rodarte KA, Crooker IK, et al. Scoping review of Culex mosquito life history trait heterogeneity in response to temperature. Parasites Vectors. 2023;16: 200. doi:10.1186/s13071-023-05792-3

29. Johnson LR, Ben-Horin T, Lafferty KD, McNally A, Mordecai E, Paaijmans KP, et al. Understanding uncertainty in temperature effects on vector-borne disease: a Bayesian approach. Ecology. 2015;96: 203–213. doi:10.1890/13-1964.1

30. Shocket MS, Ryan SJ, Mordecai EA. Temperature explains broad patterns of Ross River virus transmission. eLife. 2018;7: e37762. doi:10.7554/eLife.37762

31. Lambrechts L, Paaijmans KP, Fansiri T, Carrington LB, Kramer LD, Thomas MB, et al. Impact of daily temperature fluctuations on dengue virus transmission by Aedes aegypti. Proc Natl Acad Sci USA. 2011;108: 7460–7465. doi:10.1073/pnas.1101377108

32. Mordecai EA, Paaijmans KP, Johnson LR, Balzer C, Ben-Horin T, de Moor E, et al. Optimal temperature for malaria transmission is dramatically lower than previously predicted. Thrall P, editor. Ecol Lett. 2013;16: 22–30. doi:10.1111/ele.12015

33. Dietz K. The estimation of the basic reproduction number for infectious diseases. Stat Methods Med Res. 1993;2: 23–41. doi:10.1177/096228029300200103

34. Smith DL, Battle KE, Hay SI, Barker CM, Scott TW, McKenzie FE. Ross, Macdonald, and a Theory for the Dynamics and Control of Mosquito-Transmitted Pathogens. Chitnis CE, editor. PLoS Pathog. 2012;8: e1002588. doi:10.1371/journal.ppat.1002588

35. Spanoudis CG, Andreadis SS, Tsaknis NK, Petrou AP, Gkeka CD, Savopoulou–Soultani M. Effect of Temperature on Biological Parameters of the West Nile Virus Vector Culex pipiens form ‘molestus’ (Diptera: Culicidae) in Greece: Constant vs Fluctuating Temperatures. Journal of Medical Entomology. 2019;56: 641–650. doi:10.1093/jme/tjy224

36. Loetti V, Schweigmann N, Burroni N. Development rates, larval survivorship and wing length of Culex pipiens (Diptera: Culicidae) at constant temperatures. Journal of Natural History. 2011;45: 2203–2213. doi:10.1080/00222933.2011.590946

37. Gunay F, Alten B, Ozsoy ED. Estimating reaction norms for predictive population parameters, age specific mortality, and mean longevity in temperature-dependent cohorts of Culex quinquefasciatus Say (Diptera: Culicidae). Journal of Vector Ecology. 2010;35: 354–362. doi:10.1111/j.1948-7134.2010.00094.x

38. Reisen WK. Effect of Temperature on Culex tarsalis (Diptera: Culicidae) from the Coachella and San Joaquin Valleys of California. Journal of Medical Entomology. 1995;32: 636–645. doi:10.1093/jmedent/32.5.636

39. Mogi M. Temperature and Photoperiod Effects on Larval and Ovarian Development of New Zealand Strains of Culex quinquefasciatus (Diptera: Culicidae). Annals of the Entomological Society of America. 1992;85: 58–66. doi:10.1093/aesa/85.1.58

40. Tekle A. The Physiology of Hibernation and Its Role in the Geographical Distribution of Populations of the Culex Pipiens Complex. The American Journal of Tropical Medicine and Hygiene. 1960;9: 321–330. doi:10.4269/ajtmh.1960.9.321

41. Madder DJ, Surgeoner GA, Helson BV. Number of Generations, Egg Production, and Developmental Time of Culex Pipiens and Culex Restuans (Diptera: Culicidae) in Southern Ontario1. Journal of Medical Entomology. 1983;20: 275–287. doi:10.1093/jmedent/20.3.275

42. Buth JL, Brust RA, Ellis RA. Development time, oviposition activity and onset of diapause in Culex tarsalis, Culex restuans and Culiseta inornata in southern Manitoba. J Am Mosq Control Assoc. 1990;6: 55–63.

43. Teng H-J, Apperson CS. Development and Survival of Immature Aedes albopictus and Aedes triseriatus (Diptera: Culicidae) in the Laboratory: Effects of Density, Food, and Competition on Response to Temperature. J Med Entomol. 2000;37: 40–52. doi:10.1603/0022-2585-37.1.40

44. Ciota AT, Matacchiero AC, Kilpatrick AM, Kramer LD. The Effect of Temperature on Life History Traits of Culex Mosquitoes. J Med Entomol. 2014;51: 55–62. doi:10.1603/ME13003

45. Mpho M, Callaghan A, Holloway GJ. Temperature and genotypic effects on life history and fluctuating asymmetry in a field strain of Culex pipiens. Heredity. 2002;88: 307–312. doi:10.1038/sj.hdy.6800045

46. Ruybal JE, Kramer LD, Kilpatrick AM. Geographic variation in the response of Culex pipiens life history traits to temperature. Parasites Vectors. 2016;9: 116. doi:10.1186/s13071-016-1402-z

47. Muturi EJ, Lampman R, Costanzo K, Alto BW. Effect of Temperature and Insecticide Stress on Life-History Traits of Culex restuans and Aedes albopictus (Diptera: Culicidae). jnl med entom. 2011;48: 243–250. doi:10.1603/ME10017

48. Olejnícek J, Gelbic I. Differences in response to temperature and density between two strains of the mosquito, Culex pipiens molestus forskal. J Vector Ecol. 2000;25: 136–145.

49. Dodson BL, Kramer LD, Rasgon JL. Effects of larval rearing temperature on immature development and West Nile virus vector competence of Culex tarsalis. Parasites Vectors. 2012;5: 199. doi:10.1186/1756-3305-5-199

50. Brust RA. Weight And Development Time Of Different Stadia Of Mosquitoes Reared At Various Constant Temperatures. Can Entomol. 1967;99: 986–993. doi:10.4039/Ent99986-9

51. Boerlijst SP, Johnston ES, Ummels A, Krol L, Boelee E, Van Bodegom PM, et al. Biting the hand that feeds: Anthropogenic drivers interactively make mosquitoes thrive. Science of The Total Environment. 2023;858: 159716. doi:10.1016/j.scitotenv.2022.159716

52. Ezeakacha NF, Yee DA. The role of temperature in affecting carry-over effects and larval competition in the globally invasive mosquito Aedes albopictus. Parasites Vectors. 2019;12: 123. doi:10.1186/s13071-019-3391-1

53. Shutt DP, Goodsman DW, Martinez K, Hemez ZJL, Conrad JR, Xu C, et al. A Process-based Model with Temperature, Water, and Lab-derived Data Improves Predictions of Daily Culex pipiens/restuans Mosquito Density. Gaff H, editor. Journal of Medical Entomology. 2022;59: 1947–1959. doi:10.1093/jme/tjac127

54. Watanabe K, Fukui S, Ohta S. Population of the temperate mosquito, Culex pipiens, decreases in response to habitat climatological changes in future. GeoHealth. 2017;1: 196–210. doi:10.1002/2017GH000054

55. DiSera L, Sjödin H, Rocklöv J, Tozan Y, Súdre B, Zeller H, et al. The Mosquito, the Virus, the Climate: An Unforeseen Réunion in 2018. GeoHealth. 2020;4. doi:10.1029/2020GH000253

56. Tran A, Mangeas M, Demarchi M, Roux E, Degenne P, Haramboure M, et al. Complementarity of empirical and process-based approaches to modelling mosquito population dynamics with Aedes albopictus as an example—Application to the development of an operational mapping tool of vector populations. Touzeau S, editor. PLoS ONE. 2020;15: e0227407. doi:10.1371/journal.pone.0227407

57. Liu-Helmersson J, Brännström Å, Sewe MO, Semenza JC, Rocklöv J. Estimating Past, Present, and Future Trends in the Global Distribution and Abundance of the Arbovirus Vector Aedes aegypti Under Climate Change Scenarios. Front Public Health. 2019;7: 148. doi:10.3389/fpubh.2019.00148

58. Yang HM, Boldrini JL, Fassoni AC, Freitas LFS, Gomez MC, Lima KKB de, et al. Fitting the Incidence Data from the City of Campinas, Brazil, Based on Dengue Transmission Modellings Considering Time-Dependent Entomological Parameters. Paul R, editor. PLoS ONE. 2016;11: e0152186. doi:10.1371/journal.pone.0152186

59. Caldwell JM, LaBeaud AD, Lambin EF, Stewart-Ibarra AM, Ndenga BA, Mutuku FM, et al. Climate predicts geographic and temporal variation in mosquito-borne disease dynamics on two continents. Nat Commun. 2021;12: 1233. doi:10.1038/s41467-021-21496-7

60. Ngonghala CN, Ryan SJ, Tesla B, Demakovsky LR, Mordecai EA, Murdock CC, et al. Effects of changes in temperature on Zika dynamics and control. J R Soc Interface. 2021;18: rsif.2021.0165, 20210165. doi:10.1098/rsif.2021.0165

61. Parham PE, Michael E. Modeling the Effects of Weather and Climate Change on Malaria Transmission. Environmental Health Perspectives. 2010;118: 620–626. doi:10.1289/ehp.0901256

62. B. Zayed A. Influence of Temperature Change on the Growth and Susceptibility of the Common House Mosquito, & lt;i & gt;Culex pipiens & lt;/i & gt; in Egypt to Some Insecticides. IJEE. 2019;4: 42. doi:10.11648/j.ijee.20190402.11

63. Ciota, Keyel. The Role of Temperature in Transmission of Zoonotic Arboviruses. Viruses. 2019;11: 1013. doi:10.3390/v11111013

64. Ukubuiwe AC, Olayemi IK, Arimoro FO, Omalu ICJ, Baba BM, Ukubuiwe CC, et al. Influence of rearing-water temperature on life stages’ vector attributes, distribution and utilisation of metabolic reserves in Culex quinquefasciatus (Diptera: Culicidae): implications for disease transmission and vector control. JoBAZ. 2018;79: 32. doi:10.1186/s41936-018-0045-3

65. Olayemi IK, Victoria O, Ukubuiwe AC, Jibrin AI. Effects of Temperature Stress on Pre-imaginal Development and Adult Ptero-fitness of the Vector Mosquito, Culex quinquefasciatus (Diptera: Culicidae). jmr. 2016 [cited 9 Feb 2024]. doi:10.5376/jmr.2016.06.0014

66. Shriver D, Bickley W. The effect of temperature on the hatching of eggs of the mosquito Culex pipiens quinquefasciatus say. Mosquito News. 1964;24: 137–40.

67. Oda T, Eshita Y, Uchida K, Mine M, Kurokawa K, Ogawa Y, et al. Reproductive Activity and Survival of Culex pipiens pallens and Culex quinquefasciatus (Diptera: Culicidae) in Japan at High Temperature. J Med Entomol. 2002;39: 185–190. doi:10.1603/0022-2585-39.1.185

68. Kiarie-Makara M, Ngumbi P, Lee D-K. Effects of temperature on the growth and development of culex pipiens Complex Mosquitoes (Diptera: Culicidae). Journal of Pharmacy and Biological Sciences. 2015;10: 1–10. doi:10.9790/3008-10620110

69. Mpho M, Callaghan A, Holloway GJ. Effects of temperature and genetic stress on life history and fluctuating wing asymmetry in Culex pipiens mosquitoes. Eur J Entomol. 2002;99: 405–412. doi:10.14411/eje.2002.050

70. Oda T, Mori A, Ueda M. Effects of temperatures on the oviposition and hatching of eggs in culex pipiens molestus and culex pipiens quinquefasciatus. Tropical Medicine. 1980;22: 167–180.

71. Mahmood F, Crans WJ. A thermal heat summation model to predict the duration of the gonotrophic cycle of Culiseta melanura in nature. J Am Mosq Control Assoc. 1997;13: 92–94.

72. Kilpatrick AM, Meola MA, Moudy RM, Kramer LD. Temperature, Viral Genetics, and the Transmission of West Nile Virus by Culex pipiens Mosquitoes. Buchmeier MJ, editor. PLoS Pathog. 2008;4: e1000092. doi:10.1371/journal.ppat.1000092

73. Dohm DJ, O’Guinn ML, Turell MJ. Effect of Environmental Temperature on the Ability of Culex pipiens (Diptera: Culicidae) to Transmit West Nile Virus. J Med Entomol. 2002;39: 221–225. doi:10.1603/0022-2585-39.1.221

74. Reisen WK, Fang Y, Martinez VM. Effects of Temperature on the Transmission of West Nile Virus by Culex tarsalis (Diptera: Culicidae). J Med Entomol. 2006;43: 309–317. doi:10.1093/jmedent/43.2.309

75. Cornel AJ, Jupp PG, Blackburn NK. Environmental Temperature on the Vector Competence of Culex univittatus (Diptera: Culicidae) for West Nile Virus. Journal of Medical Entomology. 1993;30: 449–456. doi:10.1093/jmedent/30.2.449

76. Rueda LM, Patel KJ, Axtell RC, Stinner RE. Temperature-Dependent Development and Survival Rates of Culex quinquefasciatus and Aedes aegypti (Diptera: Culicidae). Journal of Medical Entomology. 1990;27: 892–898. doi:10.1093/jmedent/27.5.892

77. Shelton R. The effect of temperatures on development of eight mosquito species. Mosquito News. 1973;33: 1–12.

78. Oda T, Uchida K, Mori A, Mine M, Eshita Y, Kurokawa K, et al. Effects of high temperature on the emergence and survival of adult Culex pipiens molestus and Culex quinquefasciatus in Japan. J Am Mosq Control Assoc. 1999;15: 153–156.

79. Li J, Tang J, Zhu G, Yang M, Zhou H, Zhang M, et al. Effect of temperature on development and reproduction of three kind of mosquitoes. China Tropical Medicine. 2019;19: 939–943. doi:10.13604/j.cnki.46-1064/r.2019.10.08

80. van der Linde T de K, Hewitt P, Nel A, van der Westhuizen M. Development rates and percentage hatching of culex (Culex) theileri; Theobald (Diptera: culicidae) eggs at various constant temperatures. Journal of the Entomological Society of Southern Africa. 53: 17–26.

81. Mahmood F, Crans WJ. Effect of Temperature on the Development of Culiseta melanura (Diptera: Culicidae) and its Impact on the Amplification of Eastern Equine Encephalomyelitis Virus in Birds. Journal of Medical Entomology. 1998;35: 1007–1012. doi:10.1093/jmedent/35.6.1007

82. Andreadis SS, Dimotsiou OC, Savopoulou-Soultani M. Variation in adult longevity of Culex pipiens f. pipiens, vector of the West Nile Virus. Parasitol Res. 2014;113: 4315–4319. doi:10.1007/s00436-014-4152-x

83. Reisen WK, Milby MM, Presser SB, Hardy JL. Ecology of Mosquitoes and St. Louis Encephalitis Virus in the Los Angeles Basin of California, 1987–1990. Journal of Medical Entomology. 1992;29: 582–598. doi:10.1093/jmedent/29.4.582

84. Nayar JK. Effects of constant and fluctuating temperatures on life span of Aedes taeniorhynchus adults. Journal of Insect Physiology. 1972;18: 1303–1313. doi:10.1016/0022-1910(72)90259-4

85. Rohatgi A. WebPlotDigitizer. 2018 [cited 3 Jun 2024]. Available: https://apps.automeris.io/wpd4/

86. Amarasekare P, Savage V. A Framework for Elucidating the Temperature Dependence of Fitness. The American Naturalist. 2012;179: 178–191. doi:10.1086/663677

87. Briere J-F, Pracros P, Le Roux A-Y, Pierre J-S. A Novel Rate Model of Temperature-Dependent Development for Arthropods. Environ Entomol. 1999;28: 22–29. doi:10.1093/ee/28.1.22

88. Carpenter B, Gelman A, Hoffman MD, Lee D, Goodrich B, Betancourt M, et al. Stan : A Probabilistic Programming Language. J Stat Soft. 2017;76. doi:10.18637/jss.v076.i01

89. Stan Development Team. RStan: the R interface to Stan. Available: https://mc-stan.org/

90. Kushmaro A, Friedlander TA, Levins R. Temperature Effects on the Basic Reproductive Number (R0) Of West Nile Virus, Based On Ecological Parameters: Endemic Vs. New Emergence Regions. J Trop Dis. 2015;s1. doi:10.4172/2329-891X.1000S1-001

91. Holicki CM, Ziegler U, Răileanu C, Kampen H, Werner D, Schulz J, et al. West Nile Virus Lineage 2 Vector Competence of Indigenous Culex and Aedes Mosquitoes from Germany at Temperate Climate Conditions. Viruses. 2020;12: 561. doi:10.3390/v12050561

92. Jansen S, Heitmann A, Lühken R, Leggewie M, Helms M, Badusche M, et al. Culex torrentium: A Potent Vector for the Transmission of West Nile Virus in Central Europe. Viruses. 2019;11: 492. doi:10.3390/v11060492

93. Vogels CB, Göertz GP, Pijlman GP, Koenraadt CJ. Vector competence of European mosquitoes for West Nile virus. Emerging Microbes & Infections. 2017;6: 1–13. doi:10.1038/emi.2017.82

94. Fros JJ, Geertsema C, Vogels CB, Roosjen PP, Failloux A-B, Vlak JM, et al. West Nile Virus: High Transmission Rate in North-Western European Mosquitoes Indicates Its Epidemic Potential and Warrants Increased Surveillance. Liang S, editor. PLoS Negl Trop Dis. 2015;9: e0003956. doi:10.1371/journal.pntd.0003956

95. Vogels CBF, Fros JJ, Göertz GP, Pijlman GP, Koenraadt CJM. Vector competence of northern European Culex pipiens biotypes and hybrids for West Nile virus is differentially affected by temperature. Parasites Vectors. 2016;9: 393. doi:10.1186/s13071-016-1677-0

96. Linthout C, Martins AD, De Wit M, Delecroix C, Abbo SR, Pijlman GP, et al. The potential role of the Asian bush mosquito Aedes japonicus as spillover vector for West Nile virus in the Netherlands. Parasites Vectors. 2024;17: 262. doi:10.1186/s13071-024-06279-5

97. Wöhnke E, Vasic A, Raileanu C, Holicki CM, Tews BA, Silaghi C. Comparison of vector competence of Aedes vexans Green River and Culex pipiens biotype pipiens for West Nile virus lineages 1 and 2. Zoonoses and Public Health. 2020;67: 416–424. doi:10.1111/zph.12700

98. Von Schmalensee L, Hulda Gunnarsdóttir K, Näslund J, Gotthard K, Lehmann P. Thermal performance under constant temperatures can accurately predict insect development times across naturally variable microclimates. Ghalambor C, editor. Ecology Letters. 2021;24: 1633–1645. doi:10.1111/ele.13779

99. Bernhardt JR, Sunday JM, Thompson PL, O’Connor MI. Nonlinear averaging of thermal experience predicts population growth rates in a thermally variable environment. Proc R Soc B. 2018;285: 20181076. doi:10.1098/rspb.2018.1076

100. Shocket MS. Fluctuating temperatures have a surprising effect on disease transmission. PLoS Biol. 2023;21: e3002288. doi:10.1371/journal.pbio.3002288

101. Rezende EL, Castañeda LE, Santos M. Tolerance landscapes in thermal ecology. Fox C, editor. Functional Ecology. 2014;28: 799–809. doi:10.1111/1365-2435.12268

102. Rezende EL, Bozinovic F, Szilágyi A, Santos M. Predicting temperature mortality and selection in natural Drosophila populations. Science. 2020;369: 1242–1245. doi:10.1126/science.aba9287

103. Liu-Helmersson J, Rocklöv J, Sewe M, Brännström Å. Climate change may enable Aedes aegypti infestation in major European cities by 2100. Environmental Research. 2019;172: 693–699. doi:10.1016/j.envres.2019.02.026

104. Bacaër N, Guernaoui S. The epidemic threshold of vector-borne diseases with seasonality: The case of cutaneous leishmaniasis in Chichaoua, Morocco. J Math Biol. 2006;53: 421–436. doi:10.1007/s00285-006-0015-0

105. Marini G, Poletti P, Giacobini M, Pugliese A, Merler S, Rosà R. The Role of Climatic and Density Dependent Factors in Shaping Mosquito Population Dynamics: The Case of Culex pipiens in Northwestern Italy. Hwang J-S, editor. PLoS ONE. 2016;11: e0154018. doi:10.1371/journal.pone.0154018

106. Lambert B, North A, Godfray HCJ. A Meta-analysis of Longevity Estimates of Mosquito Vectors of Disease. Ecology; 2022 May. doi:10.1101/2022.05.30.494059

107. Gargan TP, Higbee GA, El Said S, Gad A, Bailey CL. The Effect of Laboratory Colonization on the Vector-Pathogen Interactions of Egyptian Culex pipiens and Rift Valley Fever Virus *. The American Journal of Tropical Medicine and Hygiene. 1983;32: 1154–1163. doi:10.4269/ajtmh.1983.32.1154

108. Ross QE, Yuill TM, Craig GB, Grimstad PR. Aedes Triseriatus and La Crosse Virus: Geographic Variation in Vector Susceptibility and Ability to Transmit *. The American Journal of Tropical Medicine and Hygiene. 1977;26: 990–996. doi:10.4269/ajtmh.1977.26.990

109. Kilpatrick AM, Ebel GD, Reddy MR, Fonseca DM, Kramer LD. Spatial and Temporal Variation in Vector Competence of Culex pipiens and Cx. restuans Mosquitoes for West Nile Virus. The American Journal of Tropical Medicine and Hygiene. 2010;83: 607–613. doi:10.4269/ajtmh.2010.10-0005

110. Brown JJ, Pascual M, Wimberly MC, Johnson LR, Murdock CC. Humidity – The overlooked variable in the thermal biology of MOSQUITO-BORNE disease. Ecology Letters. 2023; ele.14228. doi:10.1111/ele.14228

