## Supplementary material for "Disclosing temperature sensitivity of West Nile virus transmission: novel computational approaches to mosquito-pathogen trait responses": S1 Text

### SI1 Details on functions fitted to mosquito-pathogen traits

#### Juvenile mosquito development rate

We model the juvenile mosquito development rate $\delta_J(T)$ from egg hatch to adult emergence (i.e., encompassing the larva and pupa stage) by a modified Briére function [1] that depends on temperature $(T)$:

$$f^B(T; q, T_{\min}, T_{\max}) = \begin{cases} \frac{q}{c} T(T - T_{\min})\sqrt{T_{\max} - T}, & \max(0, T_{\min}) < T < T_{\max} \\ 0, & \text{else} \end{cases} \qquad (SI1.1)$$

The Briére function describes a left-skewed unimodal response to temperature and has proven to be a suitable model for mosquito development rates in several previous works [2–4]. We chose a constant scaling factor $c = 100000$ to bring the parameter $q$ on a similar scale as $T_{\min}$ and $T_{\max}$ which helps to improve speed and stability of the MCMC fitting procedure. Although the present data shows only little evidence of a decrease in development rates at high temperatures, we chose to describe this trait with a unimodal function instead of a monotonic function to follow the established theory for ectotherm metabolic rates [5]. To fit this trait, we utilized the reciprocal of development time data measured from egg hatch to adult emergence. We only made one exception to this rule where one of the studies included in our analysis measured development time to adult emergence starting from the second larva instar stage [6], a small inconsistency that we were willing to accept in favour of having a larger dataset. We want to highlight an important technicality that improves our approach of fitting development rates compared to some previous analyses [2–4,7,8]. Specifically, we did not infer development rate performance from data on juvenile survival. Specifically, we agree with the recommendation of Von Schmalensee et al. [9] that 100% end mortality in juvenile development stages should not be used as an indication of a development rate of zero. In fact, in these cases no observation of the development time was possible. These observations only indicate that survival is highly unlikely for the time it takes to complete development but do not indicate how fast juveniles would have developed, would they have survived, nor how fast development would progress given only brief, potentially non-lethal, exposures to the respective temperature. Adding such "artificial" data points when fitting parametric functions not only impacts model fits at the extreme temperatures but also biases results at moderate temperatures and should therefore be avoided.

#### Juvenile mosquito survival

As for juvenile development, we modelled juvenile survival $p_J(T)$ from egg hatch to adult emergence. To describe the typically symmetric temperature response of this trait [5], we used a modified quadratic function:

$$f^Q(T; q, T_{\min}, T_{\max}) = \begin{cases} \min\left(\frac{q}{c}(T - T_{\min})(T_{\max} - T), 1\right), & T_{\min} < T < T_{\max} \\ 0, & \text{else} \end{cases} \qquad (SI1.2)$$

As for the Briére function introduced earlier, we used constant scaling factor $c = 1000$ to bring the parameter $q$ on a similar scale as $T_{\min}$ and $T_{\max}$. To fit this trait, we used data reporting the percentage of juvenile mosquitoes surviving from egg hatch to adult emergence. As for juvenile development we included data from one study in which juvenile survival was measured from the second larva instar to adult emergence [6].

**Egg viability and egg hatching rate**

Our approaches to model egg viability $p_E(T)$ defined as the percentage of eggs hatching at a given temperature and egg hatching rate $\delta_E(T)$ measured as the inverse of the time from egg laying to larva emergence follow a similar reasoning as for juvenile traits. As a result, we model the temperature response of egg viability using the modified quadratic function $f^Q(T; q, T_{\min}, T_{\max})$ (Equation $(S1.2)$) and egg hatching rate by the modified Briére function $f^B(T; q, T_{\min}, T_{\max})$ (Equation $(S1.1)$). In case of the egg hatching rate, we use a scaling factor of $q = 16000$ to bring the parameter on a similar scale as for the juvenile development rate.

**Adult lifespan**

Like for the other mosquito life-history traits, the temperature dependence of mosquito adult lifespan $\mathrm{lf}(T)$ is often observed to follow a unimodal response [3,10]. However, as noted in previous work [7], the present data does not indicate a decrease in adult lifespan at low temperatures for any of the considered species. In fact, several of the species of interest in this analysis diapause in the adult stage during winter. For these species it is not obvious at which cold temperatures adult lifespan would cease and which shape the temperature response should have in this temperature range. Therefore, we follow the previous work [7] and model adult lifespan by a linearly decreasing function truncated at zero:

$$f^L(T; \alpha, \beta) = \begin{cases} -\beta T + \alpha, & T < \frac{\alpha}{\beta} \\ 0, & \text{else} \end{cases} \qquad (SI1.3)$$

As a conservative approach to trait performance at low temperatures, we plateau the linear function after model fitting at the lowest observed temperature point in the dataset across all species (14°C). During the MCMC simulations, we generated posterior samples for the parameters $\beta$ and $T_{\max} \coloneqq \alpha/\beta$ since we found it more intuitive to define hyperpriors for the population-level parameters of the transformed parameter $T_{\max}$ than for the intercept $\alpha$ directly. When data was available for adult mosquito males and females separately, only female data was used to fit this trait.

**Biting rate**

To derive the relationship between temperature and adult biting rate $a(T)$ we fitted the modified Briére function $f^B(T; q, T_{\min}, T_{\max})$ (Equation $(S1.1)$) with a scaling factor of $q = 43000$ to bring the parameter on a similar scale as for the juvenile development rate. The data utilized for model fitting represent the reciprocal of the gonotrophic cycle duration, measured from blood meal ingestion to egg laying. As for juvenile development rates we decided to fit a unimodal function despite limited evidence for a decrease of biting rates at high temperatures in the data. With this decision we again follow the established theory [5]. As for juvenile development rate we advise against the practice to infer a biting rate of zero from the observations that no adult mosquito survived the time from taking a blood meal over blood digestion to egg maturation and eventually laying eggs [3,4,7]. These observations do not convey clear information about the length of the gonotrophic cycle at a given temperature but rather offer rough information about the length of the gonotrophic cycle relative to adult lifespan.

**Fecundity**

Mosquitoes from the genus *Culex* typically bite once per gonotrophic cycle and lay their eggs in single egg rafts. Therefore, we model the female mosquito egg laying rate (eggs per female per day) $\beta(T)$, as the product of biting rate $a(T)$ and eggs per egg raft $\beta_{ER}$:

$$\beta(T) = a(T)\beta_{\mathrm{ER}} \qquad (SI1.4)$$

Data on the potential temperature dependence of eggs per egg raft $\beta_{\mathrm{ER}}$ of *Culex* mosquitoes is very limited. We could only identify three studies providing data on the temperature sensitivity of this trait for *Cx.* mosquitoes [11,12]. The data is confined to the temperature range 15-30°C and while the data on Cx. pipiens molestus shows the most notable reduction in $\beta_{\mathrm{ER}}$ at high temperature, the remaining data indicates that the trait is somewhat stable over the observed temperature range, although with tendencies for highest trait values at intermediate temperatures. Overall, we found the information on $\beta_{\mathrm{ER}}$ too scarce to fit a temperature-dependent function. Therefore, we decided to set this trait to a constant value determined by the mean across all observations in the dataset (given by 140).

In a previous analysis the expression for eggs per female per day was multiplied by another temperature dependent trait representing the proportion of female mosquitoes that lay an egg raft after taking a blood meal (therein called “proportion ovipositing”) [7]. Including this additional trait would be extend the model if it describes an additional, temperature-sensitive aspect of mosquito biology that is independent of the traits described so far. Indeed, in the study by Tekle on *Culex quinquefasciatus* and *Culex pipiens* [13] it was observed that mosquitoes that didn’t lay eggs at 10°C and 15°C showed completely developed eggs when dissected. On the other hand, it was also found that *Culex quinquefasciatus* held at 5°C did simply not live long enough to digest their blood meals. Similarly, the study by Mahmood and Crans on *Culiseta melanura* [14] found that none of the blood fed mosquitoes survived long enough to lay eggs at 34°C. This indicates that the temperature sensitivity of the proportion of ovipositing female mosquitoes arises at least in part from the relation between gonotrophic cycle duration and adult female lifespan. In fact, these two traits already indicate that the proportion of ovipositing females would decrease both at very low temperatures (because the gonotrophic cycle becomes extremely long) and high temperatures (where adult lifespan approaches zero). This effect can already be incorporated into models based on the derived temperature response of adult lifespan and biting rate. By multiplying the expression for eggs per female per day by the proportion of ovipositing females we would risk accounting for the described effect twice. Therefore, we decided not to include the proportion of ovipositing females as a separate trait into our models.

**Mosquito infection probability and extrinsic incubation period**

To successfully utilize mosquitoes as transmission vectors, viruses have to overcome several barriers after the mosquito ingested infected blood [15]. Here, we model mosquitoes’ ability to transmit WNV as a two-component process. The first component, hereafter referred to as mosquito infection probability $b_M(T)$ (which we conceptualize as classifying a mosquito to be competent), reflects the intrinsic susceptibility of mosquitoes to develop a WNV midgut infection when exposed to infected blood and lacks a temporal dimension. The second component describes the average time required for a competent mosquito to develop a salivary gland infection after exposure to infected blood (here equated with the ability of the mosquito to transmit the virus when biting a susceptible host), which is called the extrinsic incubation period $\mathrm{EIP}(\mathrm{T})$. To describe the temperature-dependence of these two traits we used data from vector competence experiments that tested mosquitoes for WNV infection in their bodies or their salivary glands for at least three different time points after exposure to infected blood.

Specifically, to describe the mosquito infection probability $b_M(T)$, we utilized data sampling the percentage of mosquitoes with an infected body upon all exposed mosquitoes (# infected mosquitoes/# exposed mosquitoes) which was available for *Culex pipiens* exposed to WNV genotypes NY99 and WN02 [16,17]. A visual inspection of this data showed no obvious temporal trends with regards to the number of days post infection (dpi). Moreover, increases in $b_M(T)$ with temperature seemed to be well described by a sigmoidal function:

$$b_M(T;\alpha,\beta) = \frac{1}{1+e^{-(\beta T+\alpha)}} \qquad (SI1.5)$$

To describe the temperature-dependence of the extrinsic incubation period $\mathrm{EIP}(T)$ we follow a similar approach as described by Brady et al. [18]. We fit this trait based on data that sampled the percentage of mosquitoes with detectable virus in the salivary glands (hereafter referred to as transmitting mosquitoes) upon the mosquitoes with an infected body (# transmitting mosquitoes/# infected mosquitoes). This data was available for WNV genotype NY99 in *Culex tarsalis* [19] and South-African genotype H442 in *Culex univittatus* [20]. The available data indicates that the percentage of transmitting mosquitoes upon infected mosquitoes increases monotonically with dpi and that the required dpi to reach 50% of transmitting mosquitoes (here considered as the expected extrinsic incubation period) decreases monotonically with temperature. Moreover, the study by Cornel on *Culex univittatus* indicates that, if observations are conducted for a sufficiently large number of dpi, this percentage eventually reaches 100% independent of the temperature setting (although at the lowest temperature of 14°C the percentage showed large fluctuations for high dpi ranging from 34% to 100%) [20]. Therefore, for a given temperature $T$ we modelled the relationship between dpi $t$ and the percentage of transmitting mosquitoes upon all infected mosquitoes with a cumulative Gaussian distribution:

$$\Phi(t;\mathrm{EIP}(T;\alpha,\beta),\sigma_{\mathrm{EIP}}^2) \qquad (SI1.6)$$

We modelled the mean $\mathrm{EIP}(T)$ of this Gaussian distribution (i.e., the point in time at which 50% of the infected mosquitoes are expected to be transmitting), which serves as our estimate for the extrinsic incubation period, with an exponential decay function:

$$\mathrm{EIP}(T;\alpha,\beta) = \exp(-\beta T+\alpha) \qquad (SI1.7)$$

Additional data to inform $\mathrm{EIP}(T)$ was available from the study by Kilpatrick et al. on WN02 and NY99 in *Culex pipiens* [16]. However, this study did not report the percentage of transmitting mosquitoes upon all infected mosquitoes but upon all exposed mosquitoes (# transmitting mosquitoes/# exposed mosquitoes). Therefore, before this data was used as part of the fitting of $\mathrm{EIP}(T)$, it was rescaled by the inverse of our predictions for the mosquito infection probability $b_M(T)$ (# exposed mosquitoes/# infected mosquitoes).

**Considerations of alternative models for the pathogen-related traits**

The extrinsic incubation period model introduced above assumes a monotonic relationship between dpi and the percentage of transmitting mosquitoes. For some mosquito-pathogen combinations it has been observed that for a given temperature the relationship between dpi and the percentage of transmitting mosquitoes is not necessarily monotonic. In particular experiments on *Plasmodium falciparum* in *Anopheles stephensi* [21] and Western Equine encephalitis virus in *Cx. tarsalis* [22] at moderate to high temperatures demonstrated an initial increase in the fraction of transmitting mosquitoes followed by a decrease along dpi. In these cases, the approach described above would seem insufficient. However, none of the available data on WNV infection in *Cx.* mosquitoes demonstrated compelling evidence for the presence of such an effect.

In contrast to our modelling approach, some previous studies have described the temperature response of vector competence, and in some cases of the extrinsic incubation rate (inverse of the extrinsic incubation period), by unimodal functions [3,7,8,23,24]. Three of these studies used overlapping data sources to our study [7,23,24]. We look here at the reasoning these studies applied to fit unimodal functions and outline why we decided against it. For the sake of simplicity, we explain this with reference to the study by Shocket et al. [7] and contrast their approach to ours. A similar logic applies to the other two studies.

We fit a monotonic temperature response function for the mosquito infection probability (the probability that an exposed mosquito develops a midgut infection, here conceptualized as classifying the mosquito to be competent) because the available data does not support a unimodal response (see Figure 11 in the main text). We also found that a monotonic model for the extrinsic incubation period (which implies a monotonic model for the extrinsic incubation rate), when it is defined as in our work (see above), is satisfactory to describe the data available for this trait. Only in the case of WNV H442 in *Cx. univittatus* the data suggested an increase of the extrinsic incubation period between the 26°C and 30°C temperature setting which our model fails to capture (see Figure 12C in the main text). We concluded that the data does not provide enough evidence to motivate a unimodal model for the extrinsic incubation rate.

In contrary, Shocket et al. [7] conceptualized vector competence as the product of two probabilities: infection efficiency (probability that an exposed mosquitoes develops a disseminated infection, i.e. virus positive legs indicating that the virus spread beyond the midgut) and transmission efficiency (probability that a mosquitoes with disseminated infection develops a salivary gland infection). To fit the temperature response function for infection efficiency and transmission efficiency, or directly for vector competence as the product of the two, Shocket et al. [7] extracted the corresponding data on mosquitoes with disseminated infection and on transmitting mosquitoes from the highest dpi settings observed within the experiment for each temperature. The data extracted like this indeed appear to indicate temperature-dependent traits that first increase with temperature and eventually decrease (i.e., potentially extending the mosquito infection probability as defined by us). For example, in the data on WN02 in *Cx. pipiens* in Kilpatrick et al. [16] the fraction of transmitting mosquitoes in the highest available dpi observations increases from 15°C to 22°C and is slightly lower for 32°C than 22°C (see Figure 1C in Kilpatrick et al. [16] and Figure 12A in the main text). We want to note, however, that the traits derived like this depend on the lifespan of mosquitoes at the given temperature as well as on the total time length of the experimental studies. This is because these two factors impact the maximum dpi at which observation are generated in each temperature setting. At low temperatures the incubation period of mosquitoes can be extremely long, and experiments might be discontinued before mosquitoes have the chance to develop disseminated or salivary gland infections. In this case the observations extracted do not really reflect biological traits. High temperatures, on the other side, imply a short mosquito lifespan impacting the maximum observable dpi, making it again unlikely to observe a high percentage of mosquitoes with disseminated or salivary gland infections.

Shocket et al. [7] do not clearly state a definition for the extrinsic incubation rate, how it relates to the transmission efficiency trait, and how they collected the respective data to fit the extrinsic incubation rate. But it seems to us that the data for the extrinsic incubation rate was collected in a way that it roughly represents the reciprocal of the average time it takes for a mosquito to develop a salivary gland infection conditioned on reaching that state within the experiment. This definition makes the extrinsic incubation rate/period depend on mosquito lifespan and the total time length of the experimental studies, too. In contrast, we define the extrinsic incubation period as the dpi at which 50% of infected mosquitoes are expected to have a salivary gland infection (without the need to ever observe such a high percentage within the experiment) under the assumption that every mosquito with an infected body has the intrinsic ability to reach this state would it live long enough. In this way, we avoid that our estimates depend on the time length of the experimental studies. This definition also makes it straightforward to describe the effect observed at high temperatures by comparing the estimates for adult mosquito lifespan with the estimates for the extrinsic incubation period. For example, this relationship is incorporated into the basic reproduction number $R_0(T)$ via the term $e^{-\mu_M(T)\mathrm{EIP}(T)}$ that describes the fraction of mosquitoes that survive the extrinsic incubation period.

Introducing an additional (temperature-dependent) trait to our work, like the infection or transmission efficiency, would only be required if a part of the mosquitoes that show infected bodies lack the intrinsic susceptibility to ever develop a disseminated or a salivary gland infection. This would, for example, be indicated by the data if the percentage of transmitting mosquitoes upon all mosquitoes with infected body would stabilize below 100% with respect to dpi. However, this is not evident from the available data (see Figure 12A-D in the main text). The data provided by the works of Kilpatrick et al. and Reisen et al. [16,19], show no clear indication that the percentage of transmitting mosquitoes would not further increase if observations would have been possible at higher dpi. In the study by Cornel [20] observations were possible at exceptionally high dpi and the percentage of transmitting mosquitoes eventually reached 100% in all temperature settings (although notably fluctuating at 14°C). According to these observations, a need for describing vector competence of *Cx.* species for WNV by an additional (temperature-dependent) trait currently lacks clear empirical evidence. However, new experimental studies could try to shed further light on this discussion.

### SI2 Summary of mosquito-pathogen temperature response fits

Tables SI2.1-8 below provide a summary of the posterior parameter estimates (mean and 95% credible intervals) of the expected temperature response for all species and traits considered, including estimates of the parameters of hierarchical priors (population-level means, between-species standard deviations, between-experiment standard deviations).

**Table SI2.1.** Posterior mean and 95% credible interval of parameter estimates of hierarchical priors and of the expected temperature response by species (including the optimal temperature) for the juvenile development rate $\delta_J(T)$ modelled by a modified Briére function. *hierachical prior on log-scale of parameter

**Juvenile development rate $\delta_J(T)$ – Briére function**

| | $q$ | $T_{\mathrm{max}}$ | $T_{\mathrm{min}}$ | $T_{\mathrm{opt}}$ |
|---|---|---|---|---|
| *Ae. solicitans* [25] (n=7) | 5.18 (2.95 – 8.72) | 43.49 (39.59 – 49.54) | 2.25 (-3.01 – 7.82) | 35.04 (31.98 – 39.77) |
| *Ae. triseriatus* [25] (n=6) | 3.45 (2.14 – 5.25) | 41.76 (36.39 – 46.66) | 0.37 (-7.90 – 5.31) | 33.46 (29.26 – 37.22) |
| *Cs. inornata* [25,26] (n=7) | 3.22 (2.09 – 4.62) | 42.28 (36.28 – 48.16) | 0.47 (-6.94 – 5.14) | 33.89 (29.09 – 38.53) |
| *Cs. melanura* [27] (n=6) | 2.54 (1.27 – 4.06) | 41.87 (35.96 – 47.16) | 2.70 (-2.69 – 9.54) | 33.79 (29.09 – 37.98) |
| *Cx. pipiens molestus* [28–30] (n=17) | 2.99 (2.00 – 4.13) | 43.82 (39.85 – 50.76) | 2.98 (-1.86 – 8.15) | 35.38 (32.30 – 40.79) |

| | | | | |
|---|---|---|---|---|
| *Cx. pipiens pallens* [29,31] (n=9) | 3.04 (1.98 – 4.30) | 43.59 (39.69 – 49.74) | 1.80 (-3.65 – 6.86) | 35.06 (32.03 – 39.92) |
| *Cx. pipiens* [13,32–39] (n=62) | 3.96 (2.95 – 5.17) | 41.90 (38.83 – 45.84) | 1.57 (-2.80 – 5.19) | 33.69 (31.43 – 36.61) |
| *Cx. quinquefasciatus* [6,13,25,34,40,41] (n=42) | 3.31 (2.46 – 4.27) | 43.18 (40.63 – 46.75) | 1.76 (-3.04 – 5.98) | 34.74 (32.81 – 37.38) |
| *Cx. restuans* [25,26,34,37,42] (n=23) | 4.25 (2.95 – 5.86) | 42.71 (38.67 – 47.86) | 0.59 (-5.03 – 4.64) | 34.24 (31.20 – 38.15) |
| *Cx. salinarius* [25] (n=6) | 3.47 (2.20 – 5.16) | 42.75 (37.68 – 48.65) | 1.06 (-5.54 – 5.86) | 34.32 (30.29 – 39.08) |
| *Cx. tarsalis* [26,43] (n=13) | 2.94 (1.94 – 4.13) | 43.28 (39.72 – 48.82) | 1.80 (-3.69 – 6.59) | 34.82 (32.06 – 39.12) |
| Population level mean $\mu$ | 1.21* (0.91 – 1.50) | 42.78 (39.88 – 46.64) | 1.60 (-2.42 – 4.94) | – |
| Between species variability $\sigma$ | 0.29* (0.05 – 0.61) | 1.92 (0.07 – 5.77) | 2.17 (0.09 – 6.31) | – |
| Between experiment variability $\sigma^{\exp}$ | 0.21* (0.07 – 0.33) | 2.56 (0.60 – 4.72) | 3.79 (1.61 – 5.94) | – |

**Table SI2.2.** Posterior mean and 95% credible interval of parameter estimates of hierarchical priors and of the expected temperature response by species (including the optimal temperature) for the juvenile survival $p_J(T)$ modelled by a modified quadratic function. *hierachical prior on log-scale of parameter

**Juvenile survival $p_J(T)$ – Quadratic function**

| | $q$ | $T_{\max}$ | $T_{\min}$ | $T_{\mathrm{opt}}$ |
|---|---|---|---|---|
| *Ae. nigromaculis* [44] (n=6) | 4.42 (3.52 – 5.53) | 38.02 (34.50 – 42.50) | 10.79 (7.46 – 14.04) | 24.40 (21.90 – 27.07) |
| *Ae. solicitans* [25] (n=8) | 4.34 (3.43 – 5.24) | 38.33 (34.84 – 42.42) | 10.71 (7.49 – 13.70) | 24.52 (22.14 – 26.94) |
| *Ae. triseriatus* [25,45] (n=10) | 4.49 (3.66 – 5.67) | 34.89 (32.17 – 37.56) | 6.17 (2.18 – 9.51) | 20.53 (18.17 – 22.67) |
| *Ae. vexans* [44] (n=6) | 4.41 (3.51 – 5.46) | 37.96 (34.48 – 42.40) | 10.44 (7.03 – 13.85) | 24.20 (21.72 – 26.98) |
| *Cs. inornata* [25,26,44] (n=17) | 4.47 (3.62 – 5.58) | 32.09 (29.11 – 35.76) | 7.22 (4.69 – 9.60) | 19.65 (17.67 – 21.83) |
| *Cs. melanura* [27] (n=7) | 4.55 (3.66 – 6.11) | 34.90 (31.43 – 38.18) | 12.27 (8.22 – 16.21) | 23.59 (21.07 – 26.19) |
| *Cx. pipiens molestus* [28,30,46] (n=17) | 4.40 (3.55 – 5.37) | 35.51 (33.27 – 37.84) | 10.55 (8.04 – 12.75) | 23.03 (21.40 – 24.58) |
| *Cx. pipiens pallens* [11,31] (n=10) | 4.29 (3.38 – 5.14) | 38.79 (35.56 – 42.34) | 10.05 (7.54 – 12.57) | 24.42 (22.41 – 26.53) |

| | | | | |
|---|---|---|---|---|
| *Cx. pipiens* [13,33–39] (n=65) | 4.41 (3.73 – 5.15) | 36.14 (34.86 – 37.52) | 7.38 (5.56 – 9.06) | 21.76 (20.69 – 22.80) |
| *Cx. quinquefasciatus* [6,11,13,25,34,40,41,46–48] (n=69) | 4.44 (3.72 – 5.25) | 36.89 (35.52 – 38.28) | 9.44 (7.94 – 10.89) | 23.17 (22.22 – 24.13) |
| *Cx. restuans* [25,26,34,37,42] (n=25) | 4.34 (3.49 – 5.22) | 34.94 (32.93 – 37.01) | 5.09 (1.64 – 8.16) | 20.01 (18.06 – 21.83) |
| *Cx. salinarius* [25] (n=8) | 4.43 (3.57 – 5.54) | 34.29 (30.91 – 37.51) | 8.04 (4.52 – 11.20) | 21.16 (18.86 – 23.43) |
| *Cx. tarsalis* [26,49] (n=9) | 4.42 (3.52 – 5.49) | 36.46 (33.81 – 39.41) | 10.33 (7.45 – 12.92) | 23.39 (21.40 – 25.37) |
| Population level mean $\mu$ | 1.48* (1.34 – 1.62) | 36.08 (34.28 – 37.99) | 9.13 (7.21 – 11.09) | – |
| Between species variability $\sigma$ | 0.07* (0.00 – 0.23) | 2.59 (0.68 – 4.92) | 2.79 (1.19 – 5.07) | – |
| Between experiment variability $\sigma^{\text{exp}}$ | 0.18* (0.02 – 0.33) | 1.88 (1.12 – 2.76) | 1.47 (0.19 – 2.79) | – |

**Table SI2.3.** Posterior mean and 95% credible interval of parameter estimates of hierarchical priors and of the expected temperature response by species (including the optimal temperature) for the egg development rate $\delta_E(T)$ modelled by a modified Briére function. *hierachical prior on log-scale of parameter

| Egg development rate $\boldsymbol{\delta_E(T)}$ – Briére function | | | | |
|---|---|---|---|---|
| | $\boldsymbol{q}$ | $\boldsymbol{T_{\max}}$ | $\boldsymbol{T_{\min}}$ | $\boldsymbol{T_{\text{opt}}}$ |
| *Cs. melanura* [27] (n=4) | 3.18 (1.87 – 4.79) | 51.06 (41.45 – 63.07) | 3.40 (-2.62 – 8.95) | 41.22 (33.71 – 50.58) |
| *Cx. pipiens molestus* [28,29] (n=11) | 3.66 (2.40 – 5.25) | 52.78 (44.31 – 65.32) | 2.99 (-2.74 – 7.93) | 42.55 (35.98 – 52.28) |
| *Cx. pipiens pallens* [29,31] (n=9) | 3.65 (2.37 – 5.30) | 52.07 (44.01 – 64.01) | 2.64 (-3.34 – 7.54) | 41.94 (35.69 – 51.18) |
| *Cx. pipiens* [32,37] (n=9) | 3.43 (2.20 – 4.95) | 51.69 (43.24 – 63.86) | 2.53 (-3.59 – 7.43) | 41.62 (35.11 – 51.12) |
| *Cx. quinquefasciatus* [50] (n=4) | 3.52 (2.15 – 5.37) | 50.75 (42.29 – 62.58) | 2.20 (-4.45 – 7.31) | 40.84 (34.24 – 50.03) |
| *Cx. restuans* [37] (n=6) | 3.37 (2.08 – 5.01) | 50.90 (41.46 – 63.06) | 2.15 (-4.49 – 7.23) | 40.95 (33.59 – 50.38) |
| *Cx. theileri* [51] (n=9) | 3.47 (2.16 – 5.18) | 51.70 (43.44 – 63.83) | 3.07 (-2.92 – 8.10) | 41.69 (35.30 – 51.02) |
| Population level mean $\mu$ | 1.22* (0.82 – 1.59) | 51.45 (43.64 – 62.72) | 2.72 (-2.63 – 7.10) | – |
| Between species standard deviation $\sigma$ | 0.16* (0.04 – 0.35) | 1.94 (0.12 – 5.84) | 1.51 (0.10 – 4.54) | – |
| Between experiment variability $\sigma^{\text{exp}}$ | 0.15* (0.07 – 0.27) | 2.43 (0.86 – 4.85) | 3.11 (1.52 – 5.20) | – |

**Table SI2.4.** Posterior mean and 95% credible interval of parameter estimates of hierarchical priors and of the expected temperature response by species (including the optimal temperature) for adult mosquito lifespan $lf(T)$ modelled by a modified linear function. *hierachical prior on log-scale of parameter

**Adult lifespan $\mathbf{lf}(T)$ – Linear function**

| | $\boldsymbol{\beta}$ | $\boldsymbol{T_{\max}}$ |
|---|---|---|
| *Ae. taeniorhynchus* [52] (n=3) | 3.32 (1.67 – 5.73) | 35.97 (31.65 – 41.43) |
| *Cx. pipiens molestus* [28,29,46] (n=14) | 3.42 (2.13 – 5.12) | 33.34 (30.05 – 36.63) |
| *Cx. pipiens pallens* [29,53] (n=9) | 3.33 (1.96 – 5.11) | 34.59 (31.26 – 38.58) |
| *Cx. pipiens* [34,36,39,54] (n=42) | 4.42 (3.36 – 5.58) | 34.05 (32.24 – 35.89) |
| *Cx. quinquefasciatus* [34,46,47,53] (n=25) | 5.18 (3.73 – 6.84) | 33.87 (31.88 – 35.93) |
| *Cx. restuans* [34] (n=5) | 4.16 (2.13 – 7.58) | 32.48 (27.34 – 36.98) |
| *Cx. tarsalis* [43] (n=33) | 2.08 (1.50 – 2.84) | 37.66 (34.42 – 41.51) |
| Population level mean $\mu$ | 1.26* (0.85 – 1.66) | 34.57 (32.06 – 37.27) |
| Between species standard deviation $\sigma$ | 0.44* (0.19 – 0.90) | 2.48 (0.96 – 4.65) |
| Between experiment standard deviation $\sigma^{\exp}$ | 0.32* (0.19 – 0.48) | 1.86 (1.19 – 2.62) |

**Table SI2.5.** Posterior mean and 95% credible interval of parameter estimates of hierarchical priors and of the expected temperature response by species (including the optimal temperature) for adult mosquito biting rate $a(T)$ modelled by a modified Briére function. *hierachical prior on log-scale of parameter

**Biting rate $\boldsymbol{a(T)}$ – Briére function**

| | $\boldsymbol{q}$ | $\boldsymbol{T_{\max}}$ | $\boldsymbol{T_{\min}}$ | $\boldsymbol{T_{\mathrm{opt}}}$ |
|---|---|---|---|---|
| *Cs. melanura* [14] (n=4) | 3.54 (1.71 – 6.04) | 43.62 (35.01 – 57.38) | 1.15 (-8.82 – 7.89) | 35.04 (28.29 – 45.67) |
| *Cx. pipiens pallens* [31] (n=6) | 3.32 (1.71 – 5.36) | 45.66 (37.31 – 59.21) | 1.41 (-7.93 – 7.93) | 36.69 (30.19 – 47.22) |
| *Cx. pipiens* [13,37] (n=9) | 3.71 (1.86 – 6.17) | 45.10 (36.67 – 59.09) | 1.93 (-7.93 – 8.74) | 36.31 (29.79 – 47.06) |
| *Cx. quinquefasciatus* [13,55] (n=9) | 3.24 (1.56 – 5.43) | 42.72 (34.25 – 56.48) | 1.59 (-7.84 – 8.08) | 34.37 (27.77 – 44.91) |
| *Cx. tarsalis* [55] (n=5) | 4.22 (1.80 – 7.77) | 40.75 (31.37 – 56.10) | 0.16 (-10.36 – 7.00) | 32.65 (25.24 – 44.59) |
| Population level mean $\mu$ | 1.23* (0.60 – 1.75) | 43.50 (36.22 – 56.46) | 1.29 (-7.96 – 7.30) | – |

| | | | | |
|---|---|---|---|---|
| Between species variability $\sigma$ | 0.23* (0.05 – 0.51) | 2.86 (0.15 – 8.24) | 1.86 (0.12 – 5.57) | – |
| Between experiment variability $\sigma^{\text{exp}}$ | 0.20* (0.09 – 0.35) | 3.29 (0.99 – 6.37) | 3.40 (1.64 – 5.74) | – |

**Table SI2.6.** Posterior mean and 95% credible interval of parameter estimates of hierarchical priors and of the expected temperature response by species (including the optimal temperature) for egg viability $p_E(T)$ modelled by a modified quadratic function. *hierachical prior on log-scale of parameter

**Egg viability $p_E(T)$ – Quadratic function**

| | $q$ | $T_{\text{max}}$ | $T_{\text{min}}$ | $T_{\text{opt}}$ |
|---|---|---|---|---|
| *Cx. pipiens molestus* [12,28] (n=19) | 5.65 (3.67 – 8.57) | 33.27 (30.84 – 35.95) | 6.61 (2.25 – 10.14) | 19.94 (17.58 – 22.11) |
| *Cx. pipiens pallens* [31] (n=7) | 5.44 (3.13 – 8.60) | 39.44 (35.61 – 43.31) | 6.66 (2.67 – 10.70) | 23.05 (20.42 – 25.72) |
| *Cx. quinquefasciatus* [12,50] (n=11) | 5.86 (3.78 – 9.00) | 38.46 (35.14 – 41.93) | 12.38 (7.86 – 15.61) | 25.42 (22.63 – 27.86) |
| *Cx. theileri* [51] (n=13) | 6.29 (3.89 – 11.08) | 38.83 (35.34 – 42.33) | 9.78 (6.14 – 12.91) | 24.31 (21.97 – 26.58) |
| Pop. level mean $\mu$ | 1.73* (1.33 – 2.17) | 37.48 (33.87 – 41.11) | 8.88 (5.01 – 12.51) | – |
| Between species variability $\sigma$ | 0.10* (0.00 – 0.34) | 3.06 (1.49 – 5.24) | 3.05 (1.49 – 5.12) | – |
| Between experiment variability $\sigma^{\text{exp}}$ | 0.35* (0.09 – 0.64) | 1.87 (1.14 – 2.76) | 1.62 (0.41 – 3.53) | – |

**Table SI2.7.** Posterior mean and 95% credible interval of parameter estimates of hierarchical priors and of the expected temperature response by experiment for the mosquito infection probability $b_M(T)$ modelled by a sigmoidal function.

**Mosquito infection probability $b_M(T)$ – sigmoidal function**

| | $\alpha$ | $\beta$ |
|---|---|---|
| WNV NY99 in *Cx pipiens* Dohm et al. [17] (n=33) | -4.47 (-5.23 – (-3.72)) | 0.29 (0.25 – 0.33) |
| WNV WN02 in *Cx. pipiens* Kilpatrick et al. [16] (n=44) | -4.64 (-5.22 – (-4.08)) | 0.25 (0.22 – 0.28) |
| WNV NY99 in Cx. pipiens Kilpatrick et al. [16] (n=44) | -5.19 (-5.79 – (-4.61)) | 0.28 (0.25 – 0.31) |
| Population level mean $\mu$ | -4.76 (-5.40 – (-4.13)) | 0.27 (0.24 – 0.31) |

**Table SI2.8.** Posterior mean and 95% credible interval of parameter estimates of hierarchical priors and of the expected temperature response by experiment for the extrinsic incubation period $EIP(T)$ modelled by an exponential decay function.

| **Extrinsic incubation period EIP($T$) – Exponential decay function** | | |
|---|---|---|
| | $\boldsymbol{\alpha}$ | $\boldsymbol{\beta}$ |
| WNV WN02 in *Cx. pipiens* Kilpatrick et al. [16] (n=42) | 5.35 (5.10 – 5.61) | 0.09 (0.08 – 0.10) |
| WNV NY99 in Cx. pipiens Kilpatrick et al. [16] (n=42) | 5.35 (5.09 – 5.62) | 0.08 (0.07 – 0.10) |
| WNV H442 in *Cx. univittatus* Cornel et al. [20] (n=67) | 4.98 (4.74 – 5.22) | 0.11 (0.09 – 0.12) |
| WNV NY99 in *Cx. tarsalis* Reisen et al. [19] (n=39) | 5.28 (5.05 – 5.53) | 0.11 (0.10 – 0.12) |
| Population level mean $\mu$ | 5.24 (5.02 – 5.47) | 0.10 (0.08 – 0.11) |

### SI3 Formal mathematical description of Bayesian hierarchical models and (hyper)prior specifications

Below we list the formal mathematical descriptions of the Bayesian hierarchical models that we used to fit the temperature response of each mosquito-pathogen trait. In addition, Table SI3.1 provides an overview of all the (hyper)priors that we specified before model fitting. In the descriptions below $y_{ij,T}$ (and similarly $n_{j,T}$ and $p_{j,t,T}$ for the pathogen-related traits) refers to life-history trait performance data measured for species $i$ in experiment $j$ at temperature $T$ ($t$ denotes days post infection). While the life-history trait data is grouped by species and by the different experiments on each species, the pathogen-related traits are only grouped by experiment identity since here we do not have sufficient data available to estimate parameters for groups of experiments on the same mosquito species or on the same mosquito species and virus strain combination. Our reasoning behind hyperprior choices is outlined in the main text. Additionally, we assigned a vague prior to the standard deviations $s$ of the normal distribution likelihoods. The model for the extrinsic incubation period also requires a prior for the standard deviation in extrinsic incubation period $s_{\mathrm{EIP}}$ in each experiment and each temperature setting. We found that model fits were satisfactory by using a single parameter for $s_{\mathrm{EIP}}$ that is shared across experiment-temperature combinations and accordingly defined a single prior distribution for this parameter. Below, we denote the cumulative gaussian distribution by $\Phi$, the normal distribution by $N$, the half-normal distribution by $HN$, the binomial distribution by $\mathrm{Bin}$, and the gamma distribution by $\Gamma$.

**Juvenile mosquito development rate**
Likelihood:

$$y_{ij,T}|q_{ij}, T_{\mathrm{min}_{ij}}, T_{\mathrm{max}_{ij}}, s \sim N\left(f^B\left(T; q_{ij}, T_{\mathrm{min}_{ij}}, T_{\mathrm{max}_{ij}}\right), s^2\right)$$

Priors:

$$\ln(q_{ij})|q_i, \sigma_q^{\exp} \sim N\left(\ln(q_i), \sigma_q^{\exp 2}\right)$$
$$\ln(q_i)|\mu_q, \sigma_q \sim N(\mu_q, \sigma_q^2)$$
$$T_{\mathrm{min}_{ij}}|T_{\mathrm{min}_i}, \sigma_{T_{\mathrm{min}}}^{\exp} \sim N\left(T_{\mathrm{min}_i}, \sigma_{T_{\mathrm{min}}}^{\exp\ 2}\right)$$

$$T_{\min_i}|\mu_{T_{\min}}, \sigma_{T_{\min}} \sim N(\mu_{T_{\min}}, \sigma_{T_{\min}}^2)$$
$$T_{\max_{ij}}|T_{\max_i}, \sigma_{T_{\max}}^{\exp} \sim N\left(T_{\max_i}, \sigma_{T_{\max}}^{\exp\ 2}\right)$$
$$T_{\max_i}|\mu_{T_{\max}}, \sigma_{T_{\max}} \sim N(\mu_{T_{\max}}, \sigma_{T_{\max}}^2)$$
$$s \sim HN(0,1)$$

Hyperpriors:

$$\mu_q \sim N(1.5,1)$$
$$\mu_{T_{\min}} \sim N(5,10^2)$$
$$\mu_{T_{\max}} \sim N(40,10^2)$$
$$\sigma_q \sim HN(0,1)$$
$$\sigma_{T_{\min}} \sim HN(0,10^2)$$
$$\sigma_{T_{\max}} \sim HN(0,10^2)$$
$$\sigma_q^{\exp} \sim HN(0,1)$$
$$\sigma_{T_{\min}}^{\exp} \sim HN(0,10^2)$$
$$\sigma_{T_{\max}}^{\exp} \sim HN(0,10^2)$$

**Juvenile mosquito survival**

Likelihood:

$$y_{ij,T}|q_{ij}, T_{\min_{ij}}, T_{\max_{ij}}, s \sim N\left(f^Q\left(T; q_{ij}, T_{\min_{ij}}, T_{\max_{ij}}\right), s^2\right)$$

Priors:

$$\ln(q_{ij})|q_i, \sigma_q^{\exp} \sim N\left(\ln(q_i), \sigma_q^{\exp 2}\right)$$
$$\ln(q_i)|\mu_q, \sigma_q \sim N(\mu_q, \sigma_q^2)$$
$$T_{\min_{ij}}|T_{\min_i}, \sigma_{T_{\min}}^{\exp} \sim N\left(T_{\min_i}, \sigma_{T_{\min}}^{\exp\ 2}\right)$$
$$T_{\min_i}|\mu_{T_{\min}}, \sigma_{T_{\min}} \sim N(\mu_{T_{\min}}, \sigma_{T_{\min}}^2)$$
$$T_{\max_{ij}}|T_{\max_i}, \sigma_{T_{\max}}^{\exp} \sim N\left(T_{\max_i}, \sigma_{T_{\max}}^{\exp\ 2}\right)$$
$$T_{\max_i}|\mu_{T_{\max}}, \sigma_{T_{\max}} \sim N(\mu_{T_{\max}}, \sigma_{T_{\max}}^2)$$
$$s \sim HN(0,1)$$

Hyperpriors:

$$\mu_q \sim N(1.5,1)$$
$$\mu_{T_{\min}} \sim N(5,10^2)$$
$$\mu_{T_{\max}} \sim N(40,10^2)$$
$$\sigma_q \sim HN(0,1)$$
$$\sigma_{T_{\min}} \sim HN(0,10^2)$$
$$\sigma_{T_{\max}} \sim HN(0,10^2)$$
$$\sigma_q^{\exp} \sim HN(0,1)$$
$$\sigma_{T_{\min}}^{\exp} \sim HN(0,10^2)$$
$$\sigma_{T_{\max}}^{\exp} \sim HN(0,10^2)$$

**Adult mosquito lifespan**

Likelihood:

$$y_{ij,T}|\alpha_{ij}, \beta_{ij}, s \sim N\left(f^L(T; \alpha_{ij}, \beta_{ij}), s^2\right)$$
$$T_{\max_{ij}} = \frac{\alpha_{ij}}{\beta_{ij}}$$

Priors:

$$\ln(\beta_{ij})|\beta_i, \sigma_\beta^{\exp} \sim N\left(\ln(\beta_i), \sigma_\beta^{\exp 2}\right)$$
$$\ln(\beta_i)|\mu_\beta, \sigma_\beta \sim N(\mu_\beta, \sigma_\beta^2)$$
$$T_{\max_{ij}}|T_{\max_i}, \sigma_{T_{\max}}^{\exp} \sim N\left(T_{\max_i}, \sigma_{T_{\max}}^{\exp\ 2}\right)$$
$$T_{\max_i}|\mu_{T_{\max}}, \sigma_{T_{\max}} \sim N\left(\mu_{T_{\max}}, \sigma_{T_{\max}}^2\right)$$
$$s \sim HN(0,10^2)$$

Hyperpriors:

$$\mu_\beta \sim N(1.5,1)$$
$$\mu_{T_{\max}} \sim N(35,10^2)$$
$$\sigma_\beta \sim HN(0,1)$$
$$\sigma_{T_{\max}} \sim \Gamma(4.86,1.88)$$
$$\sigma_\beta^{\exp} \sim HN(0,1)$$
$$\sigma_{T_{\max}}^{\exp} \sim \Gamma(18.93,10.07)$$

**Egg hatching rate and adult mosquito biting rate**
Likelihood:

$$y_{ij,T}|q_{ij}, T_{\min_{ij}}, T_{\max_{ij}}, s \sim N\left(f^B\left(T; q_{ij}, T_{\min_{ij}}, T_{\max_{ij}}\right), s^2\right)$$

Priors:

$$\ln\left(q_{ij}\right)|q_i, \sigma_q^{\exp} \sim N\left(\ln(q_i), \sigma_q^{\exp 2}\right)$$
$$\ln(q_i)|\mu_q, \sigma_q \sim N\left(\mu_q, \sigma_q^2\right)$$
$$T_{\min_{ij}}|T_{\min_i}, \sigma_{T_{\min}}^{\exp} \sim N\left(T_{\min_i}, \sigma_{T_{\min}}^{\exp\ 2}\right)$$
$$T_{\min_i}|\mu_{T_{\min}}, \sigma_{T_{\min}} \sim N\left(\mu_{T_{\min}}, \sigma_{T_{\min}}^2\right)$$
$$T_{\max_{ij}}|T_{\max_i}, \sigma_{T_{\max}}^{\exp} \sim N\left(T_{\max_i}, \sigma_{T_{\max}}^{\exp\ 2}\right)$$
$$T_{\max_i}|\mu_{T_{\max}}, \sigma_{T_{\max}} \sim N\left(\mu_{T_{\max}}, \sigma_{T_{\max}}^2\right)$$
$$s \sim HN(0,1)$$

Hyperpriors:

$$\mu_q \sim N(1.5,1)$$
$$\mu_{T_{\min}} \sim N(10,10^2)$$
$$\mu_{T_{\max}} \sim N(38,10^2)$$
$$\sigma_q \sim \Gamma(3.21,11.08)$$
$$\sigma_{T_{\min}} \sim \Gamma(1.36,0.63)$$
$$\sigma_{T_{\max}} \sim \Gamma(1.34,0.7)$$
$$\sigma_q^{\exp} \sim \Gamma(7.92,37.61)$$
$$\sigma_{T_{\min}}^{\exp} \sim \Gamma(9.73,2.57)$$
$$\sigma_{T_{\max}}^{\exp} \sim \Gamma(4.98,1.95)$$

**Egg viability**
Likelihood:

$$y_{ij,T}|q_{ij}, T_{\min_{ij}}, T_{\max_{ij}}, s \sim N\left(f^Q\left(T; q_{ij}, T_{\min_{ij}}, T_{\max_{ij}}\right), s^2\right)$$

Priors:

$$\ln(q_{ij})|q_i, \sigma_q^{\exp} \sim N\left(\ln(q_i), \sigma_q^{\exp 2}\right)$$
$$\ln(q_i)|\mu_q, \sigma_q \sim N(\mu_q, \sigma_q^2)$$
$$T_{\min_{ij}}|T_{\min_i}, \sigma_{T_{\min}}^{\exp} \sim N\left(T_{\min_i}, \sigma_{T_{\min}}^{\exp\ 2}\right)$$
$$T_{\min_i}|\mu_{T_{\min}}, \sigma_{T_{\min}} \sim N(\mu_{T_{\min}}, \sigma_{T_{\min}}^2)$$
$$T_{\max_{ij}}|T_{\max_i}, \sigma_{T_{\max}}^{\exp} \sim N\left(T_{\max_i}, \sigma_{T_{\max}}^{\exp\ 2}\right)$$
$$T_{\max_i}|\mu_{T_{\max}}, \sigma_{T_{\max}} \sim N(\mu_{T_{\max}}, \sigma_{T_{\max}}^2)$$
$$s \sim HN(0,1)$$

Hyperpriors:

$$\mu_q \sim N(1.5,1)$$
$$\mu_{T_{\min}} \sim N(10,10^2)$$
$$\mu_{T_{\max}} \sim N(38,10^2)$$
$$\sigma_q \sim \Gamma(1.2,16.37)$$
$$\sigma_{T_{\min}} \sim \Gamma(7.96,2.85)$$
$$\sigma_{T_{\max}} \sim \Gamma(4.86,1.88)$$
$$\sigma_q^{\exp} \sim \Gamma(2.79,15.63)$$
$$\sigma_{T_{\max}}^{\exp} \sim \Gamma(18.93,10.07)$$
$$\sigma_{T_{\min}}^{\exp} \sim \Gamma(3.45,2.35)$$

**Mosquito infection probability**
Likelihood:

$$n_{j,T}|\alpha_j, \beta_j \sim \mathrm{Bin}\left(N_{j,T}, b_M(T; \alpha_j, \beta_j)\right)$$

Priors:

$$\alpha_j|\mu_\alpha \sim N(\mu_\alpha, 0.3^2)$$
$$\beta_j|\mu_\beta \sim N(\mu_\beta, 0.02^2)$$

Hyperpriors:

$$\mu_\alpha \sim N(0,10^2)$$
$$\mu_\beta \sim N(0,1)$$

**Extrinsic incubation period**
Likelihood:

$$y_{j,T}|\alpha_j, \beta_j, s_{\mathrm{EIP}}, s \sim N\left(\Phi\left(\mathrm{t}; \mathrm{EIP}(T; \alpha_j, \beta_j), s_{\mathrm{EIP}}^2\right), s^2\right)$$

Priors:

$$\alpha_j|\mu_\alpha \sim N(\mu_\alpha, 0.1)$$
$$\beta_j|\mu_\beta \sim N(\mu_\beta, 0.01^2)$$
$$s \sim HN(0,1)$$
$$s_{\mathrm{EIP}} \sim HN(0,50^2)$$

Hyperpriors:

$$\mu_\alpha \sim N(0,10^2)$$
$$\mu_\beta \sim N(0,1)$$

**Table SI3.1.** (Hyper)priors used in the Bayesian hierachical models to fit trait temperature response functions. *hierachical prior on log-scale of parameter

| Trait | Population level mean | Between-species variability | Between-experiment variability | Standard deviation |
|---|---|---|---|---|
| $\delta_J$ | $\mu_q \sim N(1.5,1)$* | $\sigma_q \sim HN(0,1)$* | $\sigma_q^{\mathrm{exp}} \sim HN(0,1)$* | $s \sim HN(0,1)$ |
| | $\mu_{T_{\max}} \sim N(40,10^2)$ | $\sigma_{T_{\max}} \sim HN(0,10^2)$ | $\sigma_{T_{\max}}^{\mathrm{exp}} \sim HN(0,10^2)$ | |
| | $\mu_{T_{\min}} \sim N(5,10^2)$ | $\sigma_{T_{\min}} \sim HN(0,10^2)$ | $\sigma_{T_{\min}}^{\mathrm{exp}} \sim HN(0,10^2)$ | |
| $p_J$ | $\mu_q \sim N(1.5,1^2)$* | $\sigma_q \sim HN(0,1)$* | $\sigma_q^{\mathrm{exp}} \sim HN(0,1)$* | $s \sim HN(0,1)$ |
| | $\mu_{T_{\max}} \sim N(38,10^2)$ | $\sigma_{T_{\max}} \sim HN(0,10^2)$ | $\sigma_{T_{\max}}^{\mathrm{exp}} \sim HN(0,10^2)$ | |
| | $\mu_{T_{\min}} \sim N(10,10^2)$ | $\sigma_{T_{\min}} \sim HalfN(0,10^2)$ | $\sigma_{T_{\min}}^{\mathrm{exp}} \sim HN(0,10^2)$ | |
| $p_E$ | $\mu_q \sim N(1.5,1)$* | $\sigma_q \sim \Gamma(1.2,16.37)$* | $\sigma_q^{\mathrm{exp}} \sim \Gamma(2.79,15.63)$* | $s \sim HN(0,1)$ |
| | $\mu_{T_{\max}} \sim N(38,10^2)$ | $\sigma_{T_{\max}} \sim \Gamma(4.86,1.88)$ | $\sigma_{T_{\max}}^{\mathrm{exp}} \sim \Gamma(18.93,10.07)$ | |
| | $\mu_{T_{\min}} \sim N(10,10^2)$ | $\sigma_{T_{\min}} \sim \Gamma(7.96,2.85)$ | $\sigma_{T_{\min}}^{\mathrm{exp}} \sim \Gamma(3.45,2.35)$ | |
| lf | $\mu_\beta \sim N(1.5,1)$* | $\sigma_\beta \sim HN(0,1)$* | $\sigma_\beta^{\mathrm{exp}} \sim HN(0,1)$* | $s \sim HN(0,10^2)$ |
| | $\mu_{T_{\max}} \sim N(35,10^2)$ | $\sigma_{T_{\max}} \sim \Gamma(4.86,1.88)$ | $\sigma_{T_{\max}}^{\mathrm{exp}} \sim \Gamma(18.93,10.07)$ | |
| $\delta_E$ | $\mu_q \sim N(1.5,1)$* | $\sigma_q \sim \Gamma(3.21,11.08)$* | $\sigma_q^{\mathrm{exp}} \sim \Gamma(7.92,37.61)$* | $s \sim HN(0,1)$ |
| | $\mu_{T_{\max}} \sim N(40,10^2)$ | $\sigma_{T_{\max}} \sim \Gamma(1.34,0.7)$ | $\sigma_{T_{\max}}^{\mathrm{exp}} \sim \Gamma(4.98,1.95)$ | |
| | $\mu_{T_{\min}} \sim N(5,10^2)$ | $\sigma_{T_{\min}} \sim \Gamma(1.36,0.63)$ | $\sigma_{T_{\min}}^{\mathrm{exp}} \sim \Gamma(9.73,2.57)$ | |
| $a$ | $\mu_q \sim N(1.5,1)$* | $\sigma_q \sim \Gamma(3.21,11.08)$* | $\sigma_q^{\mathrm{exp}} \sim \Gamma(7.92,37.61)$* | $s \sim HN(0,1)$ |
| | $\mu_{T_{\max}} \sim N(40,10^2)$ | $\sigma_{T_{\max}} \sim \Gamma(1.34,0.7)$ | $\sigma_{T_{\max}}^{\mathrm{exp}} \sim \Gamma(4.98,1.95)$ | |
| | $\mu_{T_{\min}} \sim N(5,10^2)$ | $\sigma_{T_{\min}} \sim \Gamma(1.36,0.63)$ | $\sigma_{T_{\min}}^{\mathrm{exp}} \sim \Gamma(9.73,2.57)$ | |
| $b_M$ | $\mu_\beta \sim N(0,1)$ | Not applicable | $\sigma_\beta^{\mathrm{exp}} = 0.02$ | Not applicable |
| | $\mu_\alpha \sim N(0,10^2)$ | | $\sigma_\alpha^{\mathrm{exp}} = 0.3$ | |
| EIP | $\mu_\beta \sim N(0,1)$ | Not applicable | $\sigma_\beta^{\mathrm{exp}} = 0.01$ | $s \sim HN(0,1)$ |
| | $\mu_\alpha \sim N(0,10^2)$ | | $\sigma_\alpha^{\mathrm{exp}} = 0.1$ | $s_{\mathrm{EIP}} \sim HN(0,50^2)$ |

### SI4 Population-level trait estimates and sensitivity of pathogen-related traits to between-experiment variability

Figures SI4.1 and SI4.2 show population-level temperature responses estimated for each trait by sampling from the hierarchical priors. For mosquito life-history traits we sampled temperature response function parameters from the normal distribution hierarchical priors defined by the population-level mean and between-species standard deviations. Therefore, the resulting estimates reflect our uncertainty about a species' trait temperature response in the absence of data. For the pathogen-related traits we sampled temperature response function parameters from the normal distribution hierarchical priors defined by the population-level mean and between-experiment standard deviations. Here, the resulting estimates reflect our uncertainty about the response in a new experiment on an arbitrary mosquito species and WNV strain combination.

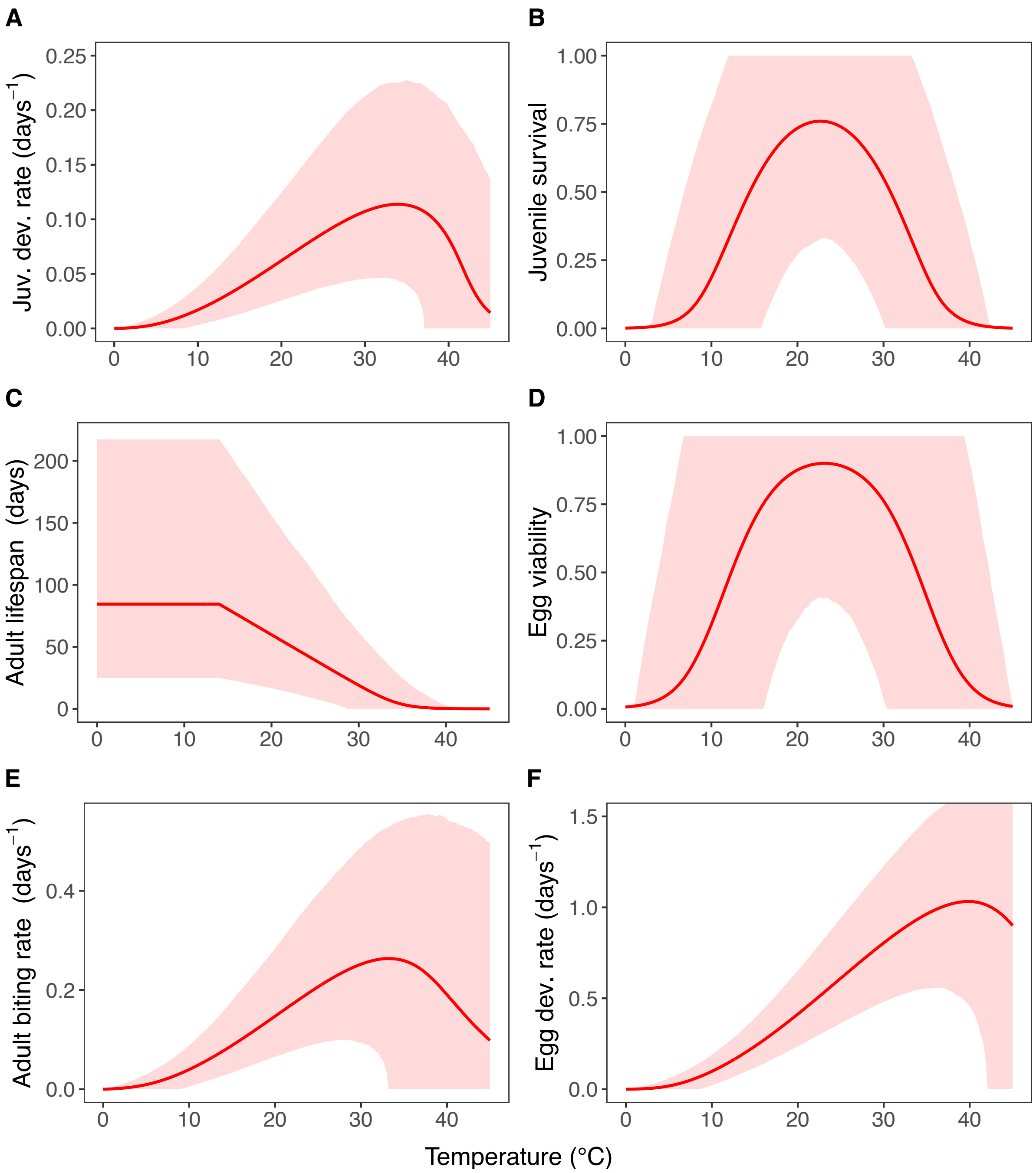


**Figure SI4.1.** Temperature responses of each life-history trait generated by sampling response function parameters from the normal distribution hierachical priors determined by the population-level means and between-species standard deviations. These responses reflect our uncertainty about a species' trait temperature response in the absence of data. Red solid lines represent posterior distribution means. Red shaded areas represent the corresponding central/equal-tailed 95% credible interval.

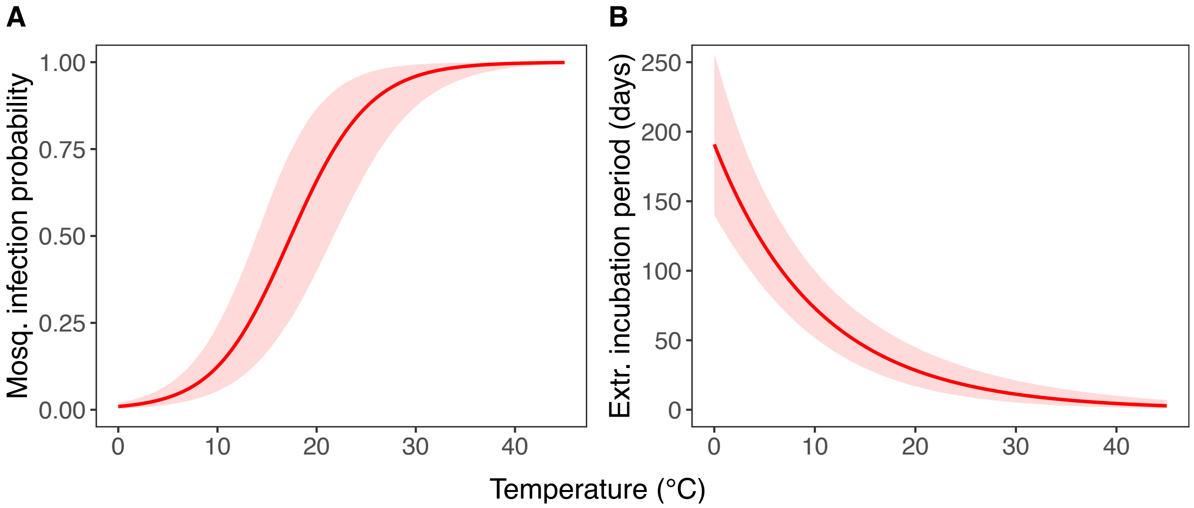


**Figure SI4.2.** Temperature responses of the pathogen-related traits generated by sampling response function parameters from the normal distribution hierachical priors determined by the population-level means and between-experiment standard deviations. These responses reflect our uncertainty about a trait temperature response in a new experiment with an arbitrary mosquito species and WNV strain combination. Red solid lines represent posterior distribution means. Red shaded areas represent the corresponding central/equal-tailed 95% credible interval.

Since we fixed the between-experiment standard deviations of the mosquito infection probability and extrinsic incubation period to subjectively chosen values, we investigated the sensitivity of our results to the choice of these values. In Figure SI4.3, we show our estimates of the expected temperature response of the mosquito infection probability in the experiment by Kilpatrick et al. [16] on WNV WN02 in Cx. pipiens and at the population-level for three different settings of the between-experiment standard deviations: a small variability setting with $\sigma_\beta^{\mathrm{exp}} = 0.01$ and $\sigma_\alpha^{\mathrm{exp}} = 0.15$, the moderate variability setting used in the main text with $\sigma_\beta^{\mathrm{exp}} = 0.02$ and $\sigma_\alpha^{\mathrm{exp}} = 0.3$, and a large variability setting with $\sigma_\beta^{\mathrm{exp}} = 0.02$ and $\sigma_\alpha^{\mathrm{exp}} = 0.3$. As can be seen from Figure SI4.3 there is no notable difference in the experiment-level temperature response estimates across the three different between-experiment variability settings. The mean of the population-level estimate is also relatively stable while the uncertainty of this estimate unsurprisingly changes between the three settings. We provide similar plots for the extrinsic incubation period in Figure SI4.4. The between-experiment variability settings used here are: a small variability setting with $\sigma_\beta^{\mathrm{exp}} = 0.005$ and $\sigma_\alpha^{\mathrm{exp}} = 0.05$, the moderate variability setting used in the main text with $\sigma_\beta^{\mathrm{exp}} = 0.01$ and $\sigma_\alpha^{\mathrm{exp}} = 0.1$, and a large variability setting with $\sigma_\beta^{\mathrm{exp}} = 0.02$ and $\sigma_\alpha^{\mathrm{exp}} = 0.2$.

We also investigated how the choice of between-experiment standard deviations for the pathogen-related traits might impact our $R_0^{\mathrm{rel}}(T)$ temperature optimum estimates. Figure SI4.5 shows the mean estimates of the optimal temperature of $R_0^{\mathrm{rel}}(T)$ for WNV in Culex pipiens and the corresponding 95% credible interval across three different settings. The estimates are based on the life-history trait temperature response estimates presented in the main text and using the three different levels of between-experiment standard deviations presented before for both the mosquito infection probability and the extrinsic incubation period. Both the mean estimate as well as the 95% credible interval of the $R_0^{\mathrm{rel}}(T)$ temperature optimum is relatively stable across the three settings.

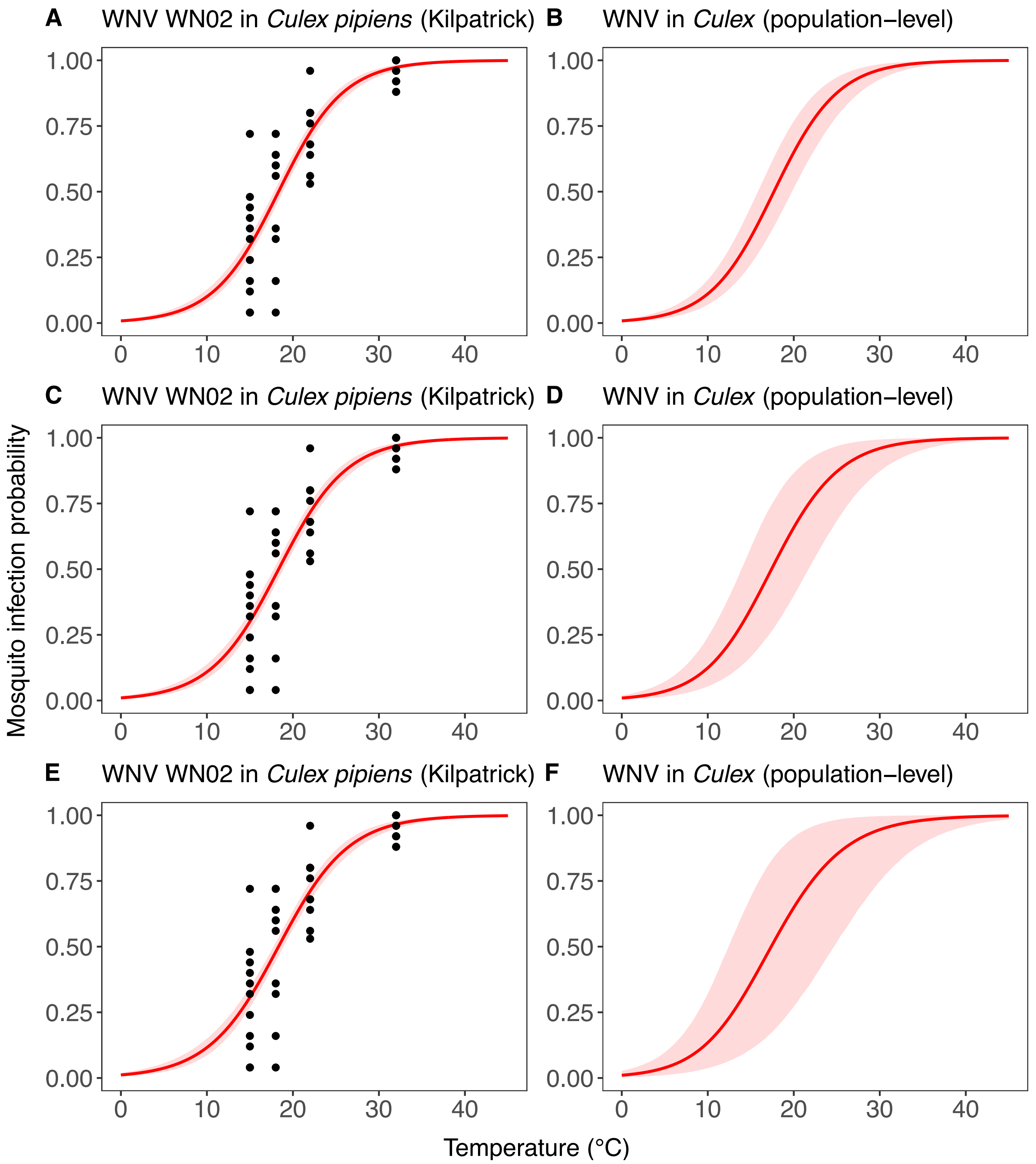


**Figure SI4.3.** (A, C, E) Estimates of the expected temperature response of the mosquito infection probability in the experiment by Kilpatrick et al. [16] on WNV WN02 in Cx. pipiens and (B, D, F) temperature response estimates at the population-level generated by sampling from the hierarchical prior determined by the population-level means and between-experiment standard deviations. The estimates are shown for three different levels of between-experiment standard deviations: (A-B) small, (C-D) moderate, and (E-F) large between-experiment variability. Black dots represent data from experimental studies. Red solid lines represent posterior distribution mean model fits. Red shaded areas represent the corresponding central/equal-tailed 95% credible interval.

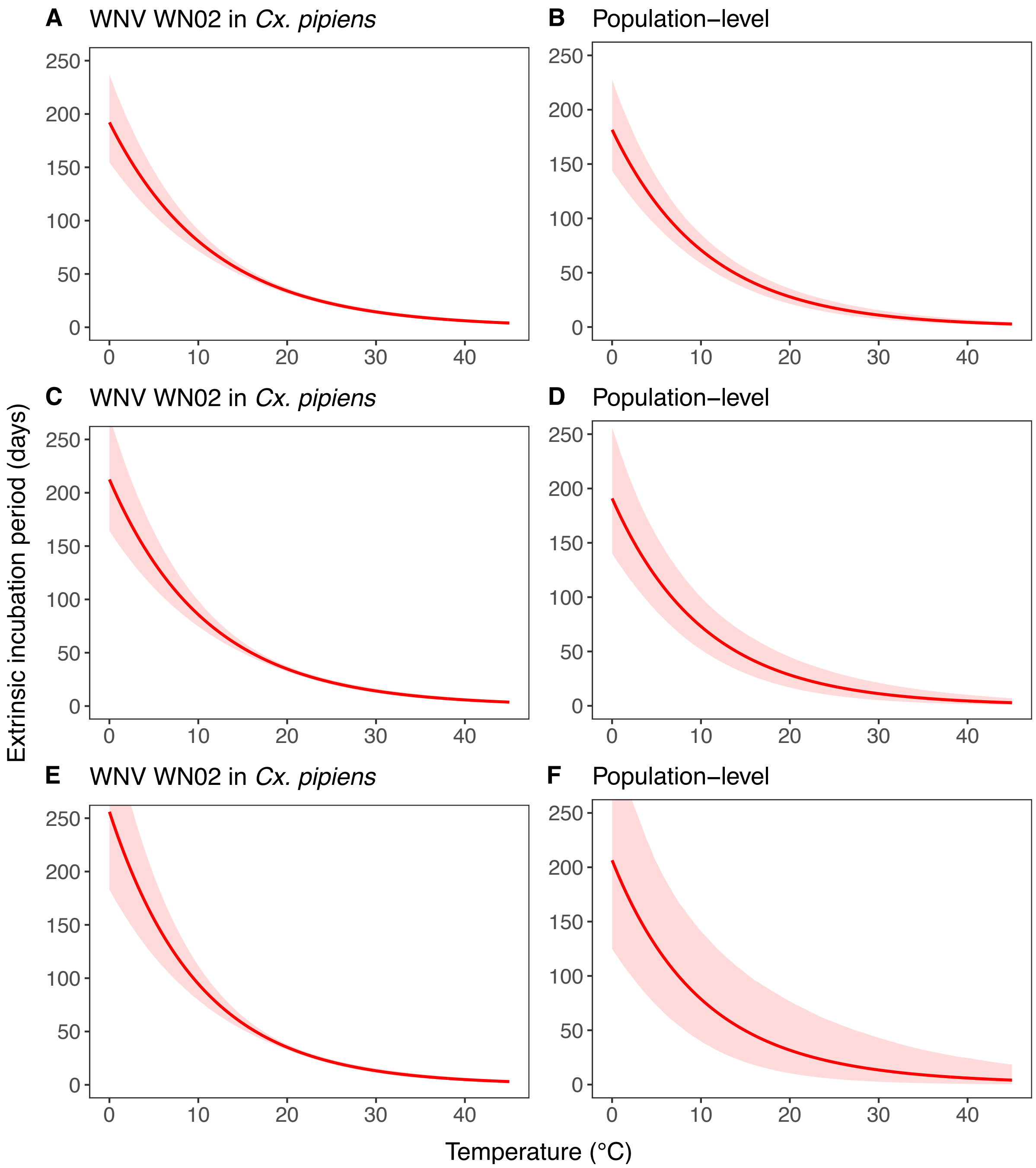


**Figure SI4.4.** (A, C, E) Estimates of the expected temperature response of the extrinsic incubation period in the experiment on WNV WN02 in Cx. pipiens and (B, D, F) temperature response estimates at the population-level generated by sampling from the hierarchical prior determined by the population-level means and between-experiment standard deviations. The estimates are shown for three different levels of between-experiment standard deviations: (A-B) small, (C-D) moderate (used in main text), and (E-F) large between-experiment variability. Black dots represent data from experimental studies. Red solid lines represent posterior distribution mean model fits. Red shaded areas represent the corresponding central/equal-tailed 95% credible interval.

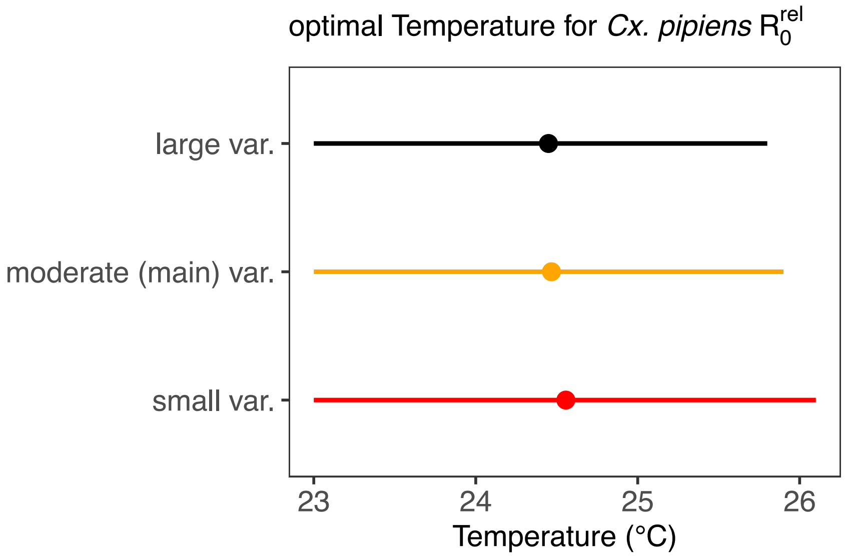


**Figure SI4.5.** Mean estimates of the optimal temperature of $R_0^{\mathrm{rel}}$ for WNV in *Culex* pipiens (dots) and the corresponding central/equal-tailed 95% credible interval (lines) based on the life-history trait temperature response estimates presented in the main text and pathogen-related traits estimated using three different levels of between-experiment standard deviations for both the mosquito infection probability and the extrinsic incubation period: small, moderate (used in main text), and large between-experiment variability.

### SI5 Importance of accounting for between-experiment variability

We fitted temperature-response functions to several mosquito-pathogen traits using a dataset that includes outcomes from experiments on different species with multiple experiments per species where available. Here, we demonstrate that it is crucial that the statistical model used for fitting the temperature response functions accounts for variability of temperature responses between experiments on the same species. The fact that, within a given trait and species, groups of datapoints come from the same experiment introduces statistical dependencies into the dataset. The sequence of datapoints along the temperature gradient of each experiment can be considered as evidencing a full temperature response curve, instead of each individual datapoint simply carrying information about trait performance at a specific temperature value. Figure SI5.1 illustrates this point with the Briére function fit to the data on juvenile development rate $\delta_J(T)$ of *Cx. pipiens* based on the hierarchical model introduced in the main text. The mean estimates for each experiment are shown as dashed lines coloured in accordance with the datapoints belonging to the respective experiment. The expected response of the species (mean (solid line) and 95% credible interval of the mean (shaded)) is shown in black and, heuristically speaking, results from the collective evidence given by the dashed curves. As can be seen from the datapoints in the Figure, the temperature response observed in each experiment systematically deviates from the expected response of the species (black solid line) along the temperature range. If the experiment identity would not impact the temperature response, the datapoints of each experiment would instead appear scattered randomly along the black curve.

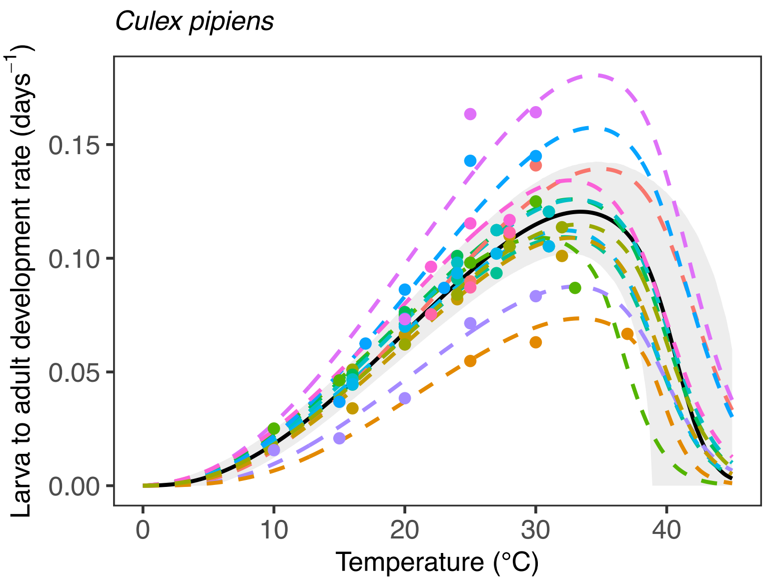


**Figure SI5.1.** Estimated temperature response of the larva to adult development rate of *Cx. pipiens*. Dots represent data from experimental studies and are coloured according to experiment identity. The coloured dashed lines are the posterior distribution mean model fits for each experiment on *Cx. pipiens*. The black solid line represents the posterior distribution mean model fit of the expected temperature response of *Cx. pipiens*. The grey shaded area represents the corresponding central/equal-tailed 95% credible interval.

Previous works often ignored these dependencies by completely pooling data across experiments [2,3,7,23,56]. We show that this can impact both the mean and the uncertainty of the expected temperature response of a species. We illustrate this by contrasting temperature response fits for juvenile development rate $\delta_J(T)$ derived from our full hierarchical model described in the main text and from a model that ignores between-experiment variability (essentially assuming $\sigma^{\text{exp}} = 0$).

Figure SI5.2A and SI5.2B contrast the model fit for *Cx. quinquefasciatus* with and without between-experiment variability. There are two striking differences. First, the model fit without between-experiment variability results in tighter uncertainty estimates for the expected temperature response of *Cx. quinquefasciatus*. This is because, if we presuppose no variability across experiments, we assume that each experiment carries much more information about the species' expected response. Second, the mean estimate of the expected temperature response deviates across the two models. Most notably, in the upper temperature range the model without experiment effect estimates that the development rate drops at lower temperatures. This happens because of the low trait values measured at the highest temperature points in the dataset. However, when the model accounts for dependencies, i.e., that these datapoints belong to experiments that estimated relatively low values along the whole temperature range (as revealed by colouring datapoints according to experiment identity) the model fit predicts that the development rate is increasing into higher temperature values (see Figure SI5.2B).

Another interesting effect is demonstrated by contrasting the model fits for *Ae. solicitans* with and without between-experiment variability (Figure SI5.2C and SI5.2D). For this species the dataset contains only data from a single experiment. Moreover, when compared to data from other species, the development rates measured for *Ae. solicitans* reach exceptionally high values. In our hierarchical model the between experiment variability $\sigma^{\text{exp}}$ is learned from the collective evidence of variabilities between experiments within all species, i.e., mostly learned from species that have data from several experiments available (like *Cx. pipiens* and *Cx. quinquefasciatus*). Partially pooling this between-experiment variability to *Ae. solicitans* via the hierarchical prior, the model assigns a high probability that the single experiment on this species represents an outlier and poorly represents its expected temperature response. This results in a large uncertainty for the expected temperature response of *Ae. solicitans* with a mean estimate that is predicted to be lower than indicated by the single experiment available. Given the large between-experiment variability seen for other species this is a logical result. Simply, believing that the single experiment is a good representation of the typical temperature response of *Ae. solicitans*, as done by the model without between-experiment variability, results in a model fit that closely follows the datapoints with very low uncertainty, despite the limited data available for this species (Figure SI5.2D).

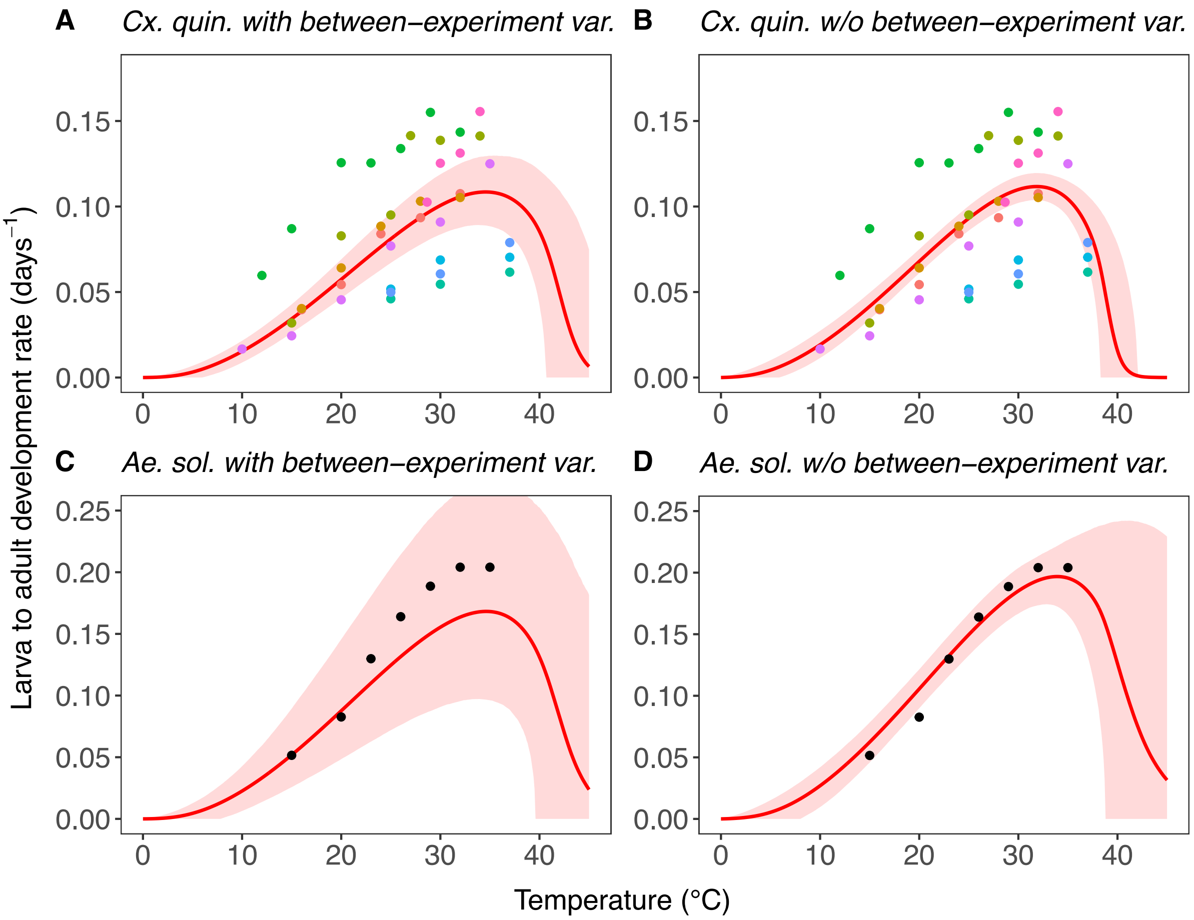


**Figure SI5.2.** Estimates of the expected temperature response of the larva to adult development rate of (A-B) *Cx. quinquefasciatus* and (C-D) *Ae. solicitans* contrasting fits derived from (A and C) the full hierarchical model and (B and D) a model neglecting between-experiment variability. Dots represent data from experimental studies and are coloured according to experiment identity. Red solid lines represent posterior distribution mean model fits. Red shaded areas represent the corresponding central/equal-tailed 95% credible interval.

### SI6 Taxonomic considerations

The taxonomy within the *Cx. pipiens* complex introduced challenges to our analysis. This complex encompasses several species including the species *Cx. pipiens* (mainly temperate) and *Cx. quinquefasciatus* (more tropical). In regions where the distribution range of *Cx. pipiens* and *Cx. quinquefasciatus* overlap the boundaries between the species get blurry with different degrees of hybridization appearing. For example, there is a recognized subspecies of *Cx. pipiens* called *Cx. pipiens pallens* which is a *pipiens* and *quinquefasciatus* hybrid that occurs in Northeast Asia [57]. In addition, *Cx. pipiens* is divided into two ecotypes namely *Cx. pipiens pipiens* and *Cx. pipiens molestus* which are morphologically indistinguishable but have distinct ecological characteristics and are hypothesized to fulfil unique roles in WNV transmission because of differences in host biting preferences [58]. Also at this taxonomic level, hybrids of the *pipiens* forms *pipiens* and *molestus* occur to varying degrees and display their own ecological characteristics. This situation leads to a complex ecological mosaic that makes predictions of vectorial capacity notoriously complicated [58]. In our analysis we investigate potential differences in the temperature responses of all these mosquitoes. Strictly speaking, this introduces an inaccuracy because we model their temperature responses with a single hierarchical prior that is then shared between species, but also subspecies, and ecotypes ignoring potential additional dependencies the lower levels of the taxonomy. Additionally, in several articles that contributed to our dataset it is only stated that the experiments were conducted on mosquitoes from the *Cx. pipiens* complex without further specification of the species (e.g., [35]). In some cases, we could allocate the data to the most likely species based on the geographical origin of the mosquitoes or based on names of lab strains [6,33,37], but other data might represent a mix of *Cx. quinquefasciatus* and *pipiens* [35]. In addition, data that was specified to the species *Cx. pipiens* seldomly specified the ecotype. We allocate this uncertain data to the fits labelled as *Cx. pipiens* accepting the uncertainty of a potential introgression by *Cx. quinquefasciatus* or *Cx. pipiens molestus*. Through a comparison of the fits labelled as *Cx. pipiens* to the fits specifically derived for *Cx. quinquefasciatus* and *Cx. pipiens molestus* we can at least have an indication of whether their temperature responses diverge. For simplicity we refer to the all the different mosquitoes in the *Cx. pipiens* complex discussed above as different species throughout the manuscript, bearing in mind that this is taxonomically inaccurate.

### SI7 Alternative mosquito abundance models

We incorporate the effect of temperature on mosquito abundance in the $R_0^{\mathrm{rel}}(T)$ model by deriving an equilibrium expression from a temperature-driven mosquito population dynamics model. In the literature, a variety of such models have been proposed. These often differ in the way in which competition within the mosquito population is described. The model that we introduce in the main text includes a quadratic mortality term to account for competition between juvenile mosquitoes in the aquatic breeding habitat, a mechanism often observed in experimental studies [26,37,38,45]. While this choice is well supported by the available knowledge, we consider also other model formulations that can be found in the literature. Below we introduce a collection of alternative model formulations. From each model we derive an equilibrium expression which we integrate into $R_0^{\mathrm{rel}}(T)$ to see if the choice of mosquito population dynamics model might influence our results on the temperature-sensitivity of transmission suitability (Figure 4 in the main text). We also contrast these models to a mosquito abundance expression utilized by earlier works on WNV temperature suitability modelling [7,23,39,56] and discuss below the lack of theoretical foundation of this expression.

**Alternative model 1**

In this model population growth is also limited by a quadratic competition-driven mortality term. In contrast to the model in the main text where the term governing competition is assumed to be temperature independent, here the competition-driven mortality follows the same temperature-dependence as the natural mortality rate. The differential equation and the corresponding adult female mosquito equilibrium are given by:

$$\begin{aligned} \dot{E} &= \beta(T)M - \delta_E(T)E - \mu_E(T)E \\ \dot{J} &= \delta_E(T)E - \left(1+\frac{J}{K}\right)\mu_J(T)J - \delta_J(T)J \\ \dot{M} &= \omega\delta_J(T)J - \mu_M(T)M \end{aligned} \qquad (SI7.1)$$

$$M_1^*(T) = K\frac{\omega^2\beta(T)p_E(T)\delta_J(T)^2}{\mu_M(T)^2\mu_J(T)}\left[1-\frac{\mu_M(T)}{\omega\beta(T)p_{EJ}(T)}\right] \qquad (SI7.2)$$

Similar terms for competition-driven mortality in the aquatic phase can be found in the literature [59,60], although often incorporated into extended versions of the model above.

**Alternative model 2**

This model neglects any direct impacts of increased juvenile density on juvenile individuals. Instead, it is assumed that juvenile crowding in breeding sites results in a reduction of egg laying by female adult mosquitoes, an assumption that was for example used in the models by Liu-Helmersson et al. [61] and Yang et al. [62]. Our model equations and equilibrium expression adopting this mechanism read:

$$\begin{aligned} \dot{E} &= \beta(T)\left(1-\frac{J}{K}\right)M - \delta_E(T)E - \mu_E(T)E \\ \dot{J} &= \delta_E(T)E - \mu_J(T)J - \delta_J(T)J \\ \dot{M} &= \omega\delta_J(T)J - \mu_M(T)M \end{aligned} \qquad (SI7.3)$$

$$M_2^*(T) = K\frac{\omega\delta_J(T)}{\mu_M(T)}\left[1-\frac{\mu_M(T)}{\omega\beta(T)p_{EJ}(T)}\right] \qquad (SI7.4)$$

**Alternative model 3**

Several models in the literature assume that high density of adult mosquitoes is limiting population growth [63,64]. In a broader sense, we incorporate this assumption into the following model and equilibrium expression by viewing adult mosquitoes as competing for breeding sites available for egg laying:

$$\begin{aligned} \dot{E} &= \beta(T)\left(1-\frac{M}{K}\right)M - \delta_E(T)E - \mu_E(T)E \\ \dot{J} &= \delta_E(T)E - \mu_J(T)J - \delta_J(T)J \\ \dot{M} &= \omega\delta_J(T)J - \mu_M(T)M \end{aligned} \qquad (SI7.5)$$

$$M_3^*(T) = K\left[1-\frac{\mu_M(T)}{\omega\beta(T)p_{EJ}(T)}\right] \qquad (SI7.6)$$

**Alternative model 4**

Some earlier works on WNV temperature suitability modelling [7,23,39,56] adopted an expression that in terms of the parameters introduced in our work could be expressed as:

$$M_4^*(T) = \frac{\beta(T)p_{EJ}(T)\delta_J(T)}{\mu_M(T)^2} \qquad (SI7.7)$$

Several other works have also used variations of this expression to capture the temperature suitability for mosquito abundance (see e.g., [2–5,8,65]). Similar as for the model in the main text and alternative models 1-3, this expression can be traced back to an equilibrium derived from a dynamical model [66]. The derived equilibrium expression was later modified by Mordecai et al. [65]. We found that the original model formulation [66] has an inconsistency in dimensionality. The model incorporated an expression for the total female adult mosquito birth rate that has dimensions [1/time]. In terms of dimensions this term could only act as a per capita birth rate but not as a total birth rate, which for consistency of the differential equations would need to have the dimensions [individuals/time]. After further calculations, this dimensionality inconsistency resulted in an equilibrium expression of the form “birth rate divided by death rate”. Apart from the fact that the incorporated birth rate does not have the right dimensionality for such an expression, equilibrium expressions of this form appear in models that assume a total birth rate that does not scale with population size, which by itself would be a questionable assumption in a mosquito population model. Therefore, it appears that the frequently used expression $(SI7.7)$ lacks a connection to a mosquito population dynamics model and therefore a proper model derivation. Outlining such a connection is crucial to make the assumptions transparent that underly mosquito population suitability models.

### SI8 Key changes applied to original dataset

The starting point of our data collection is the dataset compiled by Shocket et al. [7]. Before we extended this dataset by data from additional and more recent experimental studies, we double checked each entry in the original dataset with the article describing the corresponding experiment. Moreover, for some traits we also revised the conceptual approach. In the following we outline all key changes.

We corrected any inconsistencies that we found between dataset and original articles. For the study by Shelton et al. [25], the data noted down for *Cx. salinarius* and *Ae. solicitans* did not match with results shown in the Figures of the original article. In case of *Cx. salinarius*, for example, it seems to us that data was accidently extracted from the Figure for *Cx. quinquefasciatus*. In case of the study by Tekle et al. [13], data on juvenile survival of *Cx. pipiens* was incorrectly extracted from the corresponding Table. The data shown in Figure 4 in the study by Cornel et al. [20] was classified as (# transmitting mosquitoes/# exposed mosquitoes) although it is representing (# transmitting mosquitoes/# mosquitoes with infected body). In addition, the data shown in Figure 4 in the study by Reisen et al. [19] was classified as (# transmitting mosquitoes/# mosquitoes with disseminated infection) although it is representing (# transmitting mosquitoes/# mosquitoes with infected body).

We removed datapoints when they, upon our interpretation of the experiments, did not represent the traits as defined in Shocket et al. [7] or our work, respectively. This concerns biting rates from Ruybal et al. [36] and the observations on juvenile survival at 10°C in the study by Mogi [11]. The latter were obtained from experiments where at the end of the observation period a part of the sample population had neither developed nor died yet and we therefore judged them as insufficient to deduce juvenile development rate or survival data [11].

We removed datapoints from studies that were conducted under experimental setups that we do not view comparable to the other studies included in the dataset. Egg viability data in Rayah et al. [67] was derived by exposing eggs from *Cx. quinquefasciatus* to a given temperature for 24 hours but afterwards returned to room temperature instead of testing the effect of continuous exposure to the same temperature. We also removed the egg viability data from Group B in van der Linde et al. [51] since here eggs were exposed to room temperature for the first 12 hours after egg laying and only afterwards exposed to other temperature settings.

In the list of articles used by Shocket et al. [7] we also found some additional datapoints that were not recorded in the original dataset. This encompasses data on *Cx. pipiens* juvenile development rate for the “high elevation population” in Ruybal et al. [36] and data on juvenile development rate and survival of *Cs. inornata* from Brust and Buth et al. [26,44]. We incorporated these data in our analysis.

We recalculated mean adult mosquito lifespan from raw data shared by Ruybal et al. [36] since we could not reconstruct how the values given in the original dataset [7] were calculated from the data shown in the Figure in the primary articles’ supplementary information.

As explained in SI6 the complicated taxonomy within the *Culex pipiens* complex introduces challenges to any multi-species analysis including ours. While we aimed to derive temperature response functions for each (sub-)species/ecotype within the complex, the taxonomic order was not consistently treated in the original dataset [7]. Some of the trait data of *Cx. pipiens pallens* and *Cx. pipiens molestus* was exclusively used to generated empirical priors in this previous work [7]. This data was accordingly labelled as “Cpal” or “Cmol” in the original dataset. However, the previous analysis also includes examples where data from these mosquitoes was directly used to fit trait performance for *Culex pipiens* and were in these cases given the label “Cpip” in the dataset [7]. We aimed for a more consistent approach and used all data that was clearly identified as belonging to *Cx. pipiens pallens* or *Cx. pipiens molesuts* to estimate response functions for this *Cx. pipiens* subspecies and ecotype, respectively.

Whilst most of the data to fit juvenile traits in the original dataset [7] represents the larva and pupa stage together, there are examples where the reported data included the egg stage [29], excluded the pupa stage [43,68], was measured from the third larva instar to adult emergence [69], or measured the time to first adult emergence instead of a summary statistic [49]. To avoid mixing such heterogeneous data, we clearly separate the aquatic juvenile phase into egg stage and a juvenile stage, combining larvae and pupae, and only kept data representing these phases adequately. The only exception to this rule represents one study [6] where juvenile traits were measured starting from the second instar larva stage. In some cases juvenile development times were reported for male and female mosquitoes separately [30,43]. In these cases, we averaged the values for male and female juvenile mosquitoes.

As reasoned in detail in SI1, we agree with the recommendation of Von Schmalensee et al. [9] that 100% end mortality in juvenile development stages should not be used as an indication of a development rate of zero. Therefore, we removed any of these artificial datapoints in the original dataset (therein stored as 1000 days development time). Following the same reasoning, we remove data indicating a biting rate of zero (stored as 1000 days gonotrophic cycle duration) that were motivated from observations where adult mosquitoes did not survive the time from taking a blood meal to successfully laying eggs.

Some experimental studies varied additional factors to temperature such as larva density, larva food concentration, or insecticide concentration. In these cases, we only collected data from the lowest larva density, highest food settings, and control settings in case of insecticides to minimize the influence of juvenile competition and focus on the effects of temperature. In contrast, the original dataset [7] in some cases included trait performance across multiple settings, i.e., not only varying temperature [30,42,45].

For egg viability we confined the data to species from the genus *Culex*. We excluded data that was available for temperate *Aedes* species since for these species exposure to low temperatures can initiate diapause in the egg stage [70,71], which can make it hard to distinguish diapausing eggs (which could hatch when returned to suitable temperatures) from eggs that ultimately failed to hatch. Only the latter would be directly comparable to the observations made on *Culex* egg viability.

Our analytical approach to modelling vector competence and the extrinsic incubation period differs quite distinctly to the analysis by Shocket et al. [7] (see SI1). Therefore, we collected the data necessary to fit the temperature response of these traits (described in SI1) from scratch from the available literature identified through the original dataset [7]. Shocket et al. [7] derived the temperature response of the extrinsic incubation period of WNV in *Cx. quinquefasciatus* based on data displayed in Figure 1 (d) in the study by Paull et al. [72] that to the best of our knowledge is otherwise unpublished. In this form the data cannot be incorporated into our model for the extrinsic incubation period.

Therefore, we could not derive the temperature response of the extrinsic incubation for WNV in *Cx. quinquefasciatus*. We limited our analysis to vector competence data for WNV, while the original dataset [7] included data for other viruses.

### SI9 Overview of collected data

**Table SI9.1.** Characteristics of included laboratory experimental studies. We characterized mosquito populations that spend less than five generations in the laboratory before experimentation as having field origin. In cases where the time the mosquito population spent in the laboratory could not be identified, we classified the mosquito origin as unclear. *Studies that extend the original dataset [7]. Changes applied to previously included studies are listed in SI8.

| Study | Species | Traits | Detail of reporting at juvenile stage | Temperatures tested | Mosquito origin |
|---|---|---|---|---|---|
| Andreadis et al. (Greece, 2014) | *Cx. pipiens* | $\mathrm{lf}$ | NA | 15, 20, 25, 27.5, 30 | Field |
| Boerlijst et al. (Netherlands, 2022)* | *Cx. pipiens* | $\delta_J, p_J$ | Larva, larva-adult | 20, 25, 30 | Field |
| Brust (Canada, 1967) | *Cs. melanura*<br>*Ae. vexans*<br>*Ae. nigromaculis* | $p_J$ | Larva-adult | 5, 10, 15.5, 21, 26.5, 32 | Both |
| Buth et al. (Canada, 1990) | *Cx. tarsalis*<br>*Cx. restuans*<br>*Cs. inornata* | $\delta_J, p_J$ | Larva-adult | 15, 20, 25 | Lab |
| Ciota et al. (USA, 2014) | *Cx. pipiens*<br>*Cx. quinquefasciatus*<br>*Cx. restuans* | $\delta_J, p_J$, $\mathrm{lf}$ | Larva-adult | 16, 20, 24, 28, 32 | Both |
| Cornel et al. (South Africa, 1993) | *Cx. univittatus* | $\mathrm{EIP}$ | NA | 14, 18, 26, 30 | Both |
| Dodson et al. (USA, 2012) | *Cx. tarsalis* | $p_J$ | Larva-adult | 19, 25, 31 | Unclear |
| Fay et al. (USA, 2024)* | *Cx. pipiens* | $\beta_{\mathrm{ER}}, \delta_J$, $p_J, \mathrm{lf}$ | Larva-adult | 22, 25, 28 | Field |
| Dohm et al. (USA, 2002) | *Cx. pipiens* | $b_M$ | NA | 18, 20, 26, 30 | Field |
| Gunay et al. (Turkey, 2010)* | *Cx. quinquefasciatus* | $p_J$ | Larva-adult | 15, 20, 23, 27, 30 | Lab |
| Kilpatrick et al. (USA, 2008) | *Cx. pipiens* | $b_M, \mathrm{EIP}$ | NA | 15, 18, 22, 32 | Lab |
| Kiarie-Makara et al. (Korea, 2015) | *Cx. pipiens pallens*<br>*Cx. pipiens molestus* | $\delta_E$, $\delta_J, \mathrm{lf}$ | Larva, pupa | 20, 24, 28 | Lab |

| | | | | | |
|---|---|---|---|---|---|
| Loetti et al. (Argentina, 2011) | *Cx. pipiens* | $\delta_J, p_J$ | Instars, pupa, larva-adult | 7, 10, 15, 20, 25, 30, 33 | Field |
| Li et al. (China, 2019) | *Cx. pipiens pallens* | $a$, $\delta_E$, $\delta_J, p_E$, $p_J$ | Larva, pupa | 10, 15, 20, 25, 30, 35, 40 | Lab |
| Madder et al. (Canada, 1983) | *Cx. pipiens*<br>*Cx. restuans* | $a$, $\delta_E$, $\delta_J, p_J$ | Larva-adult | 15, 17, 20, 22, 23, 25, 30 | Both |
| Mahmood and Crans (USA, 1997) | *Cs. melanura* | $a$ | NA | 10, 16, 22, 28 | Lab |
| Mahmood (USA, 1998) | *Cs. melanura* | $\delta_E$, $\delta_J$, $p_J$ | Instars, pupa, larva-adult | 10, 16, 22, 28, 32, 34 | Lab |
| Mogi (Japan, 1992) | *Cx. quinquefasciatus*<br>*Cx. pipiens pallens* | $\beta_{\mathrm{ER}}, p_J$ | Larva-adult | 15, 20, 28 | Unclear |
| Mpho (UK, 2002) | *Cx. quinquefasciatus* | $\delta_J, p_J$ | Larva-adult | 25, 30, 37 | Both |
| Mpho (UK, 2002) | *Cx. pipiens* | $\delta_J, p_J$ | Larva-adult | 25, 30, 37 | Field |
| Muturi et al. (USA, 2011) | *Cx. restuans* | $\delta_J, p_J$ | Larva-adult | 20, 25, 30 | Field |
| Nayar (USA, 1972) | *Ae. taeniorhynchus* | lf | NA | 22, 27, 32 | Unclear |
| Oda (Japan, 1980) | *Cx. quinquefasciatus*<br>*Cx. pipiens molestus* | $\beta_{\mathrm{ER}}, p_E$ | NA | 21, 25, 28, 30 | Unclear |
| Oda (Japan, 1999) | *Cx. quinquefasciatus*<br>*Cx. pipiens molestus* | $p_J$, lf | Larva, larva-adult | 21, 25, 30 | Both |
| Oda (Japan, 2002)* | *Cx. quinquefasciatus*<br>*Cx. pipiens pallens* | lf | NA | 25, 30 | Both |
| Olayemi (Nigeria, 2016)* | *Cx. quinquefasciatus* | $p_J$ | Larva, pupa | 28, 30, 32, 34 | Lab |
| Ojenícek and Gelbic (Czech Republic, 2000) | *Cx. pipiens molestus* | $\delta_J, p_J$ | Larva-adult | 15, 22, 30 | Lab |
| Reisen et al. (USA, 1992) | *Cx. quinquefasciatus*<br>*Cx. tarsalis* | $a$ | NA | 10, 15, 20, 25, 30 | Lab |
| Reisen (USA, 1995) | *Cx. tarsalis* | $\delta_J$, lf | Larva-adult | 14, 15, 20, 25, 26, 30, 32, 35 | Field |
| Reisen et al. (USA, 2006) | *Cx. tarsalis* | EIP | NA | 14, 18, 22, 26, 30 | Lab |
| Rueda et al. (USA, 1990) | *Cx. quinquefasciatus* | $\delta_J, p_J$ | Instars, pupa, larva-adult | 15, 20, 25, 27, 30, 34 | Lab |
| Ruybal et al. (USA, 2016) | *Cx. pipiens* | $\delta_J, p_J$, lf | Larva-adult | 16, 20, 24, 27, 31, 35 | Field |

| | | | | | |
|---|---|---|---|---|---|
| Spanoudis et al. (Greece, 2019)* | *Cx. pipiens molestus* | $\delta_E, \delta_J$, $p_E, p_J$, lf | Instars, larva, pupa, larva-adult | 15, 17.5, 20, 22.5, 25, 27.5, 30, 32.5 | Lab |
| Shelton (USA, 1973) | *Cs. inornata*<br>*Cx. restuans*<br>*Cx. quinquefasciatus*<br>*Cx. salinarius*<br>*Ae. sollicitans*<br>*Ae. triseriatus* | $\delta_J, p_J$ | Instars, pupa, larva-adult | 12, 15, 20, 23, 26, 29, 32, 35 | Both |
| Shriver and Bickley (USA, 1964)* | *Cx. quinquefasciatus* | $\delta_E, p_E$ | NA | 7.2, 10, 15.6, 18.3, 23.9, 29.4, 35, 37.8 | Lab |
| Tekle (USA, 1960) | *Cx. pipiens*<br>*Cx. quinquefasciatus* | $a, \delta_J$, $p_J$ | Larva-adult | 10, 15, 20, 25, 30, 35 | Unclear |
| Teng and Apperson (USA, 2000) | *Ae. triseriatus* | $p_J$ | Larva-adult | 15, 23, 31 | Field |
| Ukubuiwe (Nigeria, 2018)* | *Cx. quinquefasciatus* | $\delta_J, p_J$ | Larva, pupa, larva-adult | 28.66, 30, 32, 34 | Lab |
| Van der Linde (South Africa, 1990) | *Cx. theileri* | $p_E$ | NA | 6, 9, 12, 15, 18, 21, 24, 27, 30, 33, 36, 39, 42 | Unclear |
| Zayed et al. (Egypt, 2019)* | *Cx. pipiens* | $\delta_E, \delta_J$ | Larva, pupa | 20, 25, 30 | Lab |

### References


1. Briere J-F, Pracros P, Le Roux A-Y, Pierre J-S. A Novel Rate Model of Temperature-Dependent Development for Arthropods. Environ Entomol. 1999;28: 22–29. doi:10.1093/ee/28.1.22

2. Villena OC, Ryan SJ, Murdock CC, Johnson LR. Temperature impacts the environmental suitability for malaria transmission by *Anopheles gambiae* and *Anopheles stephensi*. Ecology. 2022;103. doi:10.1002/ecy.3685

3. Mordecai EA, Cohen JM, Evans MV, Gudapati P, Johnson LR, Lippi CA, et al. Detecting the impact of temperature on transmission of Zika, dengue, and chikungunya using mechanistic models. Althouse B, editor. PLoS Negl Trop Dis. 2017;11: e0005568. doi:10.1371/journal.pntd.0005568

4. Shocket MS, Ryan SJ, Mordecai EA. Temperature explains broad patterns of Ross River virus transmission. eLife. 2018;7: e37762. doi:10.7554/eLife.37762

5. Mordecai EA, Caldwell JM, Grossman MK, Lippi CA, Johnson LR, Neira M, et al. Thermal biology of mosquito-borne disease. Byers J (Jeb), editor. Ecol Lett. 2019;22: 1690–1708. doi:10.1111/ele.13335

6. Mpho M, Callaghan A, Holloway GJ. Effects of temperature and genetic stress on life history and fluctuating wing asymmetry in Culex pipiens mosquitoes. Eur J Entomol. 2002;99: 405–412. doi:10.14411/eje.2002.050

7. Shocket MS, Verwillow AB, Numazu MG, Slamani H, Cohen JM, El Moustaid F, et al. Transmission of West Nile and five other temperate mosquito-borne viruses peaks at temperatures between 23°C and 26°C. eLife. 2020;9: e58511. doi:10.7554/eLife.58511

8. Johnson LR, Ben-Horin T, Lafferty KD, McNally A, Mordecai E, Paaijmans KP, et al. Understanding uncertainty in temperature effects on vector-borne disease: a Bayesian approach. Ecology. 2015;96: 203–213. doi:10.1890/13-1964.1

9. Von Schmalensee L, Hulda Gunnarsdóttir K, Näslund J, Gotthard K, Lehmann P. Thermal performance under constant temperatures can accurately predict insect development times across naturally variable microclimates. Ghalambor C, editor. Ecology Letters. 2021;24: 1633–1645. doi:10.1111/ele.13779

10. Jia P, Lu L, Chen X, Chen J, Guo L, Yu X, et al. A climate-driven mechanistic population model of Aedes albopictus with diapause. Parasites Vectors. 2016;9: 175. doi:10.1186/s13071-016-1448-y

11. Mogi M. Temperature and Photoperiod Effects on Larval and Ovarian Development of New Zealand Strains of Culex quinquefasciatus (Diptera: Culicidae). Annals of the Entomological Society of America. 1992;85: 58–66. doi:10.1093/aesa/85.1.58

12. Oda T, Mori A, Ueda M. Effects of temperatures on the oviposition and hatching of eggs in culex pipiens molestus and culex pipiens quinquefasciatus. Tropical Medicine. 1980;22: 167–180.

13. Tekle A. The Physiology of Hibernation and Its Role in the Geographical Distribution of Populations of the Culex Pipiens Complex. The American Journal of Tropical Medicine and Hygiene. 1960;9: 321–330. doi:10.4269/ajtmh.1960.9.321

14. Mahmood F, Crans WJ. A thermal heat summation model to predict the duration of the gonotrophic cycle of Culiseta melanura in nature. J Am Mosq Control Assoc. 1997;13: 92–94.

15. Vogels CB, Göertz GP, Pijlman GP, Koenraadt CJ. Vector competence of European mosquitoes for West Nile virus. Emerging Microbes & Infections. 2017;6: 1–13. doi:10.1038/emi.2017.82

16. Kilpatrick AM, Meola MA, Moudy RM, Kramer LD. Temperature, Viral Genetics, and the Transmission of West Nile Virus by Culex pipiens Mosquitoes. Buchmeier MJ, editor. PLoS Pathog. 2008;4: e1000092. doi:10.1371/journal.ppat.1000092

17. Dohm DJ, O'Guinn ML, Turell MJ. Effect of Environmental Temperature on the Ability of *Culex pipiens* (Diptera: Culicidae) to Transmit West Nile Virus. J Med Entomol. 2002;39: 221–225. doi:10.1603/0022-2585-39.1.221

18. Brady OJ, Golding N, Pigott DM, Kraemer MUG, Messina JP, Reiner Jr RC, et al. Global temperature constraints on Aedes aegypti and Ae. albopictus persistence and competence for dengue virus transmission. Parasit Vectors. 2014;7: 338. doi:10.1186/1756-3305-7-338

19. Reisen WK, Fang Y, Martinez VM. Effects of Temperature on the Transmission of West Nile Virus by *Culex tarsalis* (Diptera: Culicidae). J Med Entomol. 2006;43: 309–317. doi:10.1093/jmedent/43.2.309

20. Cornel AJ, Jupp PG, Blackburn NK. Environmental Temperature on the Vector Competence of Culex univittatus (Diptera: Culicidae) for West Nile Virus. Journal of Medical Entomology. 1993;30: 449–456. doi:10.1093/jmedent/30.2.449

21. Shapiro LLM, Whitehead SA, Thomas MB. Quantifying the effects of temperature on mosquito and parasite traits that determine the transmission potential of human malaria. Schneider D, editor. PLoS Biol. 2017;15: e2003489. doi:10.1371/journal.pbio.2003489

22. Reisen WK, Meyer RP, Presser SB, Hardy JL. Effect of Temperature on the Transmission of Western Equine Encephalomyelitis and St. Louis Encephalitis Viruses by Culex tarsalis (Diptera: Culicidae). Journal of Medical Entomology. 1993;30: 151–160. doi:10.1093/jmedent/30.1.151

23. Di Pol G, Crotta M, Taylor RA. Modelling the temperature suitability for the risk of West Nile Virus establishment in European *Culex pipiens* populations. Transbounding Emerging Dis. 2022;69. doi:10.1111/tbed.14513

24. Lambrechts L, Paaijmans KP, Fansiri T, Carrington LB, Kramer LD, Thomas MB, et al. Impact of daily temperature fluctuations on dengue virus transmission by *Aedes aegypti*. Proc Natl Acad Sci USA. 2011;108: 7460–7465. doi:10.1073/pnas.1101377108

25. Shelton R. The effect of temperatures on development of eight mosquito species. Mosquito News. 1973;33: 1–12.

26. Buth JL, Brust RA, Ellis RA. Development time, oviposition activity and onset of diapause in Culex tarsalis, Culex restuans and Culiseta inornata in southern Manitoba. J Am Mosq Control Assoc. 1990;6: 55–63.

27. Mahmood F, Crans WJ. Effect of Temperature on the Development of Culiseta melanura (Diptera: Culicidae) and its Impact on the Amplification of Eastern Equine Encephalomyelitis Virus in Birds. Journal of Medical Entomology. 1998;35: 1007–1012. doi:10.1093/jmedent/35.6.1007

28. Spanoudis CG, Andreadis SS, Tsaknis NK, Petrou AP, Gkeka CD, Savopoulou–Soultani M. Effect of Temperature on Biological Parameters of the West Nile Virus Vector Culex

pipiens form ‘molestus’ (Diptera: Culicidae) in Greece: Constant vs Fluctuating Temperatures. Journal of Medical Entomology. 2019;56: 641–650. doi:10.1093/jme/tjy224

29. Kiarie-Makara M, Ngumbi P, Lee D-K. Effects of temperature on the growth and development of culex pipiens Complex Mosquitoes (Diptera: Culicidae). Journal of Pharmacy and Biological Sciences. 2015;10: 1–10. doi:https://doi.org/10.9790/3008-10620110

30. Olejnícek J, Gelbic I. Differences in response to temperature and density between two strains of the mosquito, Culex pipiens molestus forskal. J Vector Ecol. 2000;25: 136–145.

31. Li J, Tang J, Zhu G, Yang M, Zhou H, Zhang M, et al. Effect of temperature on development and reproduction of three kind of mosquitoes. China Tropical Medicine. 2019;19: 939–943. doi:https://doi.org/10.13604/j.cnki.46-1064/r.2019.10.08

32. B. Zayed A. Influence of Temperature Change on the Growth and Susceptibility of the Common House Mosquito, <i>Culex pipiens</i> in Egypt to Some Insecticides. IJEE. 2019;4: 42. doi:10.11648/j.ijee.20190402.11

33. Mpho M, Callaghan A, Holloway GJ. Temperature and genotypic effects on life history and fluctuating asymmetry in a field strain of Culex pipiens. Heredity. 2002;88: 307–312. doi:10.1038/sj.hdy.6800045

34. Ciota AT, Matacchiero AC, Kilpatrick AM, Kramer LD. The Effect of Temperature on Life History Traits of *Culex* Mosquitoes. J Med Entomol. 2014;51: 55–62. doi:10.1603/ME13003

35. Loetti V, Schweigmann N, Burroni N. Development rates, larval survivorship and wing length of *Culex pipiens* (Diptera: Culicidae) at constant temperatures. Journal of Natural History. 2011;45: 2203–2213. doi:10.1080/00222933.2011.590946

36. Ruybal JE, Kramer LD, Kilpatrick AM. Geographic variation in the response of Culex pipiens life history traits to temperature. Parasites Vectors. 2016;9: 116. doi:10.1186/s13071-016-1402-z

37. Madder DJ, Surgeoner GA, Helson BV. Number of Generations, Egg Production, and Developmental Time of Culex Pipiens and Culex Restuans (Diptera: Culicidae) in Southern Ontario1. Journal of Medical Entomology. 1983;20: 275–287. doi:10.1093/jmedent/20.3.275

38. Boerlijst SP, Johnston ES, Ummels A, Krol L, Boelee E, Van Bodegom PM, et al. Biting the hand that feeds: Anthropogenic drivers interactively make mosquitoes thrive. Science of The Total Environment. 2023;858: 159716. doi:10.1016/j.scitotenv.2022.159716

39. Fay RL, Cruz-Loya M, Keyel AC, Price DC, Zink SD, Mordecai EA, et al. Population-specific thermal responses contribute to regional variability in arbovirus transmission with changing climates. iScience. 2024;27: 109934. doi:10.1016/j.isci.2024.109934

40. Rueda LM, Patel KJ, Axtell RC, Stinner RE. Temperature-Dependent Development and Survival Rates of Culex quinquefasciatus and Aedes aegypti (Diptera: Culicidae). Journal of Medical Entomology. 1990;27: 892–898. doi:10.1093/jmedent/27.5.892

41. Ukubuiwe AC, Olayemi IK, Arimoro FO, Omalu ICJ, Baba BM, Ukubuiwe CC, et al. Influence of rearing-water temperature on life stages' vector attributes, distribution and utilisation of metabolic reserves in Culex quinquefasciatus (Diptera: Culicidae): implications for disease transmission and vector control. JoBAZ. 2018;79: 32. doi:10.1186/s41936-018-0045-3

42. Muturi EJ, Lampman R, Costanzo K, Alto BW. Effect of Temperature and Insecticide Stress on Life-History Traits of Culex restuans and Aedes albopictus (Diptera: Culicidae). jnl med entom. 2011;48: 243–250. doi:10.1603/ME10017

43. Reisen WK. Effect of Temperature on Culex tarsalis (Diptera: Culicidae) from the Coachella and San Joaquin Valleys of California. Journal of Medical Entomology. 1995;32: 636–645. doi:10.1093/jmedent/32.5.636

44. Brust RA. WEIGHT AND DEVELOPMENT TIME OF DIFFERENT STADIA OF MOSQUITOES REARED AT VARIOUS CONSTANT TEMPERATURES. Can Entomol. 1967;99: 986–993. doi:10.4039/Ent99986-9

45. Teng H-J, Apperson CS. Development and Survival of Immature *Aedes albopictus* and *Aedes triseriatus* (Diptera: Culicidae) in the Laboratory: Effects of Density, Food, and Competition on Response to Temperature. J Med Entomol. 2000;37: 40–52. doi:10.1603/0022-2585-37.1.40

46. Oda T, Uchida K, Mori A, Mine M, Eshita Y, Kurokawa K, et al. Effects of high temperature on the emergence and survival of adult Culex pipiens molestus and Culex quinquefasciatus in Japan. J Am Mosq Control Assoc. 1999;15: 153–156.

47. Gunay F, Alten B, Ozsoy ED. Estimating reaction norms for predictive population parameters, age specific mortality, and mean longevity in temperature-dependent cohorts of Culex quinquefasciatus Say (Diptera: Culicidae). Journal of Vector Ecology. 2010;35: 354–362. doi:10.1111/j.1948-7134.2010.00094.x

48. Olayemi IK, Victoria O, Ukubuiwe AC, Jibrin AI. Effects of Temperature Stress on Pre-imaginal Development and Adult Ptero-fitness of the Vector Mosquito, *Culex quinquefasciatus* (Diptera: Culicidae). jmr. 2016 [cited 9 Feb 2024]. doi:10.5376/jmr.2016.06.0014

49. Dodson BL, Kramer LD, Rasgon JL. Effects of larval rearing temperature on immature development and West Nile virus vector competence of Culex tarsalis. Parasites Vectors. 2012;5: 199. doi:10.1186/1756-3305-5-199

50. Shriver D, Bickley W. The effect of temperature on the hatching of eggs of the mosquito Culex pipiens quinquefasciatus say. Mosquito News. 1964;24: 137–40.

51. van der Linde T de K, Hewitt P, Nel A, van der Westhuizen M. Development rates and percentage hatching of culex (Culex) theileri; Theobald (Diptera: culicidae) eggs at various constant temperatures. Journal of the Entomological Society of Southern Africa. 53: 17–26.

52. Nayar JK. Effects of constant and fluctuating temperatures on life span of Aedes taeniorhynchus adults. Journal of Insect Physiology. 1972;18: 1303–1313. doi:10.1016/0022-1910(72)90259-4

53. Oda T, Eshita Y, Uchida K, Mine M, Kurokawa K, Ogawa Y, et al. Reproductive Activity and Survival of *Culex pipiens pallens* and *Culex quinquefasciatus* (Diptera: Culicidae) in Japan at High Temperature. J Med Entomol. 2002;39: 185–190. doi:10.1603/0022-2585-39.1.185

54. Andreadis SS, Dimotsiou OC, Savopoulou-Soultani M. Variation in adult longevity of Culex pipiens f. pipiens, vector of the West Nile Virus. Parasitol Res. 2014;113: 4315–4319. doi:10.1007/s00436-014-4152-x

55. Reisen WK, Milby MM, Presser SB, Hardy JL. Ecology of Mosquitoes and St. Louis Encephalitis Virus in the Los Angeles Basin of California, 1987–1990. Journal of Medical Entomology. 1992;29: 582–598. doi:10.1093/jmedent/29.4.582

56. Moser SK, Barnard M, Frantz RM, Spencer JA, Rodarte KA, Crooker IK, et al. Scoping review of Culex mosquito life history trait heterogeneity in response to temperature. Parasites Vectors. 2023;16: 200. doi:10.1186/s13071-023-05792-3

57. Harbach RE. Culex pipiens: Species Versus Species Complex – Taxonomic History and Perspective. Journal of the American Mosquito Control Association. 2012;28: 10–23. doi:10.2987/8756-971X-28.4.10

58. Haba Y, McBride L. Origin and status of Culex pipiens mosquito ecotypes. Current Biology. 2022;32: R237–R246. doi:10.1016/j.cub.2022.01.062

59. DiSera L, Sjödin H, Rocklöv J, Tozan Y, Súdre B, Zeller H, et al. The Mosquito, the Virus, the Climate: An Unforeseen Réunion in 2018. GeoHealth. 2020;4. doi:10.1029/2020GH000253

60. Tran A, Mangeas M, Demarchi M, Roux E, Degenne P, Haramboure M, et al. Complementarity of empirical and process-based approaches to modelling mosquito population dynamics with Aedes albopictus as an example—Application to the development of an operational mapping tool of vector populations. Touzeau S, editor. PLoS ONE. 2020;15: e0227407. doi:10.1371/journal.pone.0227407

61. Liu-Helmersson J, Brännström Å, Sewe MO, Semenza JC, Rocklöv J. Estimating Past, Present, and Future Trends in the Global Distribution and Abundance of the Arbovirus Vector Aedes aegypti Under Climate Change Scenarios. Front Public Health. 2019;7: 148. doi:10.3389/fpubh.2019.00148

62. Yang HM, Boldrini JL, Fassoni AC, Freitas LFS, Gomez MC, Lima KKB de, et al. Fitting the Incidence Data from the City of Campinas, Brazil, Based on Dengue Transmission Modellings Considering Time-Dependent Entomological Parameters. Paul R, editor. PLoS ONE. 2016;11: e0152186. doi:10.1371/journal.pone.0152186

63. Caldwell JM, LaBeaud AD, Lambin EF, Stewart-Ibarra AM, Ndenga BA, Mutuku FM, et al. Climate predicts geographic and temporal variation in mosquito-borne disease dynamics on two continents. Nat Commun. 2021;12: 1233. doi:10.1038/s41467-021-21496-7

64. Ngonghala CN, Ryan SJ, Tesla B, Demakovsky LR, Mordecai EA, Murdock CC, et al. Effects of changes in temperature on Zika dynamics and control. J R Soc Interface. 2021;18: rsif.2021.0165, 20210165. doi:10.1098/rsif.2021.0165

65. Mordecai EA, Paaijmans KP, Johnson LR, Balzer C, Ben-Horin T, de Moor E, et al. Optimal temperature for malaria transmission is dramatically lower than previously predicted. Thrall P, editor. Ecol Lett. 2013;16: 22–30. doi:10.1111/ele.12015

66. Parham PE, Michael E. Modeling the Effects of Weather and Climate Change on Malaria Transmission. Environmental Health Perspectives. 2010;118: 620–626. doi:10.1289/ehp.0901256

67. Rayah EAE, Groun NAA. *Effect of temperature on hatching eggs and embryonic survival in the mosquito* Culex quinquefasciatus. Entomologia Exp Applicata. 1983;33: 349–351. doi:10.1111/j.1570-7458.1983.tb03281.x

68. Trpiš M, Shemanchuk JA. EFFECT OF CONSTANT TEMPERATURE ON THE LARVAL DEVELOPMENT OF *AEDES VEXANS* (DIPTERA: CULICIDAE). Can Entomol. 1970;102: 1048–1051. doi:10.4039/Ent1021048-8

69. Mpho M, Holloway GJ, Callaghan A. A comparison of the effects of organophosphate insecticide exposure and temperature stress on fluctuating asymmetry and life history traits in Culex quinquefasciatus. Chemosphere. 2001;45: 713–720. doi:10.1016/S0045-6535(01)00140-0

70. McHaffey DG. Photoperiod and Temperature Influences on Diapause in Eggs of the Floodwater Mosquito Aedes Vexans (Meigen) (Diptera: Culicidae)1, 2. Journal of Medical Entomology. 1972;9: 564–571. doi:10.1093/jmedent/9.6.564

71. McHaffey DG, Harwood RF. Photoperiod and Temperature Influences on Diapause in Eggs of the Floodwater Mosquito, Aedes Dorsalis (Meigen) (Diptera: Culicidae)1. Journal of Medical Entomology. 1970;7: 631–644. doi:10.1093/jmedent/7.6.631

72. Paull SH, Horton DE, Ashfaq M, Rastogi D, Kramer LD, Diffenbaugh NS, et al. Drought and immunity determine the intensity of West Nile virus epidemics and climate change impacts. Proc R Soc B. 2017;284: 20162078. doi:10.1098/rspb.2016.2078
